# Nuclear bodies protect phase separated proteins from degradation in stressed proteome

**DOI:** 10.1101/2023.04.19.537522

**Authors:** Kwan Ho Jung, Jiarui Sun, Chia-Heng Hsiung, Xiaojun Lance Lian, Yu Liu, Xin Zhang

## Abstract

RNA-binding proteins (RBPs) containing intrinsically disordered domains undergo liquid-liquid phase separation to form nuclear bodies under stress conditions. This process is also connected to the misfolding and aggregation of RBPs, which are associated with a series of neurodegenerative diseases. However, it remains elusive how folding states of RBPs changes upon the formation and maturation of nuclear bodies. Here, we describe SNAP-tag based imaging methods to visualize the folding states of RBPs in live cells via time-resolved quantitative microscopic analyses of their micropolarity and microviscosity. Using these imaging methods in conjunction with immunofluorescence imaging, we demonstrate that RBPs, represented by TDP-43, initially enters the PML nuclear bodies in its native state upon transient proteostasis stress, albeit it begins to misfolded during prolonged stress. Furthermore, we show that heat shock protein 70 co-enters the PML nuclear bodies to prevent the degradation of TDP-43 from the proteotoxic stress, thus revealing a previously unappreciated protective role of the PML nuclear bodies in the prevention of stress-induced degradation of TDP-43. In summary, our imaging methods described in the manuscript, for the first time, reveal the folding states of RBPs, which were previously challenging to study with conventional methods in nuclear bodies of live cells. This study uncovers the mechanistic correlations between the folding states of a protein and functions of nuclear bodies, in particular PML bodies. We envision that the imaging methods can be generally applied to elucidating the structural aspects of other proteins that exhibit granular structures under biological stimulus.

## Introduction

An emerging aspect in the structural organization of the cell nucleus is that a wide variety of cellular processes are organized by subnuclear compartments, collectively referred to as nuclear bodies.^1^ Unlike other membrane-bound organelles, nuclear bodies are not membrane-bound, but able to remain coherent structures by assembling specific nuclear proteins and/or RNAs to initiate distinct biochemical functions.^2^ The example includes, but not limited to, nucleolus, nuclear speckle, nuclear stress body, paraspeckle, and promyelocytic leukemia (PML) nuclear body.^3, 4^ Numerous studies have revealed that they exhibit dynamic properties and steadily exchange their components with the surrounding nucleoplasm.^2^ Furthermore, there is an emerging body of evidence showing that these structures are formed and regulated by liquid-liquid phase separation (LLPS).^5^

A commonly shared feature of proteins that are associated with nuclear bodies is that they contain low complexity domains, which are intrinsically disordered and prone to misfolding and aggregation under stress in cells.^6–9^ In particular, the cell nucleus is rich with RNA-binding proteins (RBPs) that contain prion-like domains (PLDs), such as TDP-43, FUS, and TAF15.^10–12^ PLDs are low complexity domains enriched in uncharged polar amino acid residues with compositional similarity to yeast prion proteins.^13^ Misfolding and aggregations of such proteins have been linked to several neurodegenerative diseases, including amyotrophic lateral sclerosis (ALS) and frontotemporal lobar degeneration (FTLD), demonstrating the importance of understanding the misfolding and aggregation of RBPs in cells.^14–18^ Nevertheless, despite considerable progress having been made in their purifications and structural characterizations, understanding their physiological and pathological behaviors of RBPs within live intact cells remains a challenge.

Recent studies indicate that several RBPs with PLDs are susceptible to environmental stresses or inhibition of signaling pathways, leading to the formation of nuclear puncta. For instance, TDP-43 has been shown to form granular structures in the nucleus under proteotoxic, oxidative, and heat stresses.^19–22^ Likewise, FUS has been reported to form granular structures in the nucleus upon inhibition of arginine methylation.^23^ While these findings suggest that RBPs are prone to form granular structures in response to stress, it remains unclear whether RBPs enter nuclear bodies in either native states or soluble misfolded oligomer states. This question led us to consider two working models of how RBPs enter nuclear bodies under stress (**Figure 1a**). RBPs may enter nuclear bodies under transient stress either in their native states (Model 1) or in their soluble oligomer states (Model 2). After prolonged stress, the recruited RBPs may misfold and aggregate in nuclear bodies. However, it has been difficult to distinguish these models using conventional methods based on fluorescent protein fusion, because both native states and soluble oligomeric states exhibit diffuse fluorescent signal. Thus, these methods would result in the same signal change from diffuse to punctate in both models. Herein, we set out to test these models by utilizing the AggTag method that is developed in our labrotory.^24, 25^ Combined with multi-mode imaging strategies, this method allows us to visualize the folding states of RBPs and to estimate their micropolarity and microviscosity in live cells. We show that RBPs such as FUS and TAF15 enter the nucleolus without misfolding upon inhibition of methyltransferases. Similarly, upon proteotoxic stress, TDP-43 initially enters the PML nuclear bodies in a native state, although it begins to misfold in the nuclear bodies under prolonged stress. We further find that HSPA1A, a member of the heat shock protein 70 family, is involved in the association of TDP-43 with the PML nuclear bodies in a native state to prevent the degradation of TDP-43 from the proteotoxic stress. Taken together, our findings indicate that the PML nuclear bodies play a previously unknown protective role in the prevention of degradation of proteins against stresses.

**Figure 1.**
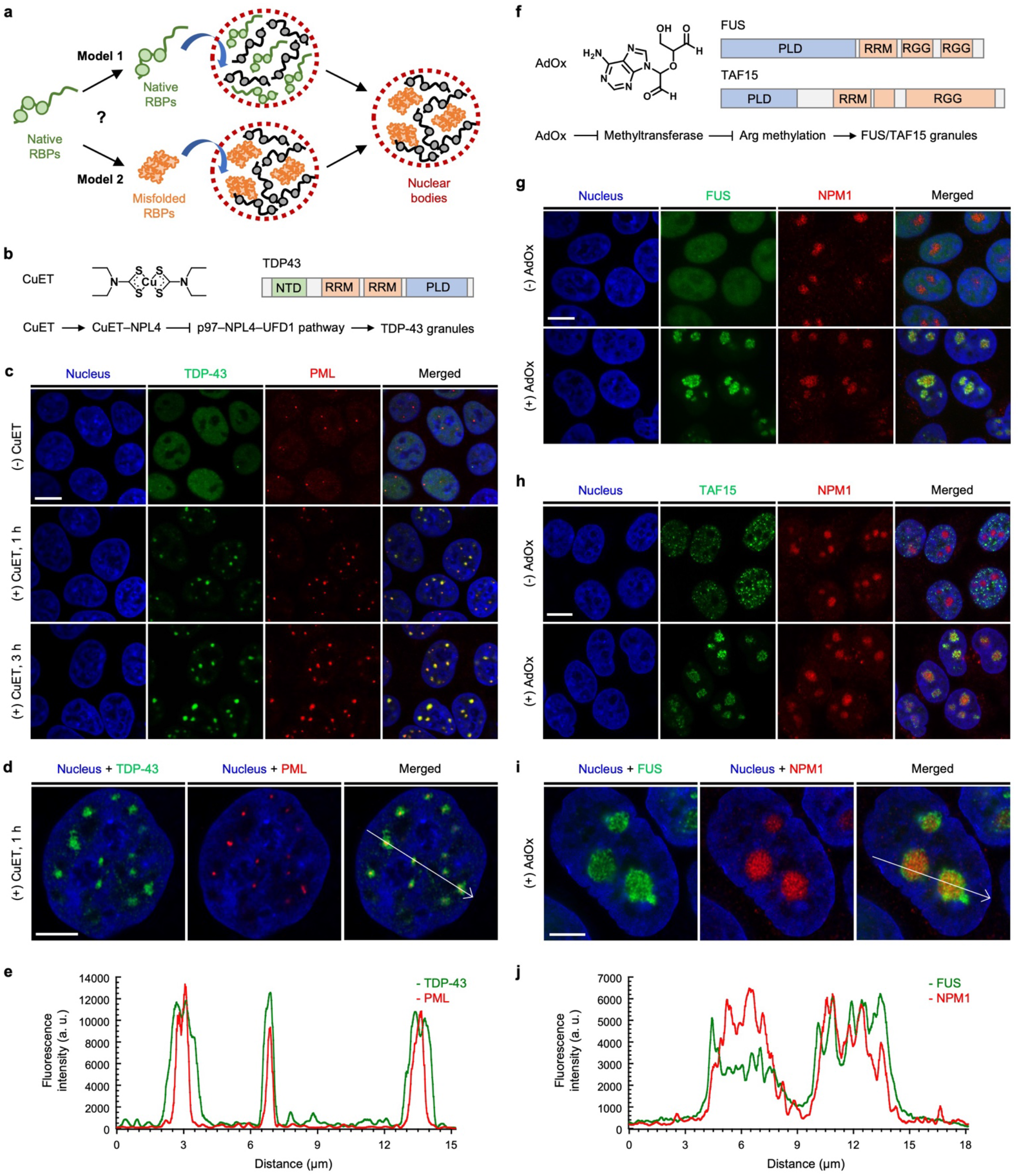
RNA-binding proteins (RBPs) containing prion-like domains (PLDs) enter nuclear bodies when specific signaling pathways are inhibited by chemicals. (**a**) Two working models of how RBPs with PLDs enter nuclear bodies. In model 1, nuclear bodies recruit RBPs in their native states. In model 2, nuclear bodies sequester RBPs in their soluble misfolded oligomer states. (**b**) Structure of CuET, domain architecture of TDP-43, and how CuET treatment induces the formation of the TDP-43 granular structures. (**c**) Immunofluorescence imaging of cells stably expressing SNAPf–TDP-43 (labeled by SNAP-Cell Oregon Green, green) and treated with CuET for 0, 1 or 3 h using anti-PML (red). Scale bar = 10 µm. (**d**), (**e**) Airyscan imaging (d) and intensity profile analysis (e) of cells stably expressing SNAPf–TDP-43 (labeled by SNAP-Cell Oregon Green, green) and treated with CuET for 1 h. PML nuclear bodies were labeled with anti-PML (red) by immunofluorescence. Scale bar = 5 µm. (**f**) Structure of AdOx, domain architectures of FUS and TAF15, and how AdOx treatment induces the formation of the FUS and TAF15 granular structures. (**g**), (**h**) Immunofluorescence imaging of cells stably expressing either FUS–SNAPf (g) or SNAPf–TAF15 (h) (labeled by SNAP-Cell Oregon Green, green) using anti-NPM1 (red) with or without AdOx pretreatment. Scale bar = 10 µm. (**i**), (**j**) Airyscan imaging (i) and intensity profile analysis (j) of cells stably expressing FUS–SNAPf (labeled by SNAP-Cell Oregon Green, green) with AdOx pretreatment. The nucleoli were labeled with anti-NPM1 (red) by immunofluorescence. Scale bar = 5 µm.

## Results

### RBPs with PLDs enter nuclear bodies by inhibition of specific cellular pathways

Previous studies have reported that bis(diethyldithiocarbamate)–copper (CuET) inhibits the p97 segregase pathway by targeting to its adaptor protein NPL4.^22, 26–28^ As the p97 segregase is a central component of the ubiquitin-proteasome system, CuET treatment was shown to induce proteotoxic stress in cells, causing TDP-43 to form granular structures in the nucleus (**Figure 1b**).^22^ In order to establish the identity of such granular structures, we generated a human embryonic kidney (HEK) 293T cell line that can stably express SNAP-tag fused TDP-43 (SNAPf–TDP-43) by doxycycline induction, using the PiggyBac XLone system.^29^ SNAP-tag, a small self-labeling protein tag that covalently and specifically reacts with *O*^6^-benzylguanine (BG) moiety,^30^ was chosen to label RBPs due to its relatively small size and minimal perturbation on phase separation behaviors of RBPs.^31^

We hypothesized that TDP-43 may enter nuclear bodies upon proteotoxic stress induced by CuET. To test this, we performed immunofluorescence imaging experiments using antibodies against the nuclear body markers. After stably expressing SNAPf–TDP-43 by doxycycline treatment and labeling the fusion protein with SNAP-Cell Oregon Green, the cell line was further treated, or not, with CuET to induce the TDP-43 granular structures. Subsequently, the cell line was stained with the antibodies against several different nuclear body markers, including NPM1 (for the nucleolus), SFPQ (for the paraspeckles), HSF1 (for the nuclear stress bodies), PML (for the PML nuclear bodies), or SC35 (for the nuclear speckles) (**Figure S1a**). After screening with several nuclear body markers, we identified that the TDP-43 granular structures were colocalized with PML (**Figure 1c**). Without CuET treatment, TDP-43 remained diffuse in the nucleus while PML formed nuclear foci that were indicative of the PML nuclear bodies. Upon treatment with CuET for 1 or 3 hours, TDP-43 formed the granular structures, and they were colocalized with PML. This result indicates that TDP-43 enters the PML nuclear bodies upon proteotoxic stress induced by CuET. On the other hand, the TDP-43 granular structures were not colocalized with SC35, HSF1, NPM1, or SFPQ, indicating that the recruitment of TDP-43 is specific to the PML nuclear bodies (**Figure S1b-c**). When we used Airyscan microscopy and z-stack analysis to confirm the colocalization, the PML nuclear bodies were found to reside in the core of the TDP-43 granular structures (**Figure 1d-e**, **Figure S2**). In addition, using fluorescence recovery after photobleaching, we found that the initially formed TDP-43 granular structures by CuET treatment showed minimal fluorescence recovery following photobleaching. This suggests that upon incorporation into PML nuclear bodies, TDP-43 become immobile, possibly due to the specific and relatively strong interaction that form the PML nuclear bodies, in comparison to the non-specific, multivalency, and weak interactions used to form liquid droplets (**Figure S3**).^32, 33^

Adenosine dialdehyde (AdOx) has been used to indirectly inhibit a family of S-adenosylmethionine-dependent methyltransferases (**Figure 1f**).^34, 35^ It has been shown that treatment with AdOx causes FUS to be in its hypomethylated states and to form nuclear granular structures.^23^ We then generated a doxycycline-inducible HEK293T cell line stably expressing SNAP-tag fused FUS (FUS–SNAPf), and conducted the same immunofluorescence imaging experiments using the antibodies against the nuclear body markers. After the cell line of FUS–SNAPf was pretreated with AdOx, doxycycline and SNAP-Cell Oregon Green were added to express and label the fusion protein. The cells were then stained with the antibodies against the different nuclear body markers (**Figure S4a**). Among several nuclear body markers, the FUS granular structures were found to be colocalized with NPM1 (**Figure 1g**). While FUS remained diffuse in the nucleus when not pretreated with AdOx, pretreatment with AdOx caused FUS to form the granular structures that were colocalized with NPM1, but not with SC35, HSF1, PML, or SFPQ, indicating that FUS in its hypomethylated state only entered nucleolus (**Figure S4b-c**). The Airyscan and z-stack imaging experiments confirmed that FUS resided within the nucleolus in cells pretreated with AdOx (**Figure S5**). As revealed by fluorescence recovery after photobleaching, FUS and NPM1 exhibited dynamic properties in the nucleolus, suggesting that the phase separation of FUS may mediate the entering of FUS to the nucleolus (**Figure S6**). Given the structural similarity among the RBP,^36, 37^ we then chose TAF15 to investigate the identity of its nuclear body formation (**Figure S7a**). As expected, we found that the TAF15 granular structures were colocalized with NPM1 among several different nuclear body markers in cells that were pretreated with AdOx (**Figures 1h and S7b-c**). This result indicates that, similar to FUS, TAF15 also enters the nucleolus when pretreated with AdOx. This was further confirmed by the Airyscan and z-stack imaging experiments, showing that TAF15 resided inside the nucleolus when cells were pretreated with AdOx (**Figure S8**). Taken together, these results indicate that when specific cellular pathways are inhibited by chemicals, RBPs with PLDs enter nuclear bodies, such as PML nuclear bodies for TDP-43 and the nucleolus for FUS and TAF15.

### Triple-color imaging assay visualizes folding states of TDP-43 entering the PML nuclear bodies during proteotoxic stress

While the conventional methods to visualize RBPs can monitor their localization into the nuclear bodies during cellular stress, they do not reveal whether the nuclear bodies recruit RBPs in native or misfolded oligomeric states because both states can exhibit diffuse characteristics in cells. In this regard, we set out to investigate the folding states of the RBPs when they initially enter the nuclear bodies under stress conditions. To this end, we utilized the previously reported AggTag method to monitor the misfolding and aggregation of a protein of interest (POI) by a turn-on of fluorescence in live cells.^24, 25^ In this method, an environment-sensitive fluorophore is conjugated to the POI via a self-labeling protein tag, and it remains fluorescently quenched when POI is folded. The formation of misfolded oligomers and insoluble aggregates of POI then buries the fluorophore in the aggregates to elicit its turn-on of fluorescence.

To utilize the AggTag method in visualizing the folding states of SNAP-tag fused TDP-43 in live cells, we synthesized three SNAP-tag substrates (**P0**, **P1**, and **P2**) and devised a triple-color imaging assay (**Figure 2a**). **P0** contains 7-amino-4-methylcoumarin whose blue emission is constitutive, reporting RBPs in all the conformational states. **P1** contains a molecular rotor derived from the Kaede chromophore, which is previously reported to activate its red fluorescence in response to soluble misfolded oligomers.^24, 25, 38^ **P2** contains a molecular rotor derived from the green fluorescent chromophore which is previously shown to activate its green fluorescence toward insoluble oligomers.^38^ Thus, in this triple-color imaging assay, we expected that proteins in native states could be visualized by **P0** fluorescence but no **P1** or **P2** fluorescence, whereas soluble oligomers and insoluble aggregates of RBPs could be detected by a sequential appearance of **P1** and **P2** fluorescence, respectively (**Figure 2b**). **P1** and **P2** are spectrally orthogonal, and the viscosity sensitivity values of **P1** and **P2** were determined to be 0.20 and 0.54, which are comparable to those of the previously reported chromophores (**Figure S9**).^38^

**Figure 2.**
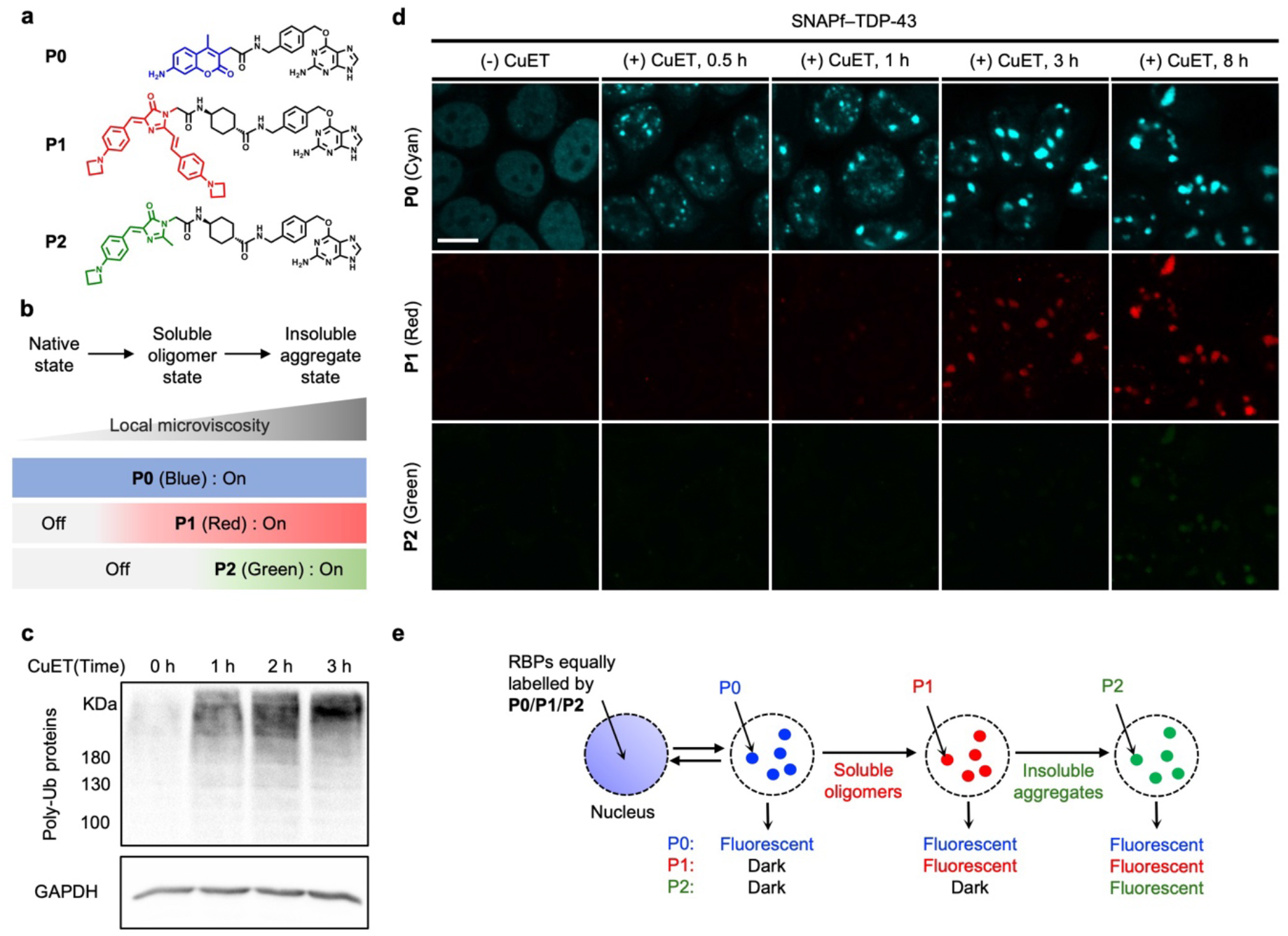
Triple-color imaging assay suggests that TDP-43 remains in its native state when initially entering PML nuclear bodies by transient proteotoxic stress, while prolonged proteotoxic stress causes TDP-43 to misfold in the nuclear bodies. (**a**) Structures of SNAP-tag substrates used in triple-color imaging assay. (**b**) In triple-color imaging assay using AggTag method, proteins in native states, soluble oligomer states, and insoluble aggregate states can be distinguished by the activations of **P1** and **P2** fluorescence. (**c**) CuET treatment leads to proteotoxic stress in cells. Western blot image showing an accumulation of poly-ubiquitylated proteins upon treatment with CuET. (**d**) Triple-color imaging assay results for TDP-43 during proteotoxic stress. Confocal fluorescence images of **P0** (cyan), **P1** (red), and **P2** (green) in cells stably expressing SNAPf–TDP-43 upon treatment with CuET for different time periods. Scale bar = 10 µm. (**e**) Schematic illustration of the triple-color imaging assay result for TDP-43. TDP-43 remains in its native state when initially entering PML nuclear bodies by transient proteotoxic stress, while prolonged proteotoxic stress causes TDP-43 to misfold after TDP-43 enters the nuclear bodies.

As previously reported, treatment with CuET causes proteotoxic stress in cells, as supported by the western blot analysis of the cell lysates treated with CuET using anti-polyubiquitin antibody (linkage-specific K48) (**Figure 2c**).^22^ We then conducted the triple-color imaging assay to test whether TDP-43 enters the PML nuclear bodies in native or misfolded oligomer states upon proteotoxic stress. The SNAPf–TDP-43 stable cell line was treated with doxycycline and the identical concentrations of the triple probes (**P0**, **P1**, and **P2**) for its inducible expression and equal labeling, respectively. After 24 hours, the cells were treated with CuET for different time periods and subsequently visualized by confocal microscopy.

In the absence of CuET, cells exhibited nuclear diffuse fluorescence from **P0**, but no fluorescence from **P1** or **P2**, indicating that TDP-43 remains in its native state in the absence of stress (**Figure 2d**, (-) CuET panel). In cells treated with CuET for 30 minutes to an hour, we observed that the diffuse **P0** fluorescence started to convert its punctate fluorescence, indicating the recruitment of TDP-43 into the nuclear bodies induced by CuET. Interestingly, we did not observe significant increase in **P1** or **P2** fluorescence (**Figure 2d**, (+) CuET, 0.5 h and 1 h panels). These imaging results suggest that, although CuET induces proteotoxic stress that causes protein misfolding, TDP-43 may stay in its native state at least initially while entering into the nuclear bodies. When we treated the cells with CuET for the longer time of 3 or 8 hours, we observed that they exhibited punctate **P1** fluorescence that was colocalized with punctate **P0** fluorescence (**Figure 2d**, (+) CuET, 3 h and 8 h panels). These observations suggest that the recruited TDP-43 may undergo misfolding within the nuclear bodies under persistent stress. The overall triple-color imaging assay results for TDP-43 led us to suggest that TDP-43 may stay in its native state during entering the PML nuclear bodies by transient stress for 30 minutes to an hour, albeit persistent stress longer than 3 hours may cause TDP-43 to misfold within the nuclear bodies (**Figure 2e**).

### FLIM imaging assays estimate micropolarity and microviscosity of TDP-43 entering PML nuclear bodies during proteotoxic stress

The triple color imaging assay is based on measuring the fluorescence intensity of molecular rotors by conventional confocal microscopy. However, relying on fluorescence intensity changes may cause biased results, as the fluorescence intensity can also be affected by several parameters, including the local concentration changes of fluorescent species, photobleaching, and laser intensity.^39^ Thus, we turned to using fluorescence lifetime imaging microscopy (FLIM), an imaging technique to measure the fluorescence lifetime of individual fluorophores at each pixel of an image.^40^ As the fluorescence lifetime is dependent on the environment of a fluorophore but not dependent on external factors, we reasoned that FLIM would be more robust and quantitative than the intensity-based method to study folding states of RBPs in live cells. In particular, by utilizing an environment-sensitive fluorophore that is either polarity- or viscosity-sensitive, we envisioned to establish a FLIM imaging assay that is able to estimate the local micropolarity or microviscosity of RBPs in live cells, respectively. This would also allow us to confirm the results from the triple-color imaging assay, as soluble misfolded oligomers have lower micropolarity and higher microviscosity than proteins in their native states.

To design the FLIM imaging assay for estimating the local micropolarity of RBPs, we synthesized a new SNAP-tag substrate, **P3**, which contains 7-dimethylamino-4-sulfamonyl-2,1,3-benzoxadiazole (SBD) fluorophore (**Figure 3a**). The SBD is a solvatochromic fluorophore whose lifetime is sensitive to the polarity of the local microenvironment.^41^ To correlate its fluorescence lifetime with dielectric constant, the fluorescence lifetimes of **P3** were measured in several different organic solvents with various dielectric constants by FLIM (**Figure S10**). As the dielectric constant of the solvent tested decreased from 33 (ethylene glycol) to 12.5 (tert-butanol), the fluorescence lifetime of **P3** in solutions linearly increased from 2.72 ns to 7.85 ns. This result confirmed its polarity-dependent fluorescence lifetime and provided a correlation equation, which enabled us to estimate the local micropolarity of RBPs in live cells (**Figure 3b**).

**Figure 3.**
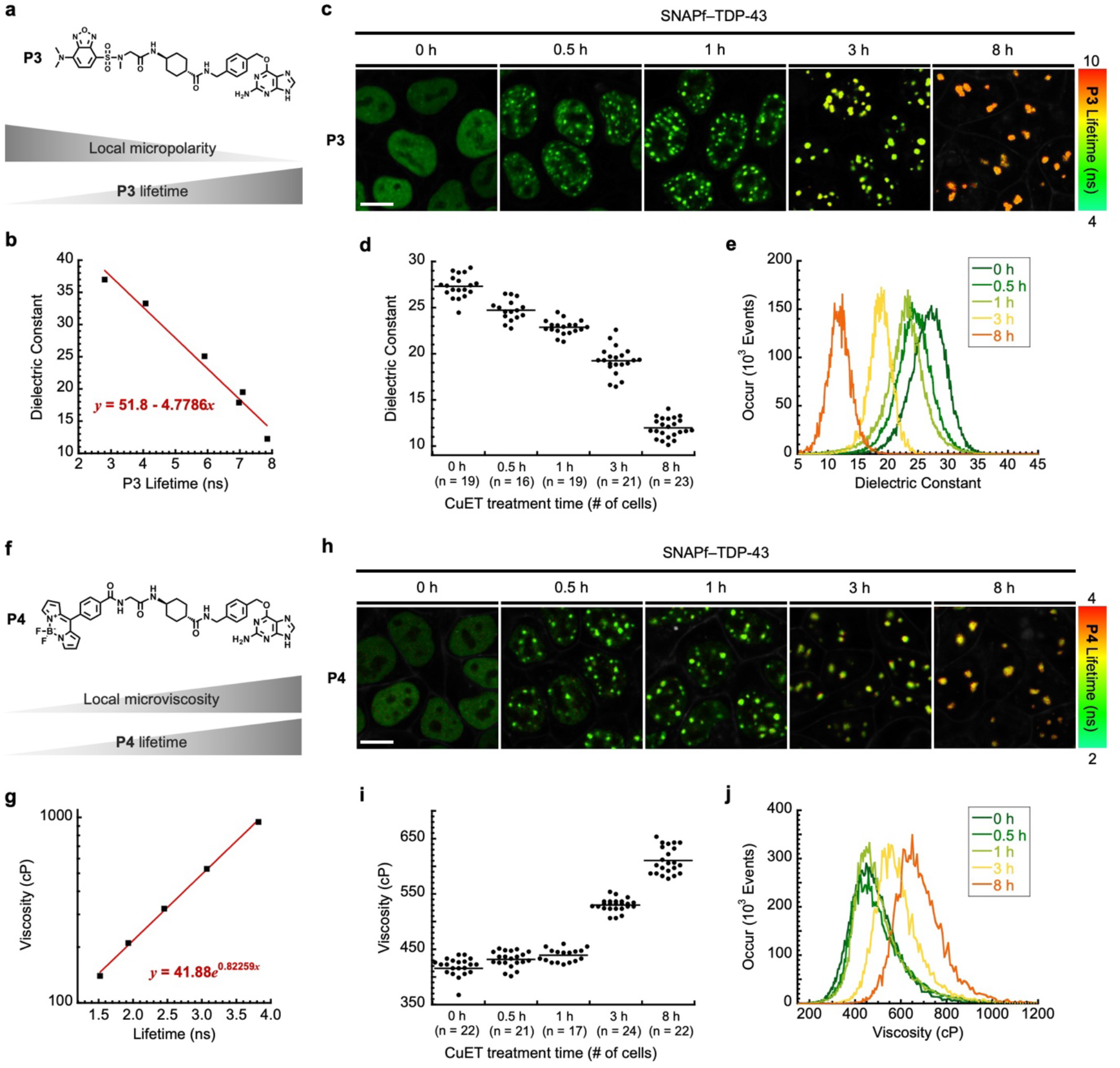
Fluorescence lifetime imaging assays using P3 and P4 estimate local micropolarity and microviscosity changes of TDP-43 entering PML nuclear bodies during proteotoxic stress by CuET. (**a**) Structure of **P3** and its polarity-dependent lifetime. As the local micropolarity decreases, the lifetime of **P3** increases. (**b**) Linear plot of dielectric constant as a function of **P3** lifetime. (**c**) Representative fluorescence lifetime images of **P3** in cells stably expressing SNAPf–TDP-43 at different time points of CuET treatment. Scale bar = 10 µm. (**d**) Dot plot of dielectric constant values of TDP-43 per cell at different time points of CuET treatment. (**e**) Combined histograms of dielectric constant values of TDP-43 at different time points of CuET treatment. (**f**) Structure of **P4** and its viscosity-dependent lifetime. As the local microviscosity increases, the lifetime of **P4** increases. (**g**) Linear plot of viscosity as a function of **P4** lifetime. Y-axis is shown in log scale. (**h**) Representative fluorescence lifetime images of **P4** in cells stably expressing SNAPf–TDP-43 at different time points of CuET treatment. Scale bar = 10 µm. (**i**) Dot plot of viscosity values of TDP-43 per cell at different time points of CuET treatment. (**j**) Combined histograms of viscosity values of TDP-43 at different time points of CuET treatment.

We then performed the FLIM imaging assay with **P3** to estimate the local micropolarity of TDP-43 during proteotoxic stress by CuET. To this end, the stable cell line of SNAPf–TDP-43 was treated with doxycycline and **P3** for the inducible expression and labeling of the fusion protein by the probe. After treating with CuET for different time periods, we subsequently conducted the FLIM imaging experiments under live cell settings (**Figure S11a-S15a**). A prediction from this assay was that if TDP-43 remains in their native state upon stress, the micropolarity of TDP-43 would not significantly change before and after the stress. On the other hand, its micropolarity would start to decrease considerably when TDP-43 becomes misfolded due to the exposure of hydrophobic residues during misfolding. In cells that were not treated with CuET, the average fluorescence lifetime of **P3** was determined to be 5.13 ± 0.26 ns (n = 19 cells, **Figure S11**), which corresponds to the dielectric constant value of 27.3 ± 1.2, consistent to the notion that the surface of a folded globular protein exhibits dielectric constant of ∼30 (**Figure 3c-e**). Upon treatment with CuET for 30 minutes and an hour, its average lifetimes were calculated to be 5.67 ± 0.24 ns (n = 16 cells, **Figure S12**) and 6.06 ± 0.16 ns (n = 19 cells, **Figure S13**), corresponding to the dielectric constants of 24.7 ± 1.1 and 22.9 ± 0.8, respectively (**Figure 3c-e**). These results are compatible with our triple-color imaging assay suggesting that TDP-43 may remain in its native state by the transient stress of CuET, as the micropolarity of TDP-43 was not significantly changed at these time points. On the other hand, when the cells were treated with CuET for the longer duration of 3 and 8 hours, the lifetime of **P3** further increased to 6.81 ± 0.32 ns (n = 21 cells, **Figure S14**) and 8.33 ± 0.22 ns (n = 23 cells, **Figure S15**), which were converted into the dielectric constants of 19.2 ± 1.5 and 12.0 ± 1.0, respectively (**Figure 3c-e**). While these lifetime values were still lower than the lifetime value (10.7 ns) obtained from the insoluble aggregates of purified **P3**•SNAPf–TDP-43 conjugates (**Figure S16**), these results indicate that as the duration of CuET treatment increases, the local micropolarity of TDP-43 decreases, and agrees with the triple color imaging assay data suggesting that TDP-43 may begin to misfold upon prolonged treatment with CuET.

To enable the FLIM imaging assay to estimate the local microviscosity of RBPs, we also synthesized a SNAP-tag substrate, **P4**, that contains phenyl-substituted boron-dipyrromethene (BODIPY) molecular rotor whose lifetime is known to be dependent on the local microviscosity (**Figure 3f**).^42–46^ To generate a calibration curve between its lifetime and viscosity, the lifetimes of **P4** were measured in binary mixtures of glycerol and ethylene glycol at different volume fractions whose viscosity values are previously reported.^47^ The lifetime of **P4** increased from 1.52 ns to 3.82 ns when the glycerol contents increased from 60% (corresponding to the viscosity of 140 cP) to 100 % (corresponding to the viscosity of 947 cP) (**Figure S17**), confirming its viscosity-dependent lifetime. By plotting and fitting the data, a correlation was derived to estimate the local microviscosity of RBPs in live cells (**Figure 3g**).

We then carried out the FLIM imaging assay using **P4** to predict the local microviscosity of TDP-43 when the cells were under proteotoxic stress. To conduct the imaging assay, SNAPf–TDP-43 was stably expressed in the presence of **P4** for its labeling, and then the cell line was treated with CuET for different time periods (**Figure S18a-S22a**). Without CuET treatment, the average lifetime of **P4** was calculated to be 2.79 ± 0.06 ns (n = 22 cells, **Figure S18**), which was equivalent to the viscosity of 415 ± 22 cP (**Figure 3h-j**). In cells that were treated with CuET for 30 minutes and an hour, its average lifetimes were determined to be 2.84 ± 0.04 ns (n = 21 cells, **Figure S19**) and 2.86 ± 0.03 ns (n = 17 cells, **Figure S20**), corresponding to the viscosity values of 431 ± 15 cP and 439 ± 12 cP, respectively (**Figure 3h-j**). Because the microviscosity of TDP-43 was not considerably changed before and after CuET treatment to an hour, these results are consistent with our triple-color imaging assay suggesting that TDP-43 may remain in its native state by the transient stress of CuET. On the other hand, when the cell line was further treated with CuET for the longer duration of 3 and 8 hours, the average lifetimes of **P4** increased to 3.08 ± 0.03 ns (n = 24 cells, **Figure S21**) and 3.26 ± 0.05 cP (n = 22 cells, **Figure S22**), respectively. These lifetime values were converted into the viscosity values of 530 ± 11 cP and 610 ± 24 cP, respectively (**Figure 3h-j**), and they were still lower than the lifetime value (3.5 ns) obtained from the insoluble aggregates of purified **P4**•SNAPf–TDP-43 conjugates (**Figure S23**). Because misfolded proteins have higher microviscosity than proteins in their native states,^38^ this suggests that TDP-43 may become misfolded by the prolonged stress of CuET. Taken together, the overall results from the triple-color and FLIM imaging assays for TDP-43 suggest that TDP-43 may remain in its native state when initially entering PML nuclear bodies by transient stress, while persistent stress may lead to misfolding of TDP-43 in the nuclear bodies.

### FUS and TAF15 enter the nucleolus in their native states upon inhibition of methyltransferases

The earlier triple-color and FLIM imaging assays with the SNAPf–TDP-43 stable cell line suggest that TDP-43 may stay in its native state when initially entering the nuclear bodies. We then sought to examine whether this observation is generalizable to other RBPs entering nuclear bodies by asking what the folding states of FUS and TAF15 are when they enter the nucleolus. To this end, we used the stable cell line of SNAP-tag fused FUS or TAF15 to conduct the triple-color and FLIM imaging assays with or without AdOx pretreatment (experimental procedures depicted in **Figure S24a-S27a**). In the absence of AdOx, the cell line stably expressing FUS–SNAPf exhibited diffuse **P0** fluorescence but no **P1** or **P2** fluorescence, indicating that FUS remains in its native state outside the nucleolus (**Figure 4a**, left panel). When AdOx was pretreated, the cell line stably expressing FUS– SNAPf displayed punctate **P0** fluorescence in the nucleus but there was no **P1** or **P2** fluorescence observed (**Figure 4a**, right panel). The observed lack of **P1** fluorescence indicates that FUS may enter the nucleolus in its native state when the cells are pretreated with AdOx. This was also compatible with our result from the fluorescence recovery after photobleaching experiment demonstrating that FUS exhibits dynamic properties in the nucleolus (**Figure S6**).

**Figure 4.**
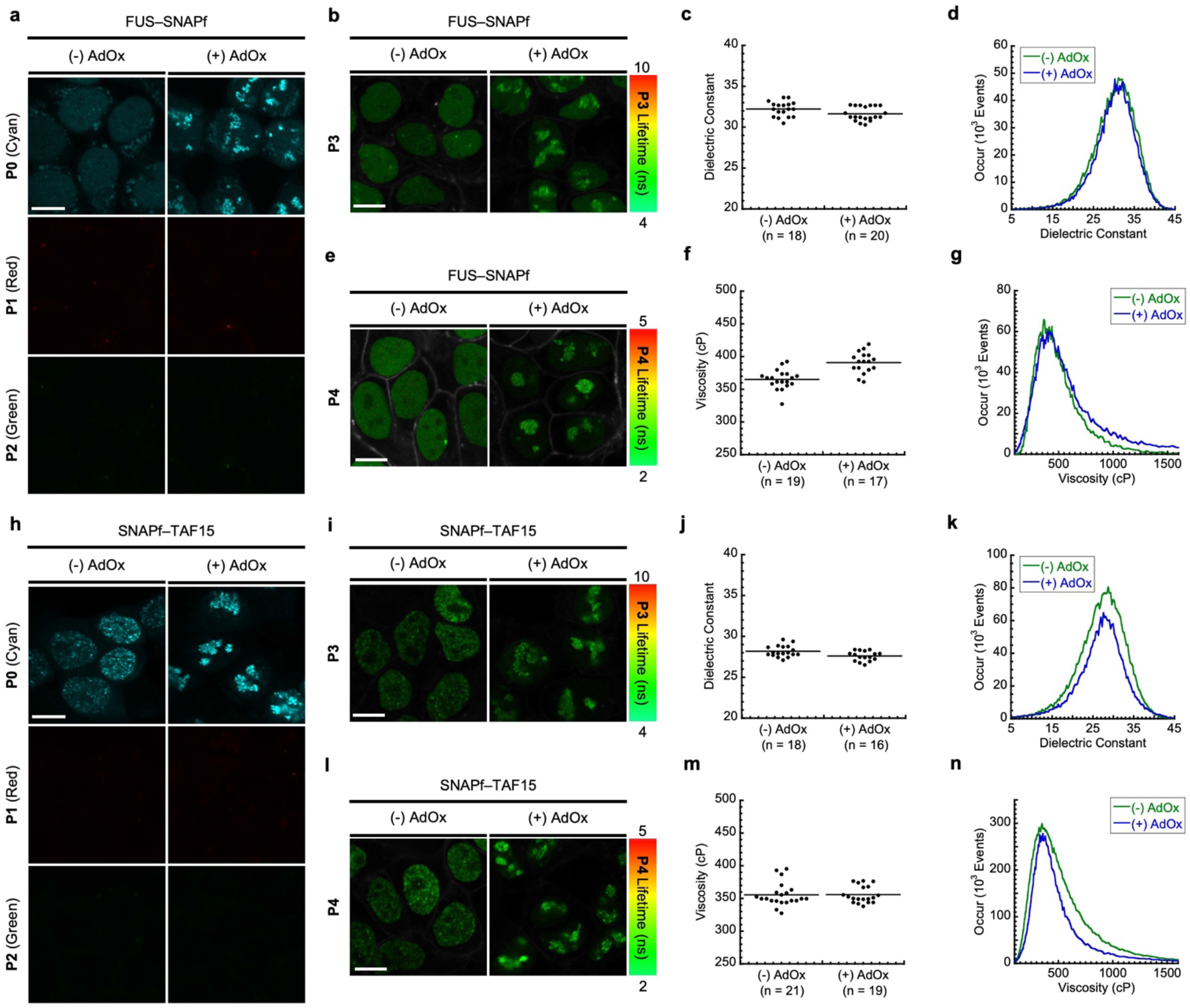
Triple-color and fluorescence lifetime imaging assays suggest that FUS and TAF15 enter the nucleolus in their native states upon methyltransferase inhibition by AdOx. (**a**) Confocal fluorescence images of **P0** (cyan), **P1** (red), and **P2** (green) in cells stably expressing FUS–SNAPf with or without AdOx pretreatment. Scale bar = 10 µm. (**b**) Representative fluorescence lifetime images of **P3** in cells stably expressing FUS–SNAPf with or without AdOx pretreatment. Scale bar = 10 µm. (**c**) Dot plot of dielectric constant values of FUS per cell with or without AdOx pretreatment. (**d**) Combined histograms of dielectric constant values of FUS with or without AdOx pretreatment. (**e**) Representative fluorescence lifetime images of **P4** in cells stably expressing FUS–SNAPf with or without AdOx pretreatment. Scale bar = 10 µm. (**f**) Dot plot of viscosity values of FUS per cell with or without AdOx pretreatment. (**g**) Combined histograms of viscosity values of FUS with or without AdOx pretreatment. (**h**) Confocal fluorescence images of **P0** (cyan), **P1** (red), and **P2** (green) in cells stably expressing SNAPf–TAF15 with or without AdOx pretreatment. Scale bar = 10 µm. (**i**) Representative fluorescence lifetime images of **P3** in cells stably expressing SNAPf–TAF15 with or without AdOx pretreatment. Scale bar = 10 µm. (**j**) Dot plot of dielectric constant values of TAF15 per cell with or without AdOx pretreatment. (**k**) Combined histograms of dielectric constant values of TAF15 with or without AdOx pretreatment. (**l**) Representative fluorescence lifetime images of **P4** in cells stably expressing SNAPf–TAF15 with or without AdOx pretreatment. Scale bar = 10 µm. (**m**) Dot plot of viscosity values of TAF15 per cell with or without AdOx pretreatment. (**n**) Combined histograms of viscosity values of TAF15 with or without AdOx pretreatment.

In the FUS–SNAPf cell line labeled by **P3**, its fluorescence lifetimes were 4.10 ± 0.19 ns (n = 18 cells, **Figure S24**) without AdOx pretreatment, and 4.22 ± 0.18 ns (n = 20 cells, **Figure S25**) with AdOx pretreatment. These corresponded to the dielectric constant values of 32.2 ± 0.9 without AdOx pretreatment and 31.6 ± 0.8 with AdOx pretreatment, indicating that there was no significant difference in the dielectric constant of FUS between whether or not it entered the nucleolus (**Figure 4b-d**). Furthermore, when **P4** was used for the FLIM imaging assay, its average lifetime values were calculated to be 2.63 ± 0.05 ns (n = 19 cells, **Figure S26**) without AdOx pretreatment, and 2.71 ± 0.05 ns (n = 17 cells, **Figure S27**) with AdOx pretreatment. These lifetime values were equivalent to the viscosity values of 365 ± 15 cP without AdOx pretreatment and 391 ± 16 cP with AdOx pretreatment (**Figure 4e-g**). Consistent with the result using **P3**, the microviscosity of FUS was not significantly different between whether or not it entered the nucleolus. Together with data from the triple color imaging assay, these FLIM imaging assay results support the notion that FUS enters the nucleolus it its native state.

We then conducted the same imaging assays for TAF15 to examine what the folding state of TAF15 is when entering the nucleolus. In cells stably expressing SNAPf–TAF15 that were labeled by the triple probes (**P0**, **P1**, and **P2**), no significant fluorescence from either **P1** or **P2** was observed while **P0** was used to visualize TAF15 both in the presence and absence of AdOx (**Figure 4h**). This result indicates that TAF15 may stay in its native state when entering the nucleolus, as adjudged by the lack of **P1** and **P2** fluorescence. In addition, the average fluorescence lifetimes of **P3** in the cell line stably expressing SNAPf–TAF15 were determined to be 4.94 ± 0.14 ns (n = 18 cells, **Figure S28**) without AdOx pretreatment, and 5.06 ± 0.12 ns (n = 16 cells, **Figure S29**) with AdOx pretreatment, corresponding the dielectric constant values of 28.2 ± 0.7 and 27.6 ± 0.6, respectively (**Figure 4i-k**). The average lifetimes of **P4** in the cell line stably expressing SNAPf–TAF15 were 2.60 ± 0.06 ns (n = 21 cells, **Figure S30**) without AdOx pretreatment, and 2.60 ± 0.04 ns (n = 19 cells, **Figure S31**) with AdOx pretreatment, corresponding the viscosity values of 356 ± 18 cP and 356 ± 13 cP, respectively (**Figure 4l-n**). As there were no significant changes in the micropolarity and microviscosity of TAF15 upon pretreatment with AdOx, these results support our interpretation that TAF15 may enter the nucleolus in its native state.

### Molecular chaperons mediate entering of TDP-43 into nuclear bodies to prevent their degradation under stress

The results from the imaging assays raised a question of how misfolding-prone TDP-43 remains its native state when entering the nuclear bodies. We postulated that molecular chaperons would be involved in keeping TDP-43 in its native state while entering the nuclear bodies under stress, as they are known to stabilize protein from misfolding and aggregation and thus to protect the proteome against stress. In support of this view, previous proteomics studies have shown that TDP-43 is constitutively bound to heat shock protein 70 (HSP70) and 40 (DNAJ) chaperon families.^48, 49^ Furthermore, recent studies have reported that certain members of heat shock protein 70 (HSP70), including HSPA1A, were found together with TDP-43 within the nuclear bodies induced by heat stress,^21^ mutations on the RNA-binding domains of TDP-43,^50^ or by oxidative stress.^51^

In this regard, we conducted an immunofluorescence analysis using an anti-HSPA1A antibody in the stable cells expressing SNAPf–TDP-43 upon treatment with CuET to ask whether the TDP-43 granular structures contain endogenous HSPA1A. After the cell line was treated with doxycycline and SNAP-Cell Oregon Green to induce its expression and labeling, it was further treated, or not, with CuET, and then stained with the anti-HSPA1A antibody. Without CuET treatment, both TDP-43 and HSPA1A remained diffuse. However, upon treatment with CuET for 1 or 3 hours, the granular structures of TDP-43 were well colocalized with those of HSPA1A that were formed (**Figure 5a**). We further confirmed the colocalization by co-expressing EBFP– HSPA1A in the cell line expressing SNAPf–TDP-43 that was labeled by SNAP-Cell Oregon Green. Using Airyscan and intensity profile analysis, it was found that the granular signals of TDP-43 were well-overlapped with those of HSPA1A, while the PML nuclear bodies still resided in the core of the TDP-43 granular structures (**Figure 5b-c**). In addition, members of DNAJA family, including DNAJA1 and DNAJA2, were found to colocalize with the TDP-43 granular structures formed by the proteotoxic stress when co-expressed (**Figure 5d-e**). These observations indicate that the proteotoxic stress by CuET leads to the co-recruitment of TDP-43 and heat shock proteins into the PML nuclear bodies, and further suggest that heat shock proteins may keep TDP-43 in its native state when entering the nuclear bodies during the stress.

**Figure 5.**
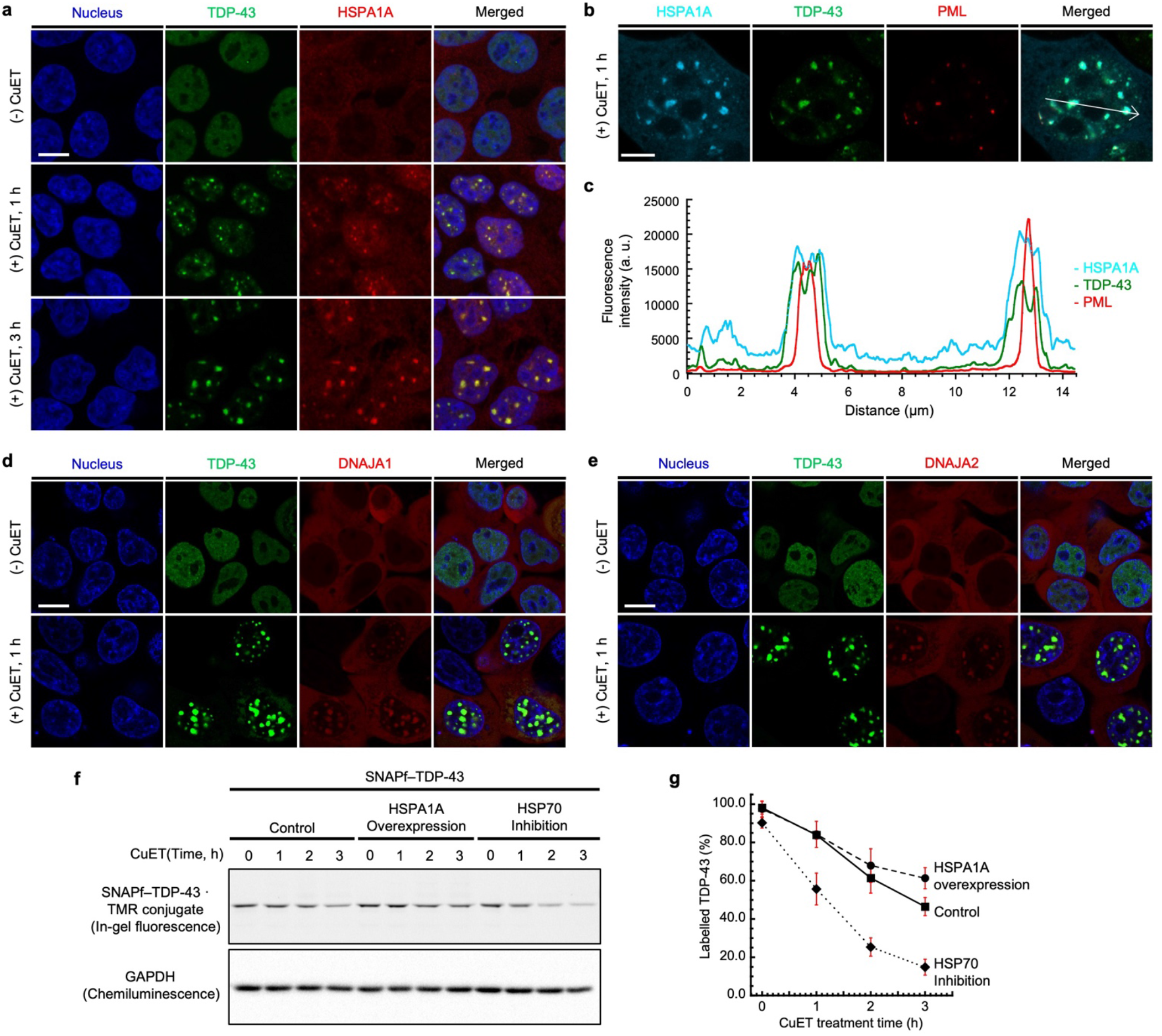
Heat shock proteins mediate the localization of TDP-43 into PML nuclear bodies to prevent its degradation under proteotoxic stress. (**a**) TDP-43 enters PML nuclear bodies together with HSPA1A upon treatment with CuET. Immunofluorescence imaging of cells stably expressing SNAPf–TDP-43 (labeled by SNAP-Cell Oregon Green, green) and treated with CuET for 0, 1 or 3 h using anti-HSPA1A (red). Scale bar = 10 µm. (**b**), (**c**) Airyscan imaging (b) and intensity profile analysis (c) of cells stably expressing SNAPf–TDP-43 (labeled by SNAP-Cell Oregon Green, green) and treated with CuET for 1 h. EBFP–HSPA1A (cyan) was co-expressed and PML nuclear bodies were labeled with anti-PML (red) by immunofluorescence. Scale bar = 5 µm. (**d**) Confocal fluorescence images of cells co-expressing EGFP–TDP-43 (green) and SNAPf–DNAJA1 (labeled by SNAP-Cell TMR Star, red) before and after treatment with CuET. (**e**) Confocal fluorescence images of cells co-expressing EGFP–TDP-43 (green) and SNAPf–DNAJA2 (labeled by SNAP-Cell TMR Star, red) before and after treatment with CuET. (**f**), (**g**) Pulse-chase experiments show that under proteotoxic stress, TDP-43 degrades slower when HSPA1A is overexpressed, and degrades faster when a family of HSP70 is inhibited. (f) In-gel fluorescence image of lysates from cells stably expressing SNAPf–TDP-43 (labeled by SNAP-Cell TMR Star) upon treatment with CuET for different time periods. Left: control, middle: HSPA1A overexpression, right: HSP70 family inhibition. (g) Quantification of fluorescent band intensity from the pulse-chase experiment.

Given the known protective role of HSPA1A,^52^ we further hypothesized that HSPA1A would prevent the degradation of TDP-43 that was caused by the proteotoxic stress. To test this hypothesis, we conducted pulse-chase experiments with the stable cells expressing SNAPf–TDP-43 to measure the half-lives of TDP-43 under different conditions. To this end, we utilized SNAP-Cell block, which is the blocking agent for SNAP-tag to irreversibly inactivate its labeling. After stably expressing SNAPf–TDP-43 and labeling with SNAP-Cell TMR star for 24 hours (pulse), we treated SNAP-Cell block for an hour to block the further labeling of newly synthesized fusion proteins before treating with CuET for different time periods (chase). The level of the fusion protein was then analyzed at each time point of CuET treatment by in-gel fluorescence scanning.

The TMR-labeled SNAPf–TDP-43 was degraded upon treatment with CuET, and its level decreased to approximately 50% in 3 hours after CuET treatment (**Figure 5f-g**). Using this as a control, we conducted the same pulse-chase experiment but overexpressed HSPA1A at the same time while expressing SNAPf–TDP-43, to ask whether the overexpression of HSPA1A affects the half-life of TDP-43 during the proteotoxic stress. When HSPA1A was overexpressed, the level of the TMR-labeled SNAPf–TDP-43 declined to about 60% in 3 hours after CuET treatment (**Figure 5f-g**). The slightly slower degradation of TDP-43 by the HSPA1A overexpression than in control indicates that the HSPA1A expression level affects the half-life of TDP-43 under the proteotoxic stress.

We further examined the half-life of TDP-43 when HSP70 was inhibited during the proteotoxic stress. To this end, we performed the same pulse-chase experiment but pretreating with VER155008, an ATPase inhibitor for the HSP70 family,^53^ at 2 hours before blocking the SNAP-tag labeling. To our surprise, the level of the labeled TDP-43 significantly declined after treatment with CuET in the cells pretreated with VER155008, leading to approximately 20% of labeled TDP-43 analyzed in 3 hours after treatment with CuET (**Figure 5f-g**). The much faster degradation of TDP-43 under the proteotoxic stress supports the hypothesis that HSPA1A may prevent the TDP-43 degradation during the proteotoxic stress. Overall, the results indicate that the recruitment of TDP-43 and HSPA1A to the nuclear bodies may play a role in protecting TDP-43 from its degradation by the proteotoxic stress.

## Discussion

In this work, we aimed to answer the questions of what the folding states of the RBPs are when they associate with nuclear bodies in mammalian cells under stress conditions, and what function the nuclear bodies play in the maintenance of these proteins. Based on what is known in literature, we first devised the two working models of how RBPs enters the nuclear bodies under stress (**Figure 1a**). In the first model, RBPs may be initially recruited into the nuclear bodies in their native states by transient stress, and then they may be converted into their soluble oligomers under persistent stress. This model was supported by studies showing that the initially-formed, liquid-like compartments of RBPs were converted to their solid-like compartments that are reminiscent of protein aggregates in vitro or under persistent cellular stress.^54–57^ In the second model, RBPs may be initially sequestered into the nuclear bodies in their soluble oligomer states upon transient stress via multi-step protein aggregation process. This was supported by studies showing that the RBPs with PLDs were susceptible to misfolding and aggregation by cellular stresses,^13^ and the solid-like, stable core assemblies were observed at the early stage of the stress granule formation.^58, 59^

However, with the currently available methods, it has been extremely challenging to address the question regarding the folding states of the proteins in live cells. Conventional techniques to study a protein associated with the nuclear bodies of live cells typically involve the genetic fusion of the protein to a fluorescent protein.^60–62^ While this method can monitor accumulation of the proteins to the nuclear bodies in live cells, it is unclear whether the proteins remain in their native states or soluble misfolded oligomer states in the nuclear bodies.^63^ To overcome the limitation, we have established new imaging assays to visualize folding states of RBPs live cells with confocal microscopy and to estimate their micropolarity and microviscosity with FLIM through the synthesis of a series of SNAP-tag substrates. We have demonstrated that our imaging methods can provide efficient means to test the models in live cells. Changes in the protein folding states from native to soluble oligomer states were accompanied with the corresponding changes of fluorescence intensity or lifetime of the probes conjugated to the proteins in the nuclear bodies of live cells. In particular, the approach using FLIM provided the additional advantage of not affecting external factors, such as the local concentration of a fluorophore or laser power.^39^

Using such imaging techniques, we have shown that our results were most compatible with the first model. Our results in the triple-color imaging assay with the SNAPf–TDP-43 stable cell line showed that neither **P1** nor **P2** did not turn on their fluorescence during which TDP-43 enters the PML nuclear bodies initially (at 0.5 – 1 h after CuET treatment), and **P1** exhibited an increase in its fluorescence intensity only after prolonged treatment with CuET longer than 3 hours (**Figure 2**). In agreement with these results, our FLIM imaging assays with **P3** and **P4** showed that the micropolarity and microviscosity of TDP-43 did not significantly alter at the early time points of CuET treatment, while the prolonged treatment with CuET led to considerable changes in its micropolarity and microviscosity (**Figure 3**). We further showed that the heat shock proteins were involved in the recruitment of TDP-43 to the PML nuclear bodies to prevent the degradation of TDP-43 by the proteotoxic stress, suggesting the previously unknown protective role of the nuclear bodies (**Figure 5**). To explain the combined results, we propose that TDP-43 enters the PML nuclear bodies initially in a native state upon proteotoxic stress. Prolonged stress may lead to the misfolding of TDP-43 in the nuclear bodies, possibly because it could exhaust the capacity of heat shock proteins to protect TDP-43 from the stress (**Figure 6a**). In the case of FUS and TAF15, the triple color imaging assay revealed that there was no appreciable turning on of fluorescence from **P1** and **P2** before and after they entered the nucleolus by AdOx (**Figure 4**). Using the FLIM assay, as assessed by the lifetimes of **P3** and **P4**, we found that the micropolarity and microviscosity of these proteins did not significantly change before and after incorporation to the nucleolus (**Figure 4**). We interpret these results to propose that FUS and TAF15 enter the nucleolus in their native states upon inhibition of methyltransferases (**Figure 6b**).

**Figure 6.**
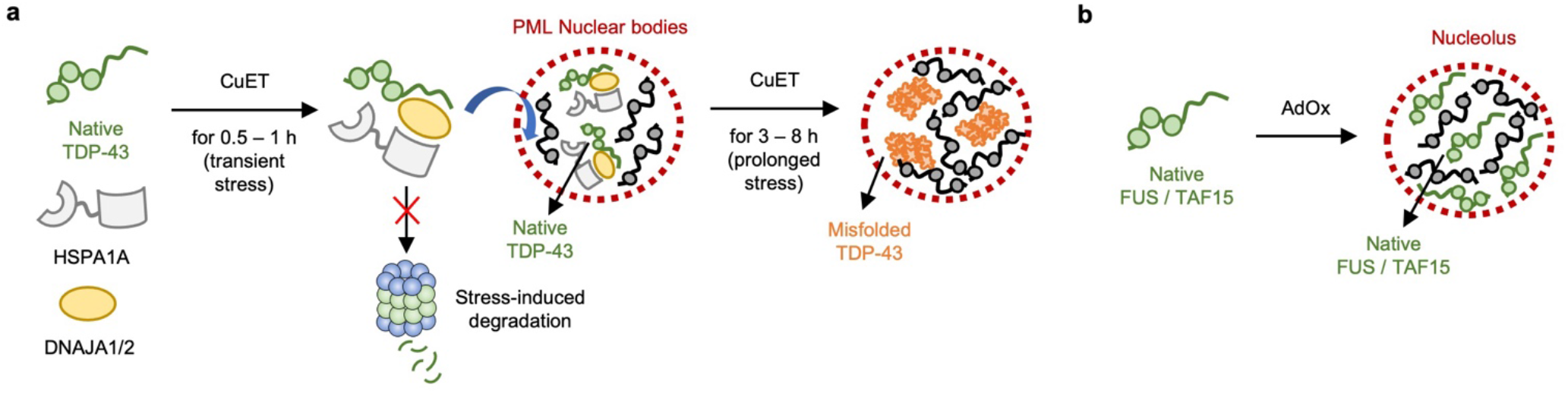
Proposed models for how RBPs behave under stress. (**a**) Upon proteotoxic stress, TDP-43 may enter the PML nuclear bodies in its native state together with heat shock proteins, including HSPA1A and DNAJA1/2. As such, the PML nuclear bodies may play a protective role in preventing the degradation of TDP-43 under transient stress. However, the protective effect may have limited capacity that can be overwhelmed by prolonged stress, resulting in misfolding of TDP-43 within the PML nuclear bodies. (**b**) Upon inhibition of arginine methylation by AdOx, FUS and TAF15 may enter the nucleolus in their native states.

Our imaging methods described in this study revealed, for the first time, the folding states of RBPs in the nuclear bodies in live cells, and they can potentially contribute to this field of research. It has been shown that the Hsp40 proteins phase separate into nuclear bodies to chaperones proteins.^64^ In particular, DNAJB1 accumulates in the ubiquitin-rich nuclear bodies, and DNAJA1 is found in nucleoli. We envision that the presented imaging methods can further be extended to studying how chaperones affect the folding states of other RBPs with PLDs and their diseases-liked mutants. Other than TDP-43, FUS, and TAF15, more than 70 human RBPs have been identified to contain PLDs.^13^ Furthermore, their ALS-linked mutations have been shown to significantly affect the biochemical properties of the full-length proteins, such as increased misfolding and aggregation propensity.^65^ Thus, we expect that there will be ALS-associated mutants of the RBPs that can follow the second model; they enter the nuclear bodies in soluble oligomer states when cells are stressed. In this case, the misfolded RBPs could diffuse in nucleus at the beginning of stress before the incorporation into the nuclear bodies, which could be visualized and measured by our imaging assays. Collectively, this work provides new insight into the structural aspects of RBPs that were previously challenging to obtain with conventional methods in live cells.

In conclusion, we design and apply SNAP-tag based imaging assays to probe the folding states of RBPs entering the nuclear bodies in live cells. The presented imaging methods were applied for three different RBPs, TDP-43, FUS and TAF15. Our imaging assay results for TDP-43 indicate that transient proteotoxic stress up to an hour causes TDP-43 to enter the PML nuclear bodies in its native state, while prolonged proteotoxic stress longer than 3 hours leads to its misfolding within the PML nuclear bodies. The imaging assay results for FUS and TAF15 show that they remain in their native states in the nucleolus upon inhibition of methyltransferases. In addition, we found that the TDP-43 nuclear puncta were colocalized with heat shock proteins, including HSPA1A, a member of HSP70 that played a role in preventing its degradation under stress. The combined results suggest that the PML nuclear bodies may play a protective role with the help of HSPA1A against transient stress by temporarily storing TDP-43 in its native states, but the prolonged stress exhausted the capacity of heat shock proteins, resulting in the misfolding of TDP-43. We envision that the presented imaging approach can be applied to other RBPs and nuclear bodies to offer the potential for studying correlations between the folding states of RBPs and functions of nuclear bodies.

## Materials and Methods

### Plasmids

The genes encoding TDP-43, FUS, and TAF15 were amplified from the plasmids of the wtTDP43– tdTOMATOHA (Addgene, plasmid #28205), GST–TEV–FUS (Addgene, Plasmid #29629), pDEST_TAF15 (Addgene, Plasmid #26379), respectively. The SNAPf–TDP-43, FUS–SNAPf, and SNAPf–TAF15 constructs were then generated by using the polymerase incomplete primer extension (PIPE) method.^66^ In order to subclone these constructs into the XLone backbone, the Xlone–GFP (Addgene, plasmid #96930) plasmid was double digested with SpeI and KpnI restriction enzymes. The digested XLone vector was then fused with the insert of the respective constructs by in-fusion cloning method according to the manufacturer’s instructions (Takara Bio Inc.). The resulting products from the respective in-fusion cloning reactions were transformed into stbl3 One-Shot competent cells according to the manufacturer’s instructions (Invitrogen), to generate the Xlone plasmids of Xlone_SNAPf–TDP-43, Xlone_FUS–SNAPf, and Xlone_SNAPf–TAF15, respectively. The correct insertion for each plasmid was confirmed by a DNA Sanger sequencing service (GeneWiz Inc.).

### Cell culture and stable cell line generation

HEK293T cells were purchased from ATCC^®^ (CRL-3216TM) and cultured in complete Dulbecco’s modified Eagle’s medium (DMEM, Gibco, 11995-065) supplemented with 10% fetal bovine serum (FBS, Gibco, 26140-079, LOT# 2098936) and 1× penicillin-streptomycin-glutamine (PSQ, Gibco, 10378-016) at 37°C and under 5% CO_2_ atmosphere in a HERA cell VIOS 160i CO_2_ incubator (Thermo Fisher Scientific). The cells were passaged using 1× TrypLE^TM^ Express (Gibco, 12605-028) when they reach at ∼ 90% confluency. To generate doxycycline-inducible stable cell lines, HEK293T cells were co-transfected with the Xlone plasmid encoding respective SNAP-tag fused RBP and a pCYL43 plasmid encoding PiggyBac transposase to stably integrate into the genomic DNA of the cells. The cells were then further selected with Blasticidin S (Sigma Aldrich), and then single colonies were isolated using cloning cylinders to generate the respective stable cell line. The expression of SNAP-tag fused RBP for each stable cell line was verified by treating with doxycycline (Sigma Aldrich) and SNAP-Cell Oregon Green (NEB).

### Immunofluorescence staining

Cells on coverslips were fixed with 4% formaldehyde for 20 minutes at room temperature following washing with phosphate buffer saline (PBS, pH = 7.4). After washing with PBS three times for 10 minutes per each time, cells were permeabilized with 0.1% Triton X-100 in PBS for 10 minutes at room temperature and then washed with PBS three times for 10 minutes per each time. Cells were then blocked with 3% bovine serum albumin (BSA, Sigma Aldrich, A9085) solution in PBS for an hour at room temperature, followed by staining with primary antibodies diluted in PBS containing 2% BSA and 0.1% Tween 20 for overnight at 4 °C. Primary antibodies and their dilutions used were as follows: rabbit anti-PML (Abcam, ab179466, 1:100), rabbit anti-nucleophosmin (Abcam, ab52644, 1:50), rabbit anti-HSF1 (Invitrogen, PA5-89420, 1:100), rabbit anti-SFPQ (Abcam, ab177149, 1:200), mouse anti-SC35 (Abcam, ab11826, 1:200), rabbit anti-HSP70 (Abcam, ab181606, 1:50). After washing with PBS three times for 10 minutes per each time, cells were stained with secondary antibodies diluted in PBS containing 2% BSA and 0.1% Tween 20 for an hour at room temperature in the dark. Secondary antibodies and their dilutions used were as follows: goat anti-Rabbit IgG (H+L) Highly Cross-Adsorbed Secondary Antibody, Alexa Fluor 568 (Invitrogen, A11036, 1:300) and Goat Anti-Mouse IgG H&L, Alexa Fluor 594 (Abcam, ab150116, 1:300). After washing with PBS three times for 10 minutes per each time, coverslips were mounted on glass slides with antifade mounting medium with DAPI (Vectashield, Vector Laboratories) and sealed with transparent nail polish. Cells were visualized by a confocal microscope equipped with a 60x objective lens.

### Triple-color imaging assay with confocal microscopy

The stable cell line of SNAPf–TDP-43, FUS–SNAPf, or SNAPf–TAF15 was seeded in complete DMEM containing 10% FBS and 1× PSQ at a density of 5 ∼ 6 × 10^5^ cells/dish in a poly-D-lysine coated, glass-bottomed dish (35 mm, MatTek Corporation). For inducible expression and labeling of SNAP-tag fused RBPs, stock solutions of doxycycline (2 mg/mL in DMSO) and SNAP-tag substrates **P0**, **P1** or **P2** (2 mM in DMSO) were prepared and stored as aliquots in the dark at –20 °C. When the cell line reached 40 ∼ 50% confluence, the stock solutions were diluted in fresh complete DMEM to generate a 2× solution of doxycycline (280 ng/mL) and the three SNAP-tag substrates (1 μM per each substrate). The resulting 2× solution was added to the cell line by replacing the half of the culture media to arrive at final concentrations of 140 ng/mL for doxycycline and 0.5 μM for each SNAP-tag substrate. The cell line was then incubated for 24 hours at 37 °C and under 5% CO_2_ atmosphere to allow the stable expression and labeling. CuET or AdOx was treated according to the procedure described below. After the cells were washed with fresh FluroBrite DMEM supplemented with 10% FBS to remove excess substrates, fluorescence from **P0**, **P1** and **P2** was visualized by a laser scanning confocal microscope (FluoView 1000, Olympus) with an objective lens of 60× magnification. Laser lines of 405 nm, 458 nm, and 543 nm were used for the excitation of **P0**, **P1** and **P2**, respectively.

### Fluorescence lifetime imaging assay

The stable cell line of SNAPf–TDP-43, FUS–SNAPf, or SNAPf-TAF15 was seeded in complete DMEM containing 10% FBS and 1× PSQ at a density of 5 ∼ 6 × 10^5^ cells/dish in a poly-D-lysine coated, glass-bottomed dish (35 mm, MatTek Corporation). For inducible expression and labeling of SNAP-tag fused RBPs, stock solutions of doxycycline (2 mg/mL in DMSO) and SNAP-tag substrates **P3** or **P4** (2 mM in DMSO) were prepared and stored as aliquots in the dark at –20 °C. When the cell line reached 40 ∼ 50% confluence, the stock solutions were diluted in fresh complete DMEM to generate a 2× solution of doxycycline (280 ng/mL) and **P3** (1 μM) or a 2× solution of doxycycline (280 ng/mL) and **P4** (1 μM). The resulting 2× solution was added to the cell line by replacing the half of the culture media to arrive at final concentrations of 140 ng/mL for doxycycline and 0.5 μM for **P3** or **P4**. The cell line was then incubated for 24 hours at 37°C and under 5% CO_2_ atmosphere to allow the stable expression and labeling. CuET or AdOx was treated according to the procedure described below. After the cells were washed with fresh FluoroBrite DMEM supplemented with 10% FBS to remove excess substrate, the fluorescence lifetime images and decays of **P3** or **P4** in the cell line were obtained by a Zeiss LSM 880 confocal microscope equipped with PicoQuant-FLIM LSM upgrade kit using an objective lens of 60× magnification. Pulsed-laser lines of 440 and 485 nm with repetition rates of 10 and 26.6 MHz were used for the excitation of **P3** and **P4**, respectively. The acquired lifetime images were then analyzed by using SymphoTime software (PicoQuant). The pixels in the region of interest (ROI) were selected, and multi-exponential reconvolution model was employed to calculate the best fit of the lifetime decay curves in the ROI.^67^

### Procedure for treatment with CuET or AdOx

CuET (Copper(II) Diethyldithiocarbamate) and AdOx (Adenosine-2’,3’-dialdehyde) were purchased from TCI and Sigma Aldrich, respectively. Stock solutions of CuET (1 mM in DMSO) and AdOx (50 mM in DMSO) were prepared and stored as aliquots in the dark at –20 °C. CuET was added to the cell line of SNAPf–TDP-43 with a final concentration of 5 μM in the cell medium at 24 hours after treating with doxycycline and the respective SNAP-tag substrate. AdOx was added to the cell line of FUS–SNAPf or SNAPf–TAF15 with a final concentration of 25 μM in the cell medium at 3 hours before treating with doxycycline and the respective SNAP-tag substrate.

### Western Blotting

Cells were washed and then lysed for 10 minutes on ice with cold RIPA buffer (Thermo Fisher Scientific) supplemented with 1× protease inhibitor cocktail (Promega). Cell lysates were then centrifuged at 21,000×g for 15 minutes at 4 °C, and then total protein concentration in the supernatant was determined by BCA assay kit (Thermo Fisher Scientific). Proteins in the supernatant were separated on a 10% polyacrylamide gel by SDS-PAGE, followed by transferring to a PVDF membrane (Bio-Rad Laboratories) using Trans Blot Turbo system (Standard 25V, 1A, 30 min, Bio-Rad Laboratories). After the membrane was blocked with 5% skim fat milk in tris-buffered saline (TBS, wt/vol) for an hour at room temperature, it was probed with primary antibodies for overnight at 4 °C. Primary antibodies and their dilutions used were as follows: rabbit anti-ubiquitin (linkage-specific K48, Abcam, ab140601, 1:2000), rabbit anti-GAPDH (Proteintech, 10494-1-AP, 1:10000). The membrane was washed with TBS containing 0.5% Tween 20 (TBS-T) three times for 10 minutes per each time, and then incubated with anti-rabbit IgG HRP-linked secondary antibody (Cell Signaling Technology, 7074, 1:1000) in TBS-T for an hour at room temperature. After washing the membrane with TBS-T three times for 10 minutes per each time, it was incubated for 5 minutes with superSignal™ West Pico PLUS chemiluminescent substrate (Thermo Fisher). Western blot images were acquired by using a ChemiDoc XRS+ system (Bio-Rad Laboratories) and processed by using Image Lab software (Bio-Rad Laboratories).

### Pulse-chase experiment

The stable cell line of SNAPf–TDP-43 was seeded in complete DMEM containing 10% FBS and 1× PSQ at a density of 5 ∼ 6 × 10^5^ cells/dish in cell culture dishes (35 mm, Greiner Bio-One). Stock solutions of doxycycline (2 mg/mL in DMSO), SNAP-Cell TMR Star (2 mM in DMSO, NEB), and SNAP-Cell Block (2 mM in DMSO, NEB) were prepared and stored as aliquots in the dark at –20 °C. When the cell line reached 40 ∼ 50% confluence, the stock solutions of doxycycline and SNAP-Cell TMR Star were diluted in fresh complete DMEM to generate a 2× solution of doxycycline (280 ng/mL) and SNAP-Cell TMR Star (4 μM). The resulting 2× solution was added to the cell line by replacing the half of the culture media to arrive at final concentrations of 140 ng/mL for doxycycline and 2 μM for SNAP-Cell TMR Star. The cell line was then incubated at 37 °C and under 5% CO_2_ atmosphere to allow the stable expression and labeling. After 24 hours, the cells were washed with fresh complete DMEM to remove unreacted probe. The stock solution of SNAP-Cell Block was diluted in fresh complete DMEM to generate a 1× solution of SNAP-Cell Block (10 μM), which were then added to the cell line by replacing the culture medium. After incubating for an hour at 37 °C and under 5% CO_2_ atmosphere to block further labeling of newly synthesized protein, CuET was added to the cell line of SNAPf–TDP-43 with a final concentration of 5 μM in the cell medium for different time periods. The cells were then washed and lysed for 10 minutes on ice with cold RIPA buffer (Thermo Fisher Scientific) supplemented with 1× protease inhibitor cocktail (Promega). Cell lysates were then centrifuged at 21,000×g for 15 minutes at 4 °C, and then total protein concentration in the supernatant was determined by BCA assay kit (Thermo Fisher Scientific). Proteins in the supernatant were separated on a 10% polyacrylamide gel by SDS-PAGE, followed by visualizing TMR fluorescence on the gel by in-gel fluorescence scanning with Gel Doc imaging system (Bio-Rad Laboratories).

### General synthetic procedure

Unless otherwise stated, all starting materials, reagents and solvents were purchased from commercial sources and used as received. All reactions were carried out under an argon atmosphere with anhydrous solvents and monitored by thin layer chromatography analysis using Silicycle® glass backed, extra hard sheets (60 Å, 250 μm thickness). All products were purified either by column chromatography over silica gel (SiliaFlash® F60, 40-63 μm), or by reversed-phase preparative HPLC (Agilent 1260, Agilent Technologies) on a C18 column with a linear gradient of buffer A (water/acetonitrile/trifluoroacetic acid, 94.9:5.0:0.1 v/v/v) and buffer B (acetonitrile/water/trifluoroacetic acid, 94.9:5.0:0.1 v/v/v) over 20 min. ^1^H and ^13^C NMR spectra were acquired on a Bruker AV-III-HD-500 spectrometer (Bruker Corp.) in CDCl_3_ or DMSO-*d*6. Chemical shifts are reported in parts per million (ppm) on the δ scale relative to the CDCl_3_, DMSO-*d*6 residual solvent peak or tetramethylsilane (TMS) as an internal standard.

### Synthetic procedure for SNAP-tag substrate P0

**Scheme S1.**
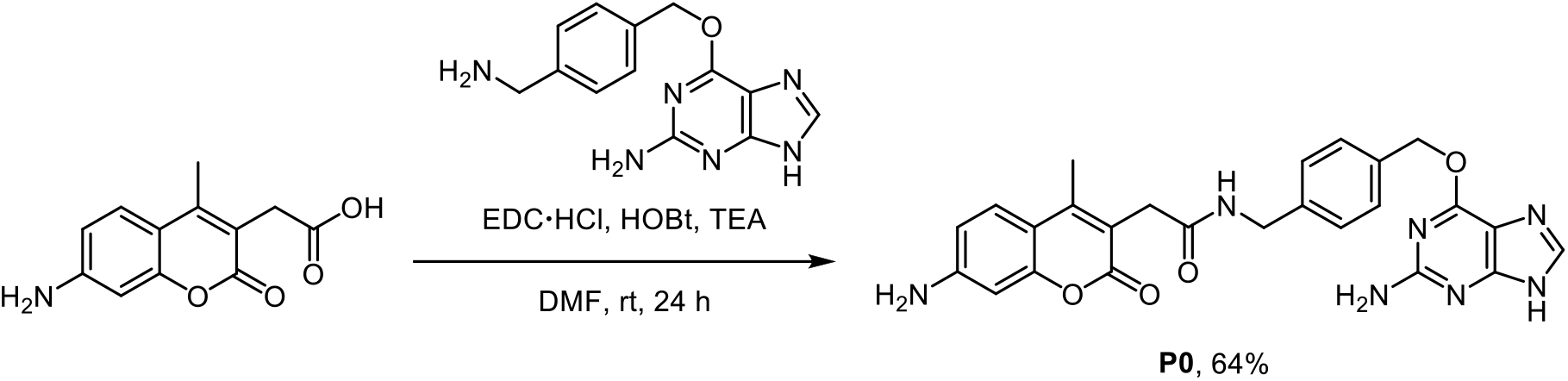
Synthetic scheme for SNAP-tag substrate **P0**.

7-Amino-4-methyl-3-coumarinyl acetic acid (23 mg, 0.1 mmol), *N*-(3-dimethylaminopropyl)-*N*′-ethylcarbodiimide hydrochloride (38 mg, 0.2 mmol, 2 equiv.), 1-hydroxybenzotriazole hydrate (31 mg, 0.2 mmol, 2 equiv.), and triethylamine (84 μL, 0.6 mmol, 6 equiv.) were dissolved in anhydrous dimethylformamide (2 mL). *O*^6^-[4-(aminomethyl)benzyl]guanine (41 mg, 0.15 mmol, 1.5 equiv.) was prepared according to the previously reported procedure^30^ and added to the solution in one portion. The reaction mixture was stirred for 24 h at room temperature under an argon atmosphere. After completion of the amide coupling, the product was directly purified by using reversed-phase preparative HPLC on a C18 column. The eluates were combined and lyophilized to afford **1** as a white solid (31 mg, yield 64%).

**P0** [**2-(7-amino-4-methyl-2-oxo-2*H*-chromen-3-yl)-*N*-(4-(((2-amino-9*H*-purin-6-yl)oxy)methyl)benzyl) acetamide**]: White solid. ^1^H NMR (500 MHz, DMSO-*d*_6_) δ 8.53 (s, 1H), 8.43 (t, *J* = 6.1 Hz, 1H), 7.49 (d, *J* = 8.2 Hz, 2H), 7.45 (d, *J* = 8.7 Hz, 1H), 7.29 (d, *J* = 8.1 Hz, 2H), 6.57 (dd, *J* = 8.7, 2.2 Hz, 1H), 6.41 (d, *J* = 2.2 Hz, 1H), 5.53 (s, 2H), 4.27 (d, *J* = 6.0 Hz, 2H), 3.46 (s, 2H), 2.27 (s, 3H). ^13^C NMR (126 MHz, DMSO-*d*_6_) δ 169.50, 161.67, 158.78, 157.88, 154.12, 153.20, 152.33, 150.01, 141.19, 140.25, 133.78, 128.91, 127.21, 126.35, 113.37, 111.37, 109.46, 106.85, 98.44, 68.37, 42.07, 33.91, 14.99. ESI-HRMS [M + H^+^]^+^ calculated 486.1884, observed 486.1878.

### Synthetic procedure for SNAP-tag substrates P1 and P2

**Scheme S2.**
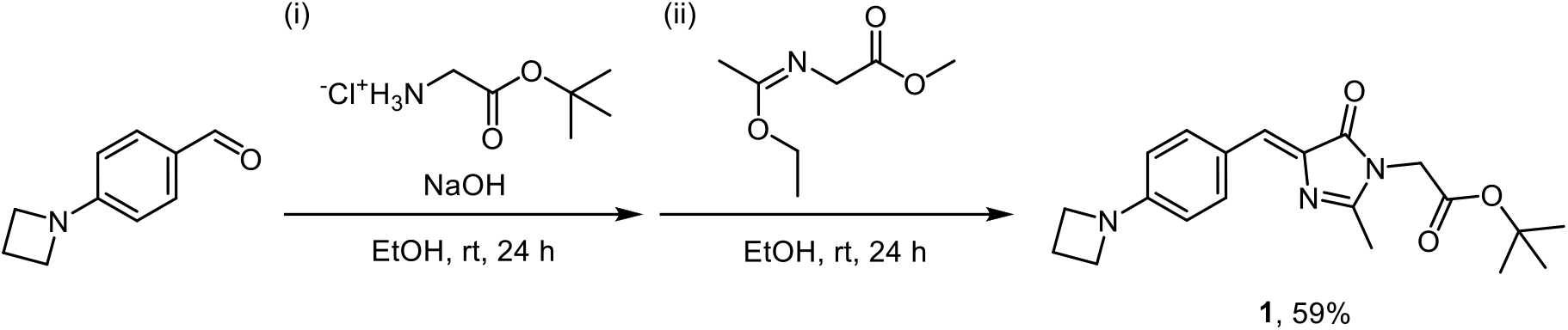
Synthetic scheme for precursor **1**.

4-(Azetidin-1-yl)benzaldehyde was prepared from 4-fluorobenzaldehyde according to the previously reported procedure.^68^ 4-(Azetidin-1-yl)benzaldehyde (3.00 g, 18.6 mmol), glycine tert-butyl ester hydrochloride (6.24 g, 37.2 mmol, 2 equiv.), and sodium hydroxide (1.49 g, 37.2 mmol, 2 equiv.) were dissolved in anhydrous ethanol (30 mL) and stirred for 24 h at room temperature under an argon atmosphere to yield the Schiff base intermediate. Methyl 2-(1-ethoxyethylideneamino)acetate (3.26 g, 20.5 mmol, 2 equiv.) was then prepared *in situ* according to the previously reported procedure^69^ and added dropwise to the reaction mixture. The reaction mixture was stirred for 24 h at room temperature under an argon atmosphere. After 24 h, water was added to the reaction mixture to quench the reaction. The product was extracted with dichloromethane, and the combined organic extracts were washed with brine, dried over sodium sulfate, and concentrated *in vacuo*. The crude residue was further purified by silica gel column chromatography to yield **1** as an orange oil (3.9 g, yield 59%).

**1 [((*Z*)-tert-butyl 2-(4-(4-(azetidin-1-yl)benzylidene)-2-methyl-5-oxo-4,5-dihydro-1*H*-imidazol-1-yl) acetate)]**. Orange oil. ^1^H NMR (500 MHz, DMSO-*d*_6_) δ 8.05 (d, *J* = 8.7 Hz, 2H), 6.90 (s, 1H), 6.41 (d, *J* = 8.8 Hz, 2H), 4.35 (s, 2H), 3.93 (t, *J* = 7.3 Hz, 4H), 2.35 (p, *J* = 7.3 Hz, 2H), 2.26 (s, 3H), 1.44 (s, 9H). ^13^C NMR (126 MHz, DMSO-*d*_6_) δ 169.05, 167.15, 159.17, 152.27, 133.93, 133.63, 127.15, 122.03, 110.25, 81.88, 51.21, 41.70, 27.48, 15.88, 14.81. ESI-HRMS [M + H^+^]^+^ calculated 356.1969, observed 356.1965.

**Scheme S3.**
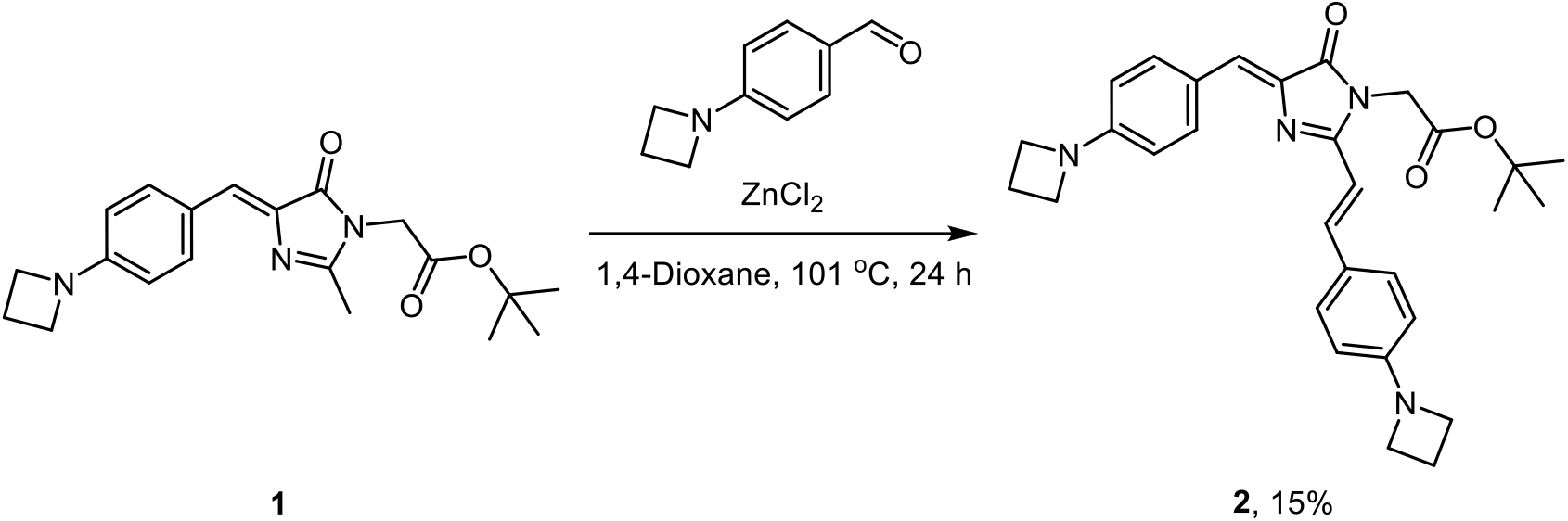
Synthetic scheme for precursor **2**.

Compound **2** (937 mg, 2.64 mmol), 4-(azetidin-1-yl)benzaldehyde (850 mg, 5.27 mmol, 2 equiv.), and zinc chloride (36 mg, 0.26 mmol, 0.1 equiv.) were dissolved in anhydrous 1,4-dioxane (20 mL). The reaction mixture was stirred under reflux for 24 h. After 24 h, the reaction mixture was cooled to room temperature and water was added to quench the reaction. The product was extracted with dichloromethane, and the combined organic layers were washed with brine, dried over sodium sulfate, and concentrated *in vacuo*. The product was further purified by silica gel column chromatography to yield **2** as a dark red solid (200 mg, yield 15%).

**2 [*Tert*-butyl 2-((*Z*)-4-(4-(azetidin-1-yl)benzylidene)-2-((*E*)-4-(azetidin-1-yl)styryl)-5-oxo-4,5-dihydro-1*H*-imidazol-1-yl)acetate**]: Dark red solid. ^1^H NMR (500 MHz, CDCl_3_) δ 8.14 (d, *J* = 8.5 Hz, 2H), 7.92 (d, *J* = 15.6 Hz, 1H), 7.43 (d, *J* = 8.6 Hz, 2H), 7.08 (s, 1H), 6.43 (d, *J* = 8.8 Hz, 2H), 6.40 – 6.36 (m, 3H), 4.41 (s, 2H), 3.99 (t, *J* = 7.3 Hz, 4H), 3.95 (t, *J* = 7.3 Hz, 4H), 2.43 – 2.40 (m, 2H), 2.39 – 2.36 (m, 2H), 1.44 (s, 9H). ^13^C NMR (126 MHz, CDCl_3_) δ 170.44, 167.16, 156.75, 152.77, 152.48, 140.24, 135.64, 134.26, 129.34, 127.90, 124.33, 123.95, 110.97, 110.77, 107.74, 82.82, 51.96, 51.78, 42.43, 28.06, 16.76, 16.67. ESI-HRMS [M + H^+^]^+^ calculated 499.2704, observed 499.2697.

**Scheme S4.**
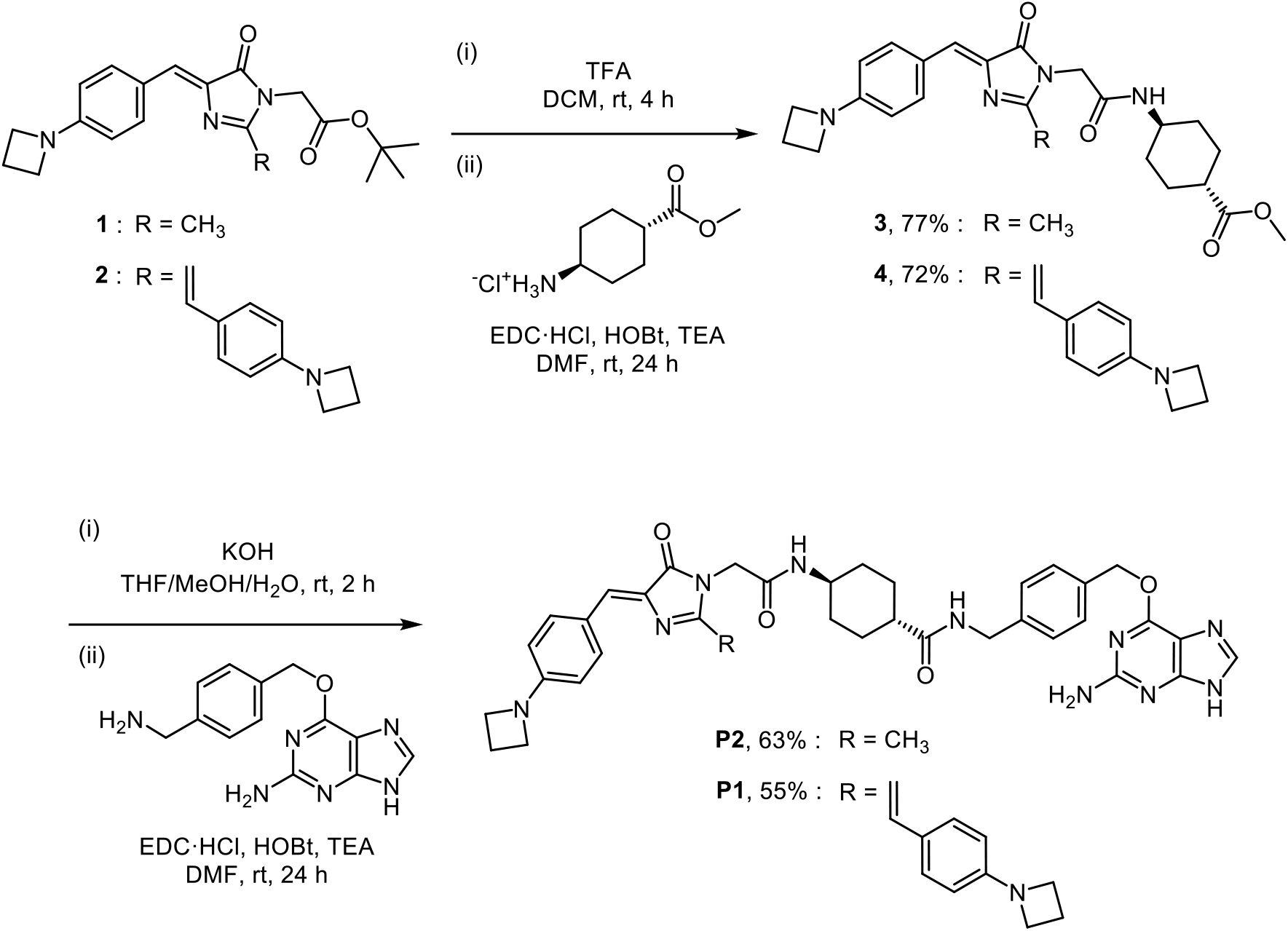
Synthetic scheme for precursor **3** and **4**, and SNAP-tag substrates **P2** and **P1**.

To a stirred solution of compound **1** (71 mg, 0.2 mmol) or **2** (100 mg, 0.2 mmol) in anhydrous dichloromethane (1 mL) was added dropwise trifluoroacetic acid (1 mL, excess equiv.). The reaction mixture was stirred for 4 h at room temperature. Upon complete deprotection of the Boc group, trifluoroacetic acid and dichloromethane were co-evaporated with toluene under reduced pressure to a dried crude residue. A solution of *N*-(3-dimethylaminopropyl)-*N*′-ethylcarbodiimide hydrochloride (77 mg, 0.4 mmol, 2 equiv.), 1-hydroxybenzotriazole hydrate (61 mg, 0.4 mmol, 2 equiv.), and triethylamine (169 μL, 1.2 mmol, 6 equiv.) in anhydrous dimethylformamide (4 mL) was added to the crude residue and stirred to dissolve. Methyl trans-4- aminocyclohexanecarboxylate hydrochloride (77 mg, 0.4 mmol, 2 equiv.) was added to the solution in one portion. The reaction mixture was stirred for 24 h at room temperature under an argon atmosphere. After completion of the amide coupling, water was added to the reaction mixture to quench the reaction. The product was extracted with dichloromethane, and the combined organic extracts were washed with brine, dried over sodium sulfate, and concentrated *in vacuo*. The product was further purified by silica gel column chromatography to yield **3** as a yellow solid (68 mg, yield 77%) or **4** as a dark red solid (84 mg, yield 72%), respectively.

**3 [(1*r*,4*r*)-Methyl 4-(2-((*Z*)-4-(4-(azetidin-1-yl)benzylidene)-2-methyl-5-oxo-4,5-dihydro-1*H*-imidazol-1-yl)acetamido)cyclohexanecarboxylate]:** Yellow solid. ^1^H NMR (500 MHz, DMSO-*d*_6_) δ 8.15 (d, *J* = 7.7 Hz, 1H), 8.05 (d, *J* = 8.7 Hz, 2H), 6.86 (s, 1H), 6.42 (d, *J* = 8.9 Hz, 2H), 4.20 (s, 2H), 3.93 (t, *J* = 7.3 Hz, 4H), 3.58 (s, 3H), 3.53 – 3.46 (m, 1H), 2.38 – 2.32 (m, 2H), 2.30 – 2.25 (m, 1H), 2.23 (s, 3H), 1.94 – 1.88 (m, 2H), 1.86 – 1.80 (m, 2H), 1.43 – 1.33 (m, 2H), 1.25 – 1.18 (m, 2H). ^13^C NMR (126 MHz, DMSO-*d*_6_) δ 175.09, 169.43, 165.80, 160.34, 152.36, 134.44, 133.66, 126.73, 122.30, 110.47, 51.39, 51.34, 47.46, 42.14, 41.36, 31.13, 27.42, 16.03, 15.22. ESI-HRMS [M + H^+^]^+^ calculated 439.2340, observed 439.2334.

**4 [(1*r*,4*r*)-methyl 4-(2-((*Z*)-4-(4-(azetidin-1-yl)benzylidene)-2-((*E*)-4-(azetidin-1-yl)styryl)-5-oxo-4,5-dihydro-1*H*-imidazol-1-yl)acetamido)cyclohexanecarboxylate].** Dark red solid. ^1^H NMR (500 MHz, CDCl_3_) δ 8.16 (d, *J* = 8.2 Hz, 2H), 7.96 (d, *J* = 15.6 Hz, 1H), 7.49 (d, *J* = 8.2 Hz, 2H), 7.11 (s, 1H), 6.54 (d, *J* = 15.6 Hz, 1H), 6.45 (d, *J* = 8.1 Hz, 2H), 6.41 (d, *J* = 8.0 Hz, 2H), 5.99 (d, *J* = 8.0 Hz, 1H), 4.38 (s, 2H), 4.04 (t, *J* = 7.3 Hz, 4H), 4.00 (t, *J* = 7.2 Hz, 4H), 3.80 – 3.72 (m, 1H), 3.66 (s, 3H), 2.48 – 2.45 (m, 2H), 2.43 – 2.40 (m, 2H), 2.21 – 2.16 (m, 1H), 2.02 – 1.96 (m, 4H), 1.56 – 1.47 (m, 2H), 1.16 – 1.09 (m, 2H). ^13^C NMR (126 MHz, CDCl_3_) δ 175.79, 170.84, 166.93, 156.22, 152.95, 152.64, 141.14, 134.93, 134.54, 129.72, 128.82, 124.13, 123.56, 110.94, 110.79, 107.02, 77.41, 77.16, 76.91, 51.97, 51.72, 48.15, 44.73, 42.25, 31.88, 29.83, 27.74, 16.76, 16.65. ESI-HRMS [M + H^+^]^+^ calculated 582.3075, observed 582.3067.

To a solution of compound **3** (44 mg, 0.1 mmol) or **4** (58 mg, 0.1 mmol) in a mixture of tetrahydrofuran/methanol/water (1.5 mL/1.0 mL/0.5 mL) was added pottasium hydroxide (28 mg, 0.5 mmol, 5 equiv.) in one portion. The reaction mixture was stirred at room temperature and the reaction progress was monitored by TLC. Upon complete deprotection of the methyl ester group, aqueous acetate buffer (pH 4.0, 0.1 M) were added to quench the reaction. The product was extracted with ethyl acetate, and the combined organic extracts were washed with brine, dried over sodium sulfate, and concentrated *in vacuo* to yield a crude residue. A solution of *N*-(3-dimethylaminopropyl)-*N*′-ethylcarbodiimide hydrochloride (38 mg, 0.2 mmol, 2 equiv.), 1-hydroxybenzotriazole hydrate (31 mg, 0.2 mmol, 2 equiv.), and triethylamine (84 μL, 0.6 mmol, 6 equiv.) in anhydrous DMF (2 mL) was added to the crude residue and stirred to dissolve. *O*^6^-[4- (aminomethyl)benzyl]guanine (54 mg, 0.2 mmol, 2.0 equiv.) was prepared according to the previously reported procedure^4^ and added to the solution in one portion. The reaction mixture was stirred for 24 h at room temperature under an argon atmosphere. After completion of the amide coupling, the product was directly purified using reversed-phase preparative HPLC on a C18 column. The eluates were combined and lyophilized to afford **P2** as an orange solid (43 mg, yield 63%) or **P1** as a dark red solid (46 mg, yield 55%), respectively.

**P2 [(1*r*,4*r*)-*N*-(4-(((2-amino-9*H*-purin-6-yl)oxy)methyl)benzyl)-4-(2-((*Z*)-4-(4-(azetidin-1-yl) benzylidene)-2-methyl-5-oxo-4,5-dihydro-1*H*-imidazol-1-yl)acetamido)cyclohexanecarboxamide].** Orange solid. ^1^H NMR (500 MHz, DMSO-*d*_6_) δ 8.27 (t, *J* = 6.1 Hz, 1H), 8.14 (d, *J* = 7.7 Hz, 1H), 8.05 (d, *J* = 8.7 Hz, 2H), 7.47 (d, *J* = 8.0 Hz, 2H), 7.25 (d, *J* = 7.9 Hz, 2H), 6.86 (s, 1H), 6.42 (d, *J* = 8.9 Hz, 2H), 5.48 (s, 2H), 4.25 (d, *J* = 5.8 Hz, 2H), 4.19 (s, 2H), 3.93 (t, *J* = 7.3 Hz, 4H), 3.52 – 3.45 (m, 1H), 2.38 – 2.32 (m, 2H), 2.23 (s, 3H), 2.16 – 2.09 (m, 1H), 1.88 – 1.81 (m, 2H), 1.81 – 1.75 (m, 2H), 1.46 – 1.38 (m,, 2H), 1.23 – 1.15 (m, 2H). ^13^C NMR (126 MHz, DMSO-*d*_6_) δ 174.70, 169.43, 165.73, 160.37, 152.38, 139.96, 134.38, 133.68, 128.75, 127.08, 126.76, 122.29, 110.49, 67.35, 51.41, 47.65, 43.14, 42.17, 41.60, 31.56, 28.17, 16.04, 15.24, 1.13. ESI-HRMS [M + H^+^]^+^ calculated 677.3307, observed 677.3296.

**P1 [(1*r*,4*r*)-*N*-(4-(((2-amino-9*H*-purin-6-yl)oxy)methyl)benzyl)-4-(2-((*Z*)-4-(4-(azetidin-1-yl) benzylidene)- 2-((*E*)-4-(azetidin-1-yl)styryl)-5-oxo-4,5-dihydro-1*H*-imidazol-1-yl)acetamido)cyclohexanecarboxamide].** Dark red solid. ^1^H NMR (500 MHz, DMSO-*d*_6_) δ 8.26 (t, *J* = 5.9 Hz, 1H), 8.20 (d, *J* = 7.8 Hz, 1H), 8.15 (d, *J* = 8.5 Hz, 2H), 7.79 (s, 1H), 7.79 (d, *J* = 15.5 Hz, 1H), 7.57 (d, *J* = 8.6 Hz, 2H), 7.44 (d, *J* = 8.0 Hz, 2H), 7.23 (d, *J* = 8.0 Hz, 2H), 6.83 (s, 1H), 6.70 (d, *J* = 15.6 Hz, 1H), 6.46 (d, *J* = 8.9 Hz, 2H), 6.43 (d, *J* = 8.7 Hz, 2H), 6.20 (d, *J* = 9.8 Hz, 2H), 5.44 (s, 2H), 4.37 (s, 2H), 4.24 (d, *J* = 5.6 Hz, 2H), 3.95 (t, *J* = 7.3 Hz, 4H), 3.91 (t, *J* = 7.3 Hz, 4H), 3.52 – 3.45 (m, 1H), 2.40 – 2.31 (m, 4H), 2.16 – 2.10 (m, 1H), 1.84 – 1.81 (m, 2H), 1.79 – 1.75 (m, 2H), 1.45 – 1.37 (m, 2H), 1.25 – 1.17 (m, 2H). ^13^C NMR (126 MHz, DMSO-*d*_6_) δ 174.70, 169.75, 165.91, 159.55, 157.72, 152.60, 152.11, 139.61, 139.16, 135.57, 135.16, 133.71, 129.50, 128.53, 127.08, 125.06, 123.74, 123.21, 110.80, 110.63, 108.53, 66.52, 51.59, 51.41, 47.61, 43.16, 42.17, 41.63, 31.61, 28.17, 16.14, 1.12. ESI-HRMS [M + H^+^]^+^ calculated 820.4042, observed 820.4029.

### Synthetic procedure for SNAP-tag substrate P3

**Scheme S5.**
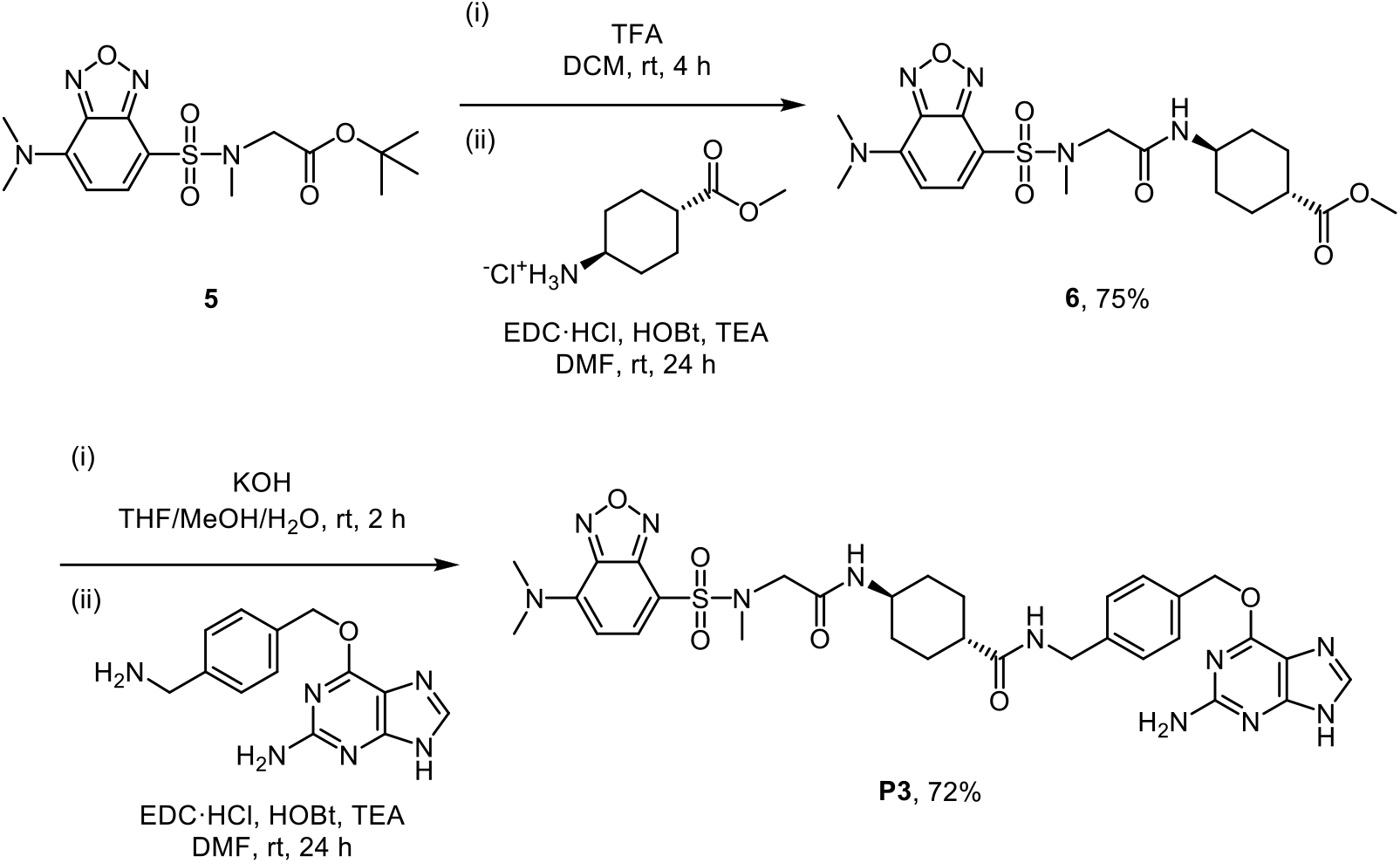
Synthetic scheme for precursor **6** and SNAP-tag substrates **P3**.

The synthetic precursor **5** was prepared from 4-chloro-2,1,3-benzoxadiazole according to the previously reported procedure.^70^ To a stirred solution of compound **5** (185 mg, 0.5 mmol) in anhydrous dichloromethane (2 mL) was added dropwise trifluoroacetic acid (2 mL, excess equiv.). The reaction mixture was stirred for 4 h at room temperature. Upon complete deprotection of the Boc group, trifluoroacetic acid and dichloromethane were co-evaporated with toluene under reduced pressure to a dried crude residue. A solution of *N*-(3-dimethylaminopropyl)-*N*′-ethylcarbodiimide hydrochloride (192 mg, 1.0 mmol, 2 equiv.), 1-hydroxybenzotriazole hydrate (153 mg, 1.0 mmol, 2 equiv.), and triethylamine (418 μL, 3.0 mmol, 6 equiv.) in anhydrous dimethylformamide (10 mL) was added to the crude residue and stirred to dissolve. Methyl trans-4-aminocyclohexanecarboxylate hydrochloride (194 mg, 1.0 mmol, 2 equiv.) was added to the solution in one portion. The reaction mixture was stirred for 24 h at room temperature under an argon atmosphere. After completion of the amide coupling, water was added to the reaction mixture to quench the reaction. The product was extracted with dichloromethane, and the combined organic extracts were washed with brine, dried over sodium sulfate, and concentrated *in vacuo*. The product was further purified by silica gel column chromatography to yield **6** as a yellow solid (170 mg, yield 75%).

**6** [**(1*r*,4*r*)-methyl 4-(2-(7-(dimethylamino)-*N*-methylbenzo[*c*][1,2,5]oxadiazole-4-sulfonamido)acetamido) cyclohexanecarboxylate**]: Yellow solid. ^1^H NMR (500 MHz, DMSO-*d*_6_) δ 7.81 (d, *J* = 8.3 Hz, 1H), 7.79 (d, *J* = 7.6 Hz, 1H), 6.24 (d, *J* = 8.5 Hz, 1H), 3.79 (s, 2H), 3.58 (s, 3H), 3.45 (s, 6H), 3.36 – 3.33 (m, 1H), 2.80 (s, 3H), 2.27 – 2.20 (m, 1H), 1.91 – 1.85 (m, 2H), 1.74 – 1.67 (m, 2H), 1.37 – 1.29 (m, 2H), 1.20 – 1.12 (m, 2H). ^13^C NMR (126 MHz, DMSO-*d*_6_) δ 175.08, 166.33, 146.73, 144.85, 142.91, 138.46, 107.28, 101.64, 51.87, 51.33, 47.10, 42.27, 41.37, 35.84, 31.00, 27.42. ESI-HRMS [M + H^+^]^+^ calculated 454.1755, observed 454.1746.

To a solution of compound **6** (44 mg, 0.1 mmol) in a mixture of tetrahydrofuran/methanol/water (1.5 mL/1.0 mL/0.5 mL) was added pottasium hydroxide (28 mg, 0.5 mmol, 5 equiv.) in one portion. The reaction mixture was stirred at room temperature and the reaction progress was monitored by TLC. Upon complete deprotection of the methyl ester group, aqueous acetate buffer (pH 4.0, 0.1 M) were added to quench the reaction. The product was extracted with ethyl acetate, and the combined organic extracts were washed with brine, dried over sodium sulfate, and concentrated *in vacuo* to yield a crude residue. A solution of *N*-(3-dimethylaminopropyl)-*N*′-ethylcarbodiimide hydrochloride (38 mg, 0.2 mmol, 2 equiv.), 1-hydroxybenzotriazole hydrate (31 mg, 0.2 mmol, 2 equiv.), and triethylamine (84 μL, 0.6 mmol, 6 equiv.) in anhydrous DMF (2 mL) was added to the crude residue and stirred to dissolve. *O*^6^-[4-(aminomethyl)benzyl]guanine (54 mg, 0.2 mmol, 2.0 equiv.) was prepared according to the previously reported procedure^4^ and added to the solution in one portion. The reaction mixture was stirred for 24 h at room temperature under an argon atmosphere. After completion of the amide coupling, the product was directly purified using reversed-phase preparative HPLC on a C18 column. The eluates were combined and lyophilized to afford **P3** as a yellow solid (50 mg, yield 72%).

**P3 [(1*r*,4*r*)-*N*-(4-(((2-amino-9*H*-purin-6-yl)oxy)methyl)benzyl)-4-(2-(7-(dimethylamino)-*N*-methylbenzo[c][1,2,5] oxadiazole-4-sulfonamido)acetamido)cyclohexanecarboxamide]:** Yellow solid. ^1^H NMR (500 MHz, DMSO-*d*_6_) δ 8.26 (t, *J* = 5.9 Hz, 1H), 8.03 (s, 1H), 7.81 (d, *J* = 8.4 Hz, 1H), 7.79 (d, *J* = 7.8 Hz, 1H), 7.46 (d, *J* = 8.1 Hz, 2H), 7.24 (d, *J* = 8.0 Hz, 2H), 6.58 (s, 1H), 6.24 (d, *J* = 8.5 Hz, 1H), 5.47 (s, 2H), 4.24 (d, *J* = 5.8 Hz, 2H), 3.79 (s, 2H), 3.45 (s, 6H), 3.37 – 3.32 (m, 1H), 2.80 (s, 3H), 2.13 – 2.07 (m, 1H), 1.77 – 1.71 (m, 4H), 1.42 – 1.34 (m, 2H), 1.18 – 1.10 (m, 2H). ^13^C NMR (126 MHz, DMSO-*d*_6_) δ 174.74, 166.30, 159.28, 159.19, 155.26, 146.77, 144.90, 142.97, 139.85, 138.55, 134.78, 128.69, 127.10, 107.21, 101.68, 67.04, 51.93, 47.31, 43.14, 42.32, 41.63, 35.91, 31.44, 28.20, 1.14. ESI-HRMS [M + H^+^]^+^ calculated 692.2722, observed 692.2717.

### Synthetic procedure for SNAP-tag substrate P4

**Scheme S6.**
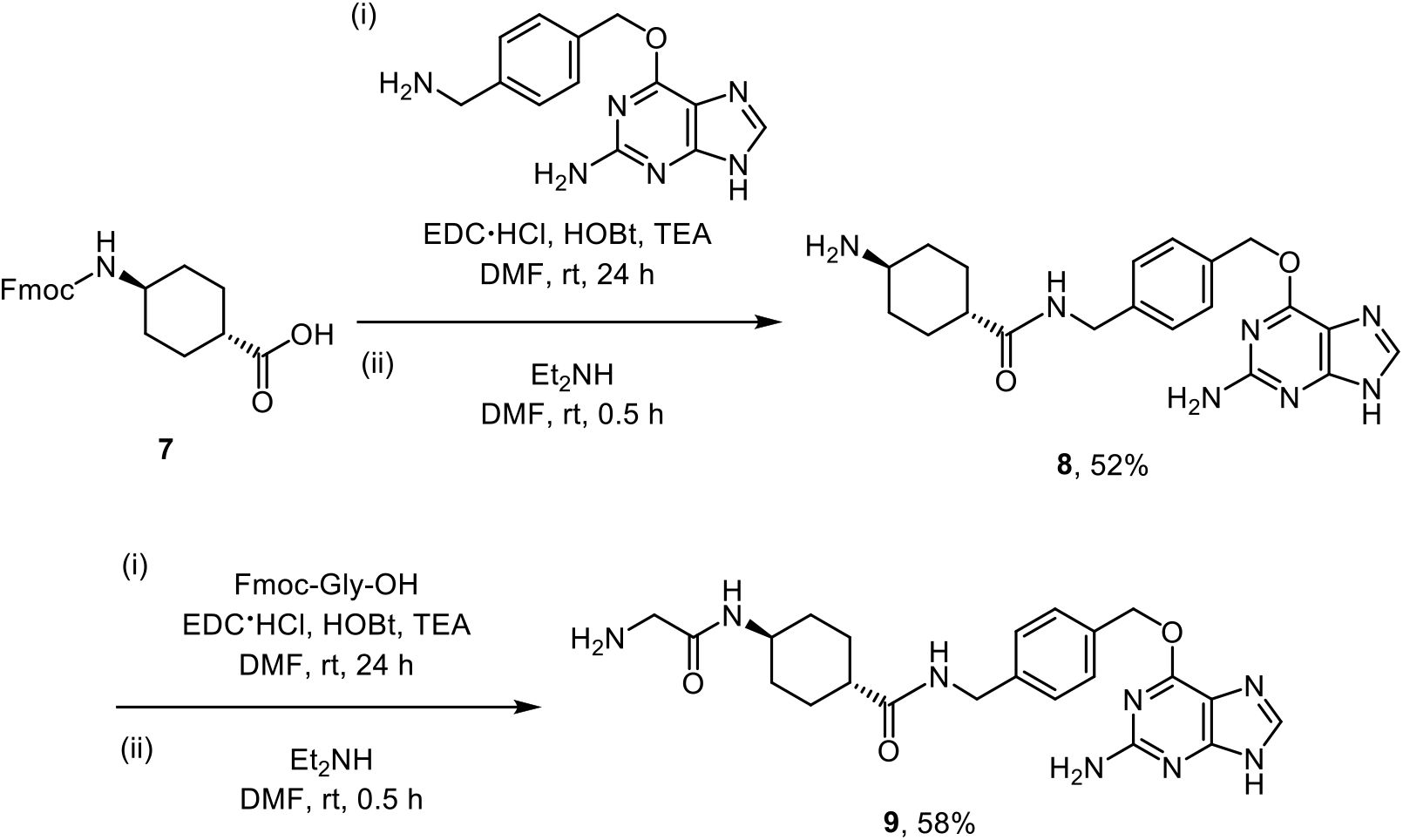
Synthetic scheme for precursor **8** and **9**.

The synthetic precursor **7** (183 mg, 0.5 mmol, 1 equiv.) was prepared from *trans*-4-aminocyclohexanecarboxylic acid hydrochloride according to the previously reported procedure.^71^ A solution of *N*-(3-dimethylaminopropyl)-*N*′-ethylcarbodiimide hydrochloride (192 mg, 1.0 mmol, 2 equiv.), 1-hydroxybenzotriazole hydrate (153 mg, 1.0 mmol, 2 equiv.), and triethylamine (418 μL, 3.0 mmol, 6 equiv.) in anhydrous dimethylformamide (10 mL) was added to **7** (183 mg, 0.5 mmol, 1 equiv.) and stirred to dissolve. *O*^6^-[4-(aminomethyl)benzyl]guanine (135 mg, 0.5 mmol, 1.0 equiv.) was prepared according to the previously reported procedure^4^ and added to the solution in one portion. The reaction mixture was stirred for 24 h at room temperature under an argon atmosphere. After completion of the amide coupling, water was added to the reaction mixture to quench the reaction. The product was extracted with ethyl acetate, and the combined organic extracts were washed with brine, dried over sodium sulfate, and concentrated *in vacuo* to yield a crude residue. To deprotect the Fmoc group, diethylamine (3 mL, excess equiv.) was added dropwise to a solution of the crude residue in anhydrous dimethylformamide (10 mL). The reaction mixture was stirred for 0.5 h at room temperature under an argon atmosphere. After complete deprotection of the Fmoc group, excess diethylamine was evaporated *in vacuo*. The product was further purified using reversed-phase preparative HPLC on a C18 column. The eluates were combined and lyophilized to afford **8** as a white solid (103 mg, yield 52%).

**8** [**(1*r*,4*r*)-4-amino-*N*-(4-(((2-amino-9*H*-purin-6-yl)oxy)methyl)benzyl)cyclohexanecarboxamide**]: White solid. ^1^H NMR (500 MHz, DMSO-*d*_6_) δ 8.39 (s, 1H), 8.37 (d, *J* = 6.0 Hz, 1H), 7.98 – 7.87 (d, 3H), 7.48 (d, *J* = 8.0 Hz, 2H), 7.25 (d, *J* = 8.0 Hz, 2H), 5.52 (s, 2H), 4.25 (d, *J* = 5.9 Hz, 2H), 3.01 – 2.93 (m, 1H), 2.17 – 2.11 (m, 1H), 1.97 – 1.94 (m, 2H), 1.82 – 1.79 (m, 2H), 1.48 – 1.40 (m, 2H), 1.34 – 1.26 (m, 2H). ^13^C NMR (126 MHz, DMSO-*d*_6_) δ 174.35, 158.98, 158.09, 153.67, 140.10, 134.10, 128.85, 127.13, 108.01, 68.00, 48.76, 42.53, 41.65, 29.54, 27.24. ESI-HRMS [M + H^+^]^+^ calculated 396.2136, observed 396.2142.

A solution of *N*-(3-dimethylaminopropyl)-*N*′-ethylcarbodiimide hydrochloride (77 mg, 0.4 mmol, 2 equiv.), 1-hydroxybenzotriazole hydrate (61 mg, 0.4 mmol, 2 equiv.), and triethylamine (167 μL, 1.2 mmol, 6 equiv.) in anhydrous dimethylformamide (5 mL) was added to Fmoc-Gly-OH (119 mg, 0.4 mmol, 2.0 equiv.) and stirred to dissolve. **8** (79 mg, 0.2 mmol, 1 equiv.) was added to the solution in one portion. The reaction mixture was stirred for 24 h at room temperature under an argon atmosphere. After completion of the amide coupling, water was added to the reaction mixture to quench the reaction. The product was extracted with ethyl acetate, and the combined organic extracts were washed with brine, dried over sodium sulfate, and concentrated *in vacuo* to yield a crude residue. To deprotect the Fmoc group, diethylamine (3 mL, excess equiv.) was added dropwise to a solution of the crude residue in anhydrous dimethylformamide (10 mL). The reaction mixture was stirred for 0.5 h at room temperature under an argon atmosphere. After complete deprotection of the Fmoc group, excess diethylamine was evaporated *in vacuo*. The product was further purified using reversed-phase preparative HPLC on a C18 column. The eluates were combined and lyophilized to afford **9** as a white solid (52 mg, yield 58%).

**9** [**(1*r*,4*r*)-*N*-(4-(((2-amino-9*H*-purin-6-yl)oxy)methyl)benzyl)-4-(2-aminoacetamido)cyclohexanecarboxamide**] : White solid. ^1^H NMR (500 MHz, DMSO-*d*_6_) δ 8.35 (d, *J* = 6.9 Hz, 2H), 8.32 (s, 1H), 8.16 (s, 1H), 8.11 (d, *J* = 5.8 Hz, 2H), 7.48 (d, *J* = 7.8 Hz, 2H), 7.25 (d, *J* = 7.8 Hz, 2H), 5.52 (s, 2H), 4.25 (d, *J* = 5.9 Hz, 2H), 3.56 – 3.52 (m, 1H), 3.50 (d, *J* = 7.3, 2H), 2.18 – 2.12 (m, 1H), 1.88 – 1.83 (m, 2H), 1.81 – 1.76 (m, 2H), 1.48 – 1.40 (m, 2H), 1.23 – 1.14 (m, 2H). ^13^C NMR (126 MHz, DMSO-*d*_6_) δ 174.77, 164.83, 159.10, 158.14, 140.13, 134.16, 128.77, 127.10, 108.59, 67.89, 47.68, 43.04, 41.63, 40.16, 31.50, 28.11. ESI-HRMS [M + H^+^]^+^ calculated 453.2357, observed 453.2351.

**Scheme S7.**
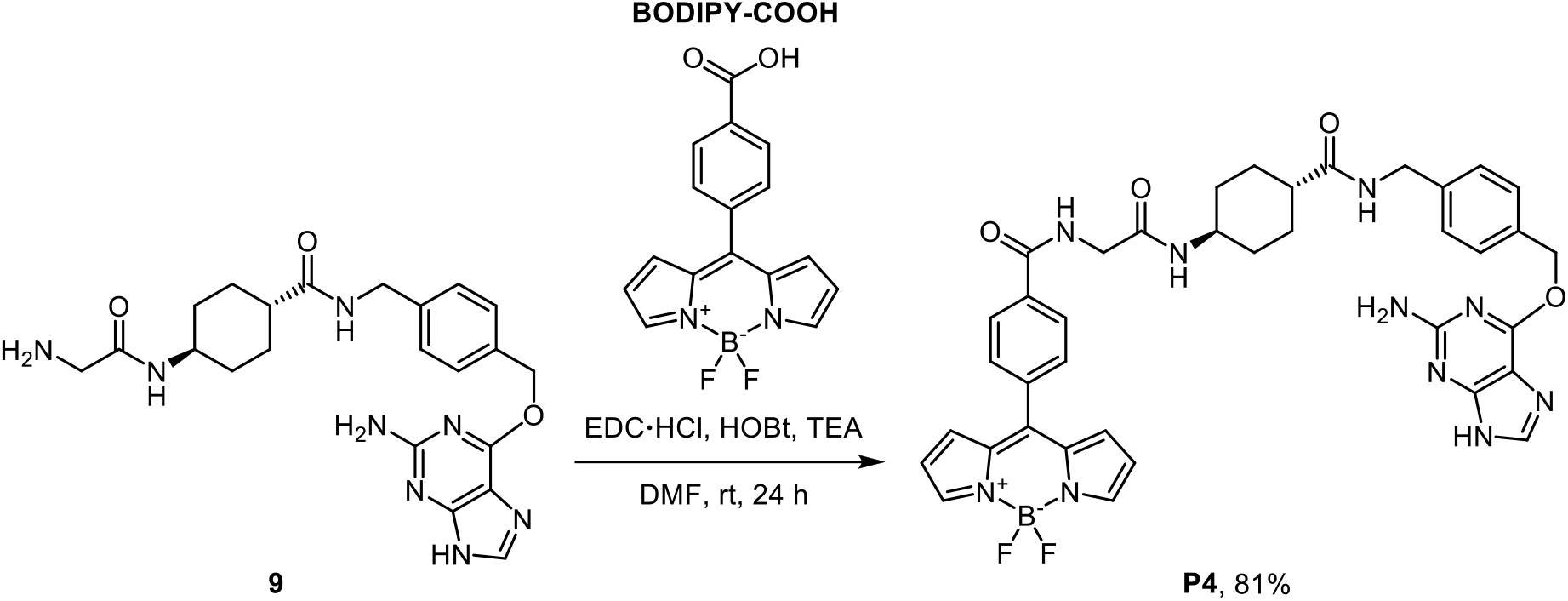
Synthetic scheme for SNAP-tag substrate **P4**.

The synthetic precursor **BODIPY-COOH** (31 mg, 0.1 mmol, 1 equiv.) was prepared according to the previously reported procedure.^72^ A solution of *N*-(3-dimethylaminopropyl)-*N*′-ethylcarbodiimide hydrochloride (38 mg, 0.2 mmol, 2 equiv.), 1-hydroxybenzotriazole hydrate (31 mg, 0.2 mmol, 2 equiv.), and triethylamine (82 μL, 0.6 mmol, 6 equiv.) in anhydrous dimethylformamide (2 mL) was added to **BODIPY-COOH** (31 mg, 0.1 mmol, 1 equiv.) and stirred to dissolve. **9** (31 mg, 0.1 mmol, 1 equiv.) added to the solution in one portion. The reaction mixture was stirred for 24 h at room temperature under an argon atmosphere. After completion of the amide coupling, the product was directly purified using reversed-phase preparative HPLC on a C18 column. The eluates were combined and lyophilized to afford **P4** as a red solid (61 mg, yield 81%).

**P4** [**10-(4-((2-(((1*r*,4*r*)-4-((4-(((2-amino-9*H*-purin-6-yl)oxy)methyl)benzyl)carbamoyl)cyclohexyl)amino)-2-oxoethyl)carbamoyl)phenyl)-5,5-difluoro-5*H*-dipyrrolo[1,2-*c*:2’,1’-*f*][1,3,2]diazaborinin-4-ium-5-uide]:** Red solid. ^1^H NMR (500 MHz, DMSO-*d*_6_) δ 8.91 (t, *J* = 5.9 Hz, 1H), 8.29 (t, *J* = 6.0 Hz, 1H), 8.18 (s, 2H), 8.09 (d, *J* = 8.3 Hz, 2H), 7.86 (d, *J* = 7.8 Hz, 1H), 7.79 (d, *J* = 8.3 Hz, 2H), 7.48 (d, *J* = 8.0 Hz, 2H), 7.26 (d, *J* = 7.9 Hz, 2H), 7.03 (d, *J* = 4.2 Hz, 2H), 6.71 (dd, *J* = 4.3, 1.9 Hz, 2H), 5.51 (s, 2H), 4.25 (d, *J* = 5.8 Hz, 2H), 3.87 (d, *J* = 5.9 Hz, 2H), 3.55 – 3.52 (m, 1H), 2.15 – 2.11 (m, 1H), 1.85 – 1.82 (m, 2H), 1.80 – 1.77 (m, 2H), 1.48 – 1.39 (m, 2H), 1.28 – 1.18 (m, 2H). ^13^C NMR (126 MHz, DMSO-*d*_6_) δ 174.73, 167.67, 165.55, 159.57, 145.21, 145.14, 139.63, 138.53, 138.36, 135.15, 134.10, 134.02, 131.79, 131.70, 128.53, 127.61, 127.08, 119.54, 66.51, 47.41, 43.19, 42.65, 41.62, 31.64, 28.27, 1.11. ^19^F NMR (471 MHz, DMSO-*d*_6_) δ -143.76, -143.82, -143.88, -143.95. ESI-HRMS [M + H^+^]^+^ calculated 747.3133, observed 747.3133.

## Acknowledgments

We thank support from the National Science Foundation CBET-1943696 (X.L.L.), the Burroughs Welcome Fund Career Award at the Scientific Interface 1013904 (X.Z.), Paul Berg Early Career Professorship (X.Z.), Lloyd and Dottie Huck Early Career Award (X.Z.), Sloan Research Fellowship FG-2018-10958 (X.Z.), PEW Biomedical Scholars Program (X.Z.), and National Institute of General Medical Sciences R35 GM133484 (X.Z.). We thank Dr. Stephen Benkovic’s lab for access to his ChemiDoc imaging instrument for western blotting. We thank Dr. Xiaojun Lance Lian’s lab for technical assistance of generating stable cell lines. We thank Dr. Gang Ning and Ms. Missy Hazen of the Penn State Microscopy Core Facility and Dr. Tatiana Laremore of the Penn State Proteomics and Mass Spectrometry Core Facility for technical assistance.

## Supplementary Figures

**Figure S1.**
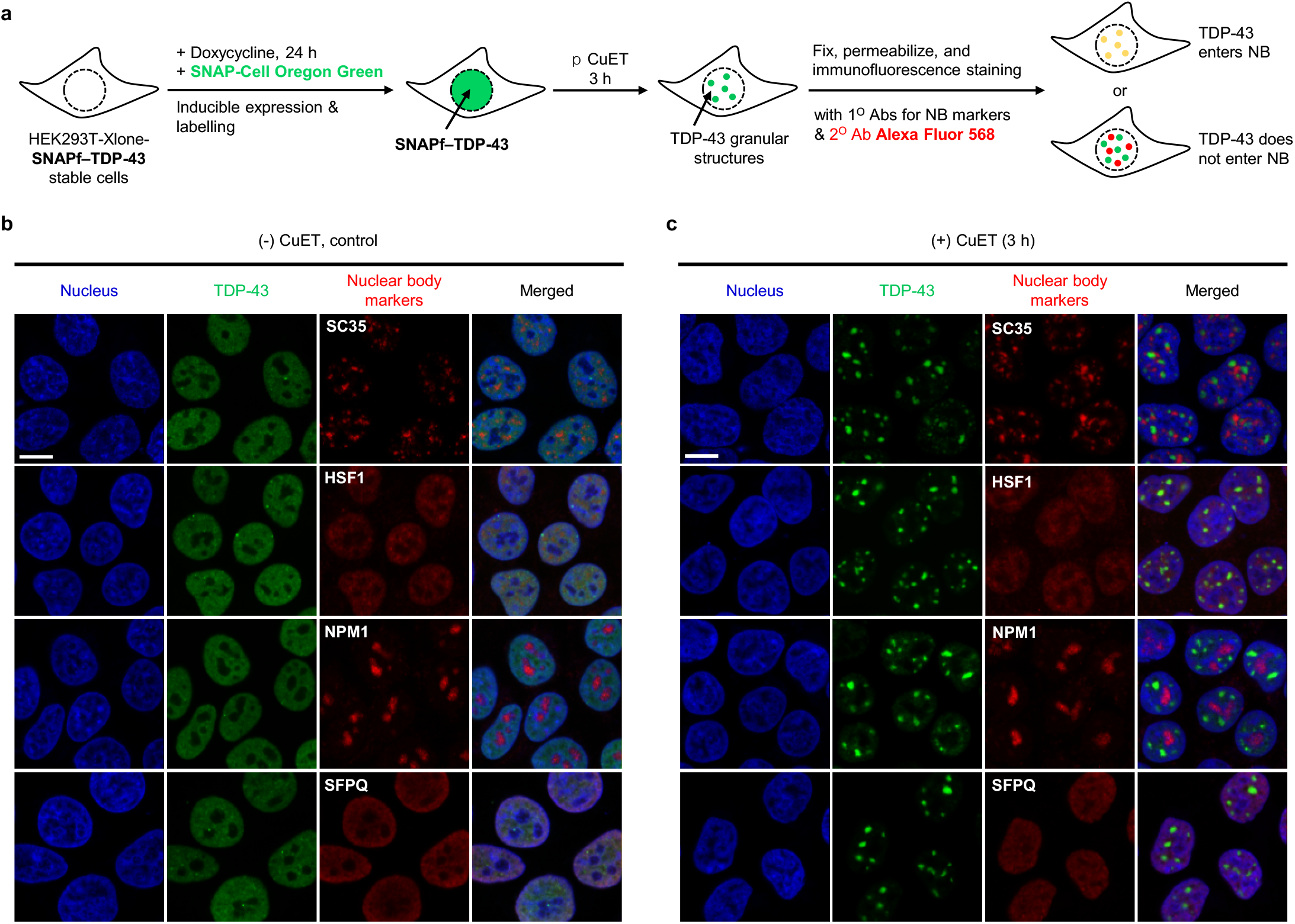
Immunofluorescence experiments show that TDP-43 does not enter other nuclear bodies, including nuclear speckles, nuclear stress bodies, nucleoli, and paraspeckles, upon treatment with CuET. (**a**) Schematic illustration of the procedure for the immunofluorescence experiments with anti-nuclear body makers (red) for cells stably expressing SNAPf–TDP-43 (green). SNAPf–TDP-43 stable cells were treated with doxycycline (140 ng/mL) and SNAP-Cell Oregon Green (0.5 μM) for 24 hours for the inducible expression and labeling of SNAPf–TDP-43, respectively. After the cells were further treated with CuET (5 μM) for 3 hours, they were fixed, permeabilized, and stained with the primary antibodies for several different nuclear body markers, and the secondary antibody conjugated with red fluorophore. (**b**), (**c**) Immunofluorescence imaging with anti-SC35 (for nuclear speckles), anti-HSF1 (for nuclear stress bodies), anti-NPM1 (for nucleoli), or anti-SFPQ (for paraspeckles) for cells stably expressing SNAPf-TDP-43 (labeled by SNAP-Cell Oregon Green, green) (b) without CuET treatment, or (c) upon treatment with CuET for 3 hours. Scale bar = 10 µm.

**Figure S2.**
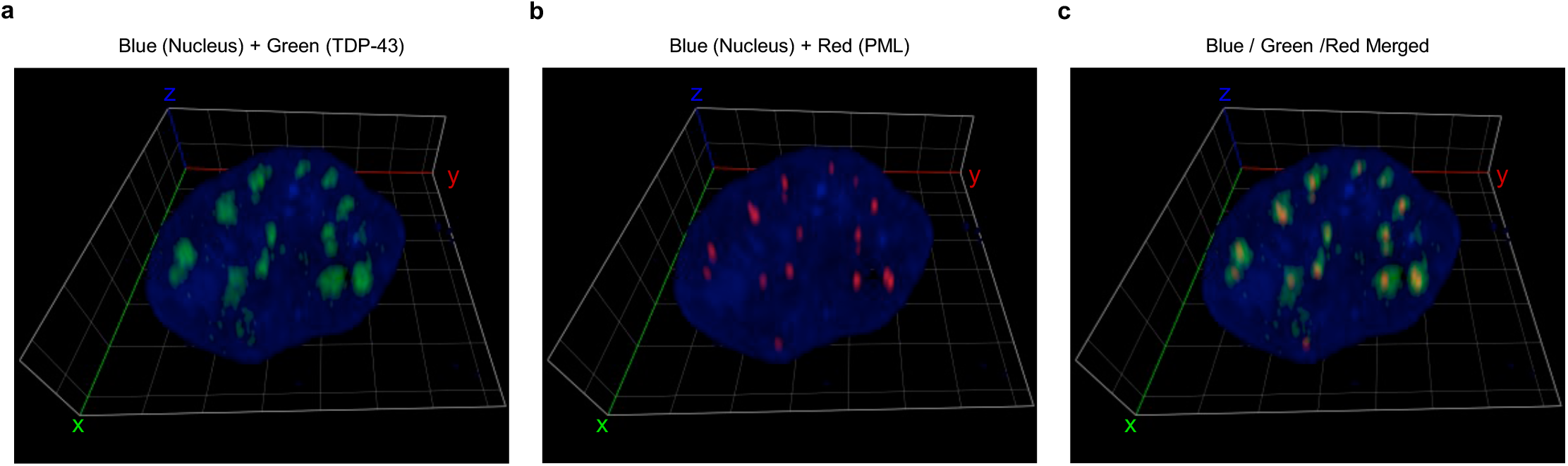
Z-stack and Airyscan imaging experiments reveal that PML nuclear bodies reside in the core of the TDP-43 granular structures formed by CuET treatment. Three-dimensional z-stack reconstruction of Airyscan imaging for the cell stably expressing SNAPf–TDP-43 (labeled by SNAP-Cell Oregon Green, green) upon treatment with CuET for 1 h. PML nuclear bodies were labeled with anti-PML (red) by immunofluorescence. (**a**) Merged image of blue (DAPI) and green (TDP-43). (**b**) Merged image of blue (DAPI) and red (PML). (**c**) Merged image of blue (DAPI), green (TDP-43) and red (PML). The unit of the grid is 3 µm.

**Figure S3.**
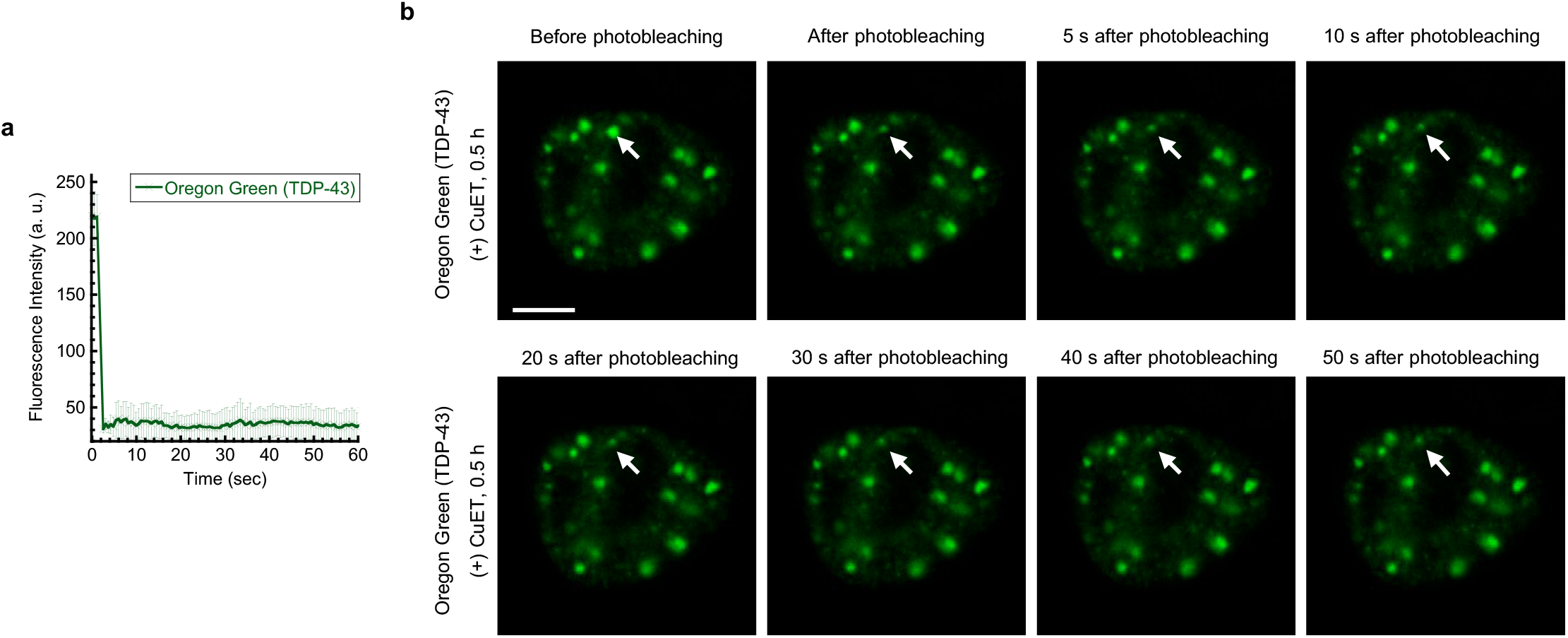
Fluorescence recovery after photobleaching experiments indicate that TDP-43 does not exhibit dynamic properties in the granular structures formed by CuET treatment. (**a**) Time-course of the fluorescence intensity profile at regions of interest from Oregon Green (TDP-43) channel in cells stably expressing SNAPf–TDP-43 (labeled by SNAP-Cell Oregon Green, green) upon treatment with CuET for 0.5 h. The error bar represents standard deviation from three independent measurements at each respective time point. (**b**) Representative time-lapse images of cells stably expressing SNAPf–TDP-43 (labeled by SNAP-Cell Oregon Green, green) with CuET treatment before and after photobleaching. The white arrow indicates the region of interest for the FRAP analysis. Scale bar = 5 µm.

**Figure S4.**
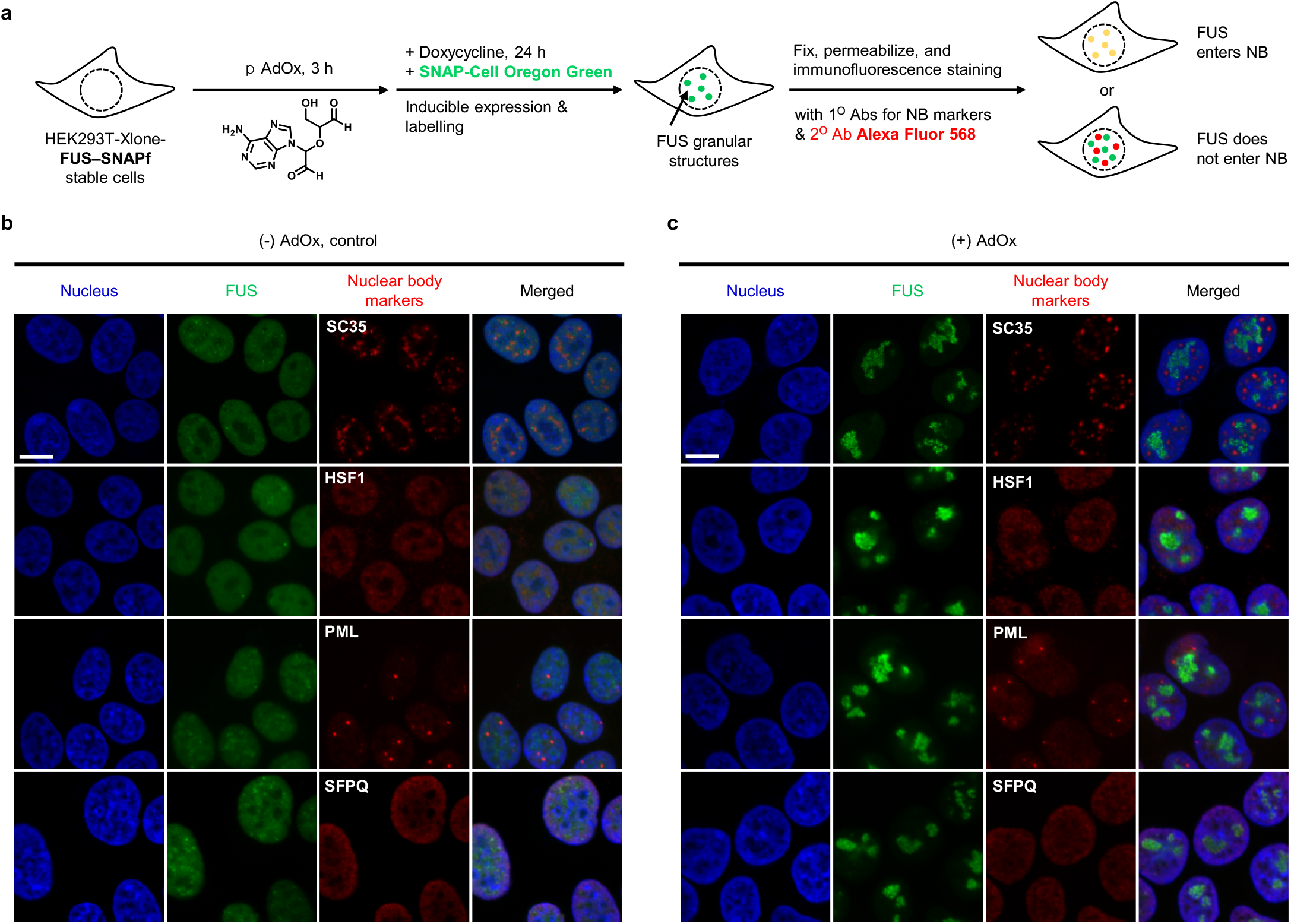
Immunofluorescence experiments show that FUS does not enter other nuclear bodies, including nuclear speckles, nuclear stress bodies, PML nuclear bodies, and paraspeckles, upon pretreatment with AdOx. (**a**) Schematic illustration of the procedure for the immunofluorescence experiments with anti-nuclear body makers (red) for cells stably expressing FUS–SNAPf (green). FUS–SNAPf stable cells were pretreated with AdOx (25 μM) or not 3 hours before they were further treated with doxycycline (140 ng/mL) and SNAP-Cell Oregon Green (0.5 μM) for 24 hours for the inducible expression and labeling of FUS–SNAPf, respectively. The cells were then fixed, permeabilized, and stained with the primary antibodies for several different nuclear body markers, and the secondary antibody conjugated with red fluorophore. (**b**), (**c**) Immunofluorescence imaging with anti-SC35 (for nuclear speckles), anti-HSF1 (for nuclear stress bodies), anti-PML (for the PML nuclear bodies), or anti-SFPQ (for paraspeckles) for cells stably expressing FUS–SNAPf (labeled by SNAP-Cell Oregon Green, green) (b) without AdOx pretreatment, or (c) with AdOx pretreatment. Scale bar = 10 µm.

**Figure S5.**
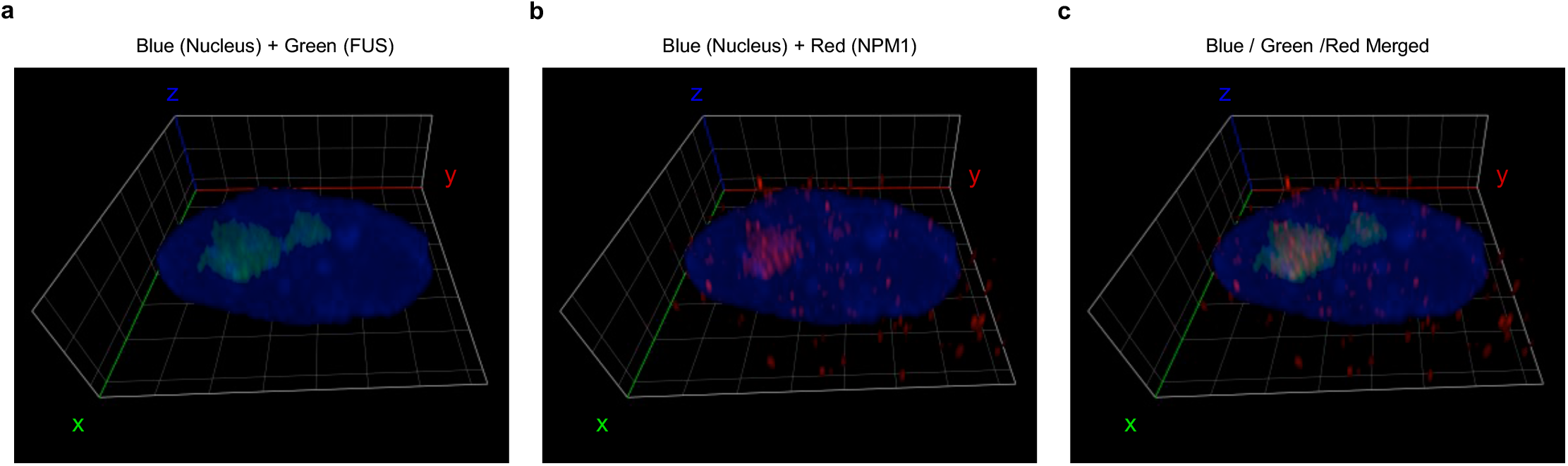
Z-stack and Airyscan imaging experiments reveal that FUS resides within the nucleolus when pretreated with AdOx. Three-dimensional z-stack reconstruction of Airyscan imaging for the cell stably expressing FUS–SNAPf (labeled by SNAP-Cell Oregon Green, green) upon AdOx treatment. Nucleoli were labeled with anti-NPM1 (red) by immunofluorescence. (**a**) Merged image of blue (DAPI) and green (FUS). (**b**) Merged image of blue (DAPI) and red (NPM1). (**c**) Merged image of blue (DAPI), green (FUS) and red (NPM1). The unit of the grid is 3 µm.

**Figure S6.**
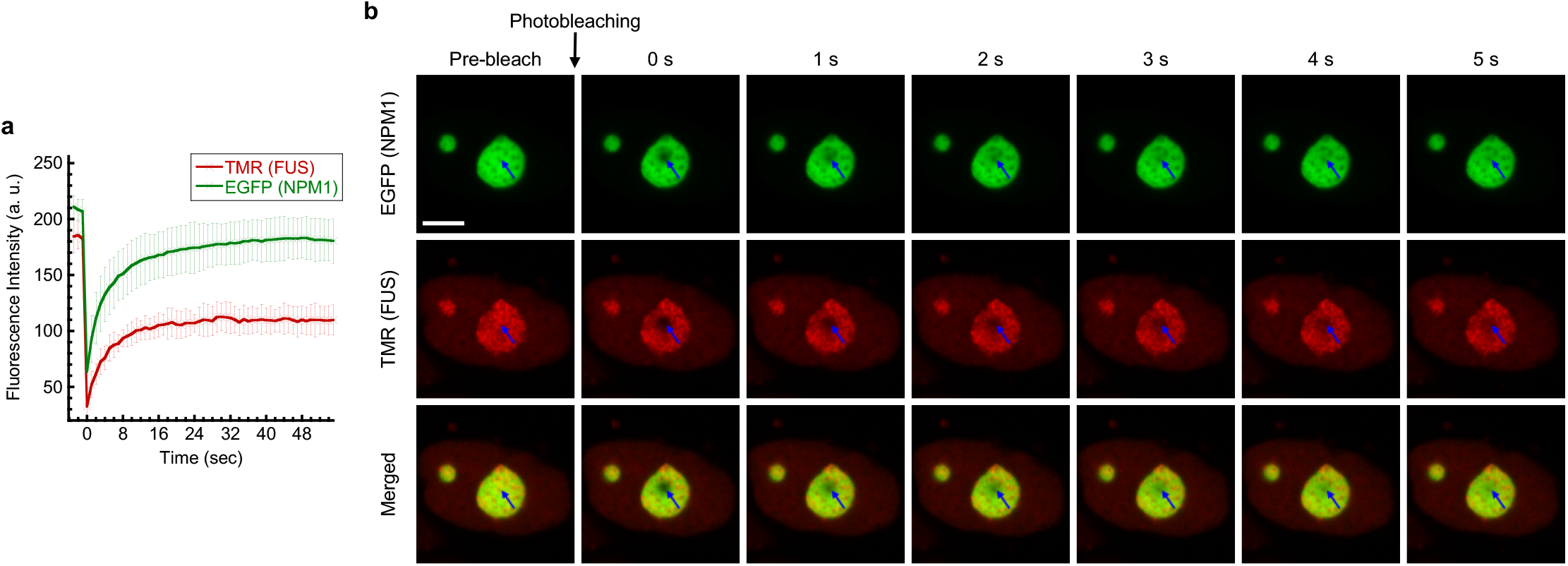
Fluorescence recovery after photobleaching experiments indicate that FUS remains diffuse in the nucleolus when pretreated with AdOx. (**a**) Time-course of the fluorescence intensity profiles at regions of interest from EGFP (NPM1) and TMR (FUS) channels in cells co-expressing EGFP–NPM1 (green) and FUS– SNAPf (labeled by SNAP-Cell TMR Star, red) with AdOx pretreatment. The error bar represents standard deviation from three independent measurements at each respective time point. (**b**) Representative time-lapse images of live cells co-expressing EGFP–NPM1 (green) and FUS–SNAPf (labeled by SNAP-Cell TMR Star, red) with AdOx pretreatment before and after photobleaching. The blue arrow indicates the region of interest for the FRAP analysis. Scale bar = 5 µm.

**Figure S7.**
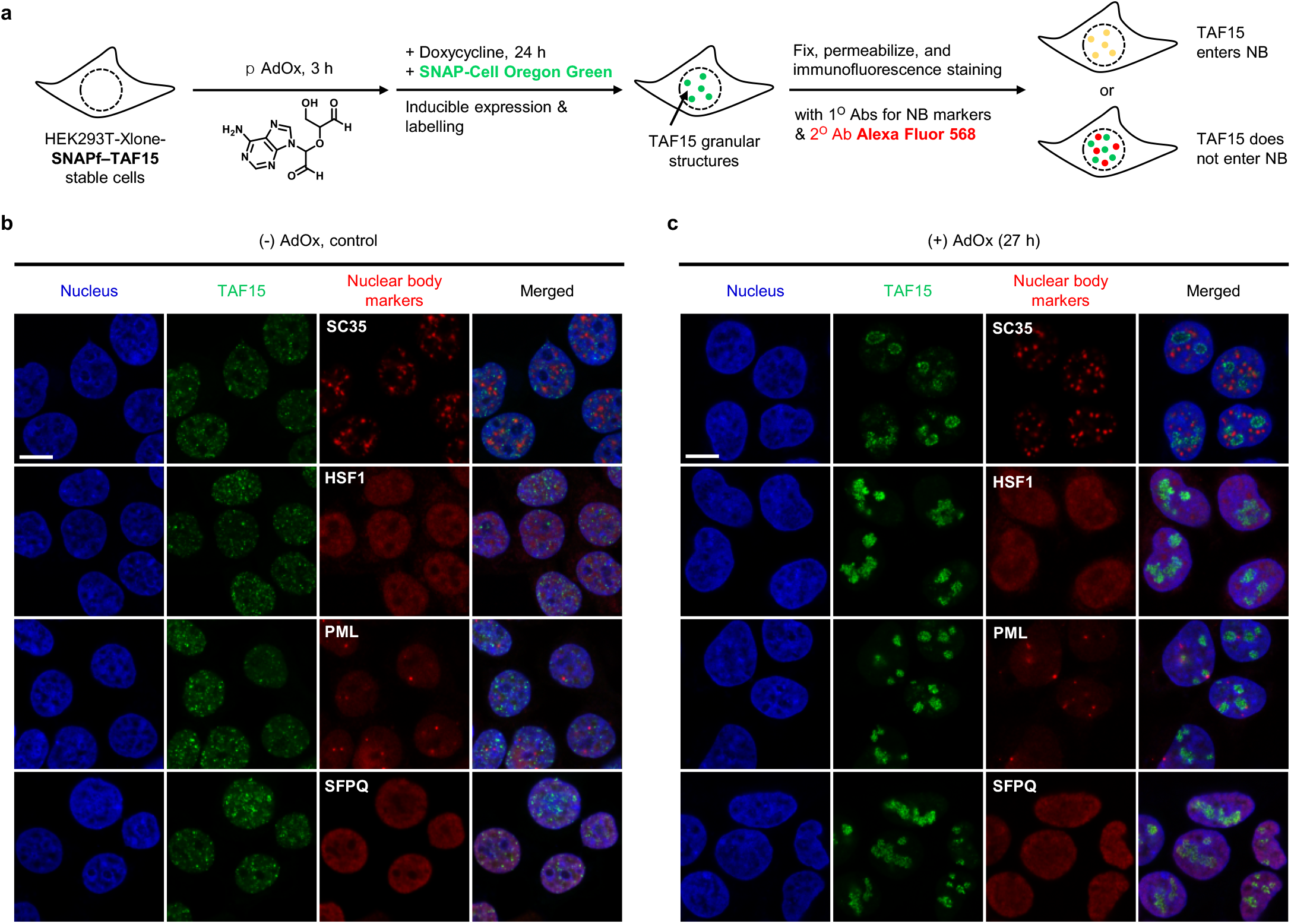
Immunofluorescence experiments show that TAF15 does not enter other nuclear bodies, including nuclear speckles, nuclear stress bodies, PML nuclear bodies, and paraspeckles, upon pretreatment with AdOx. (**a**) Schematic illustration of the general experimental procedure for the immunofluorescence imaging with anti-nuclear body makers (red) for cells stably expressing SNAPf–TAF15 (green). SNAPf–TAF15 stable cells were pretreated with AdOx (25 μM) or not 3 hours before they were further treated with doxycycline (140 ng/mL) and SNAP-Cell Oregon Green (0.5 μM) for 24 hours for the inducible expression and labeling of SNAPf–TAF15, respectively. The cells were then fixed, permeabilized, and stained with the primary antibodies for several different nuclear body markers, and the secondary antibody conjugated with red fluorophore. (**b**), (**c**) Immunofluorescence imaging with anti-SC35 (for nuclear speckles), anti-HSF1 (for nuclear stress bodies), anti-PML (for the PML nuclear bodies), or anti-SFPQ (for the paraspeckles) for cells stably expressing SNAPf–TAF15 (labeled by SNAP-Cell Oregon Green) (b) without AdOx pretreatment, or (c) with AdOx pretreatment. Scale bar = 10 µm.

**Figure S8.**
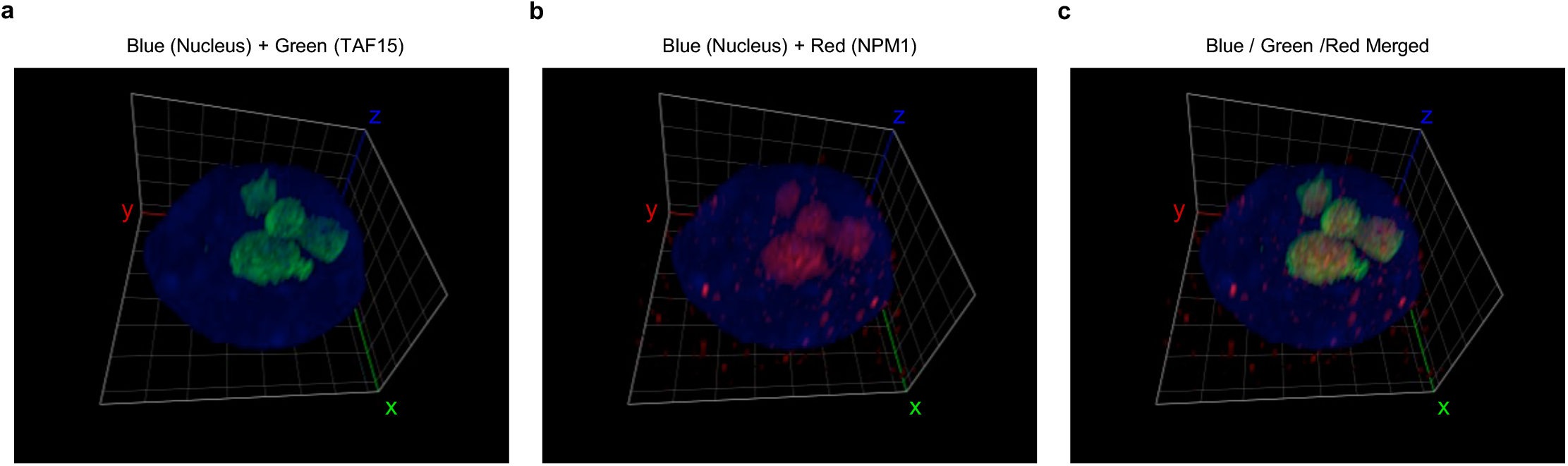
Z-stack and Airyscan imaging experiments reveal that TAF15 resides within the nucleolus when pretreated AdOx. Three-dimensional z-stack reconstruction of Airyscan imaging for the cell stably expressing SNAPf–TAF15 (labeled by SNAP-Cell Oregon Green, green) upon AdOx treatment. The nucleoli were labeled with anti-NPM1 (red) by immunofluorescence. (**a**) Merged image of blue (DAPI) and green (TAF15). (**b**) Merged image of blue (DAPI) and red (NPM1). (**c**) Merged image of blue (DAPI), green (TAF15) and red (NPM1). The unit of the grid is 3 µm.

**Figure S9.**
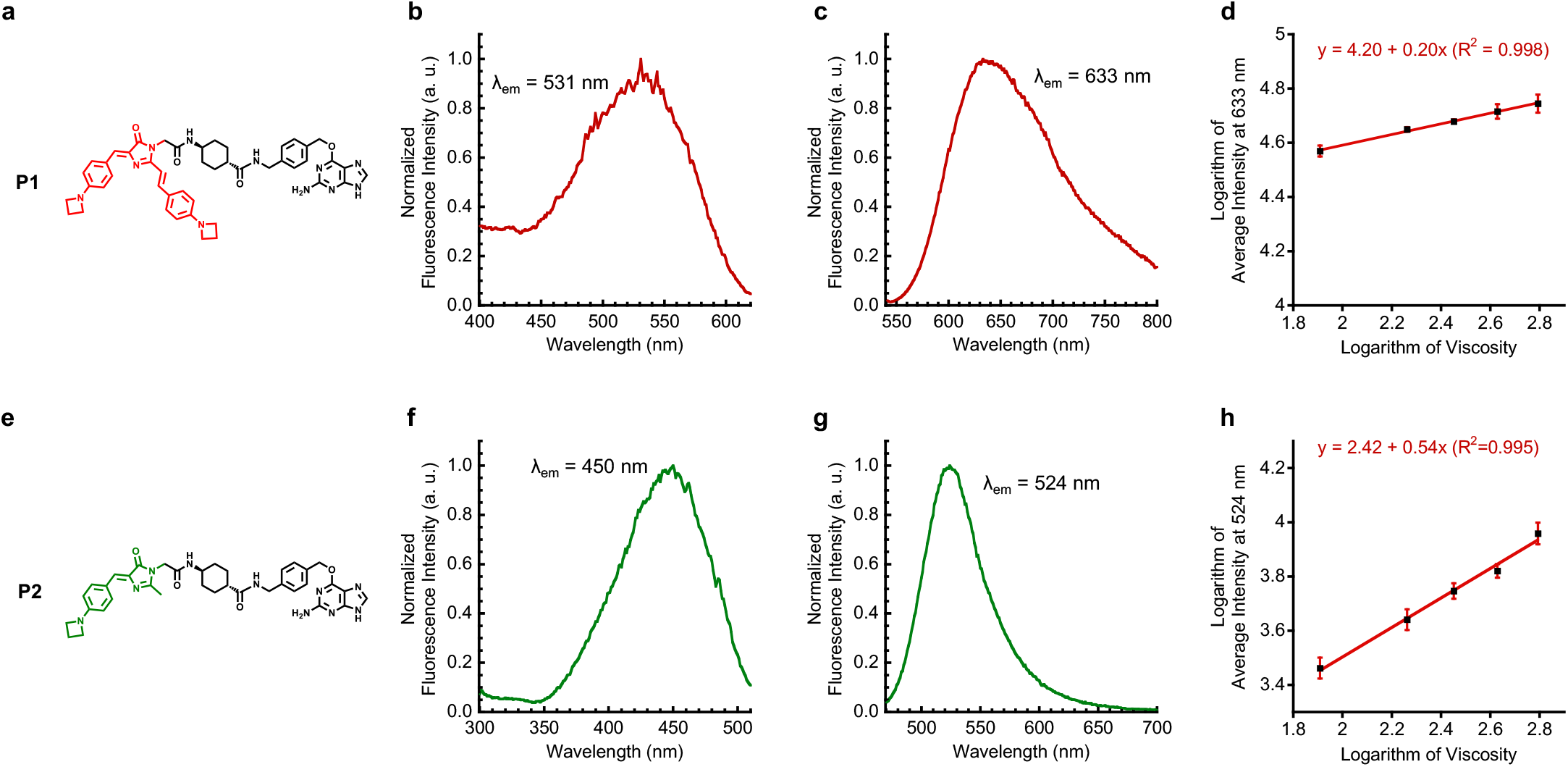
Photophysical characterization of P1 and P2 shows that they are spectrally orthogonal one another and have distinct viscosity sensitivity. (**a**) Structure of **P1**. (**b**) Fluorescence excitation spectrum of **P1** (5 μM) in glycerol. (**c**) Fluorescence emission spectrum of **P1** (5 μM) in glycerol. (**d**) The viscosity sensitivity of **P1** was determined to be 0.20 from the slope of the linear plot of the logarithm of the fluorescence emission intensity of **P1** at 633 nm as a function of logarithm of viscosity. (**e**) Structure of **P2**. (**f**) Fluorescence excitation spectrum of **P2** (5 μM) in glycerol. (**g**) Fluorescence emission spectrum of **P2** (5 μM) in glycerol. (**h**) The viscosity sensitivity of **P2** was determined to be 0.54 from the slope of the linear plot of the logarithm of the fluorescence emission intensity of **P2** at 524 nm as a function of logarithm of viscosity. All fluorescence was measured by using a Tecan infinite M1000Pro fluorescence microplate reader at room temperature.

**Figure S10.**
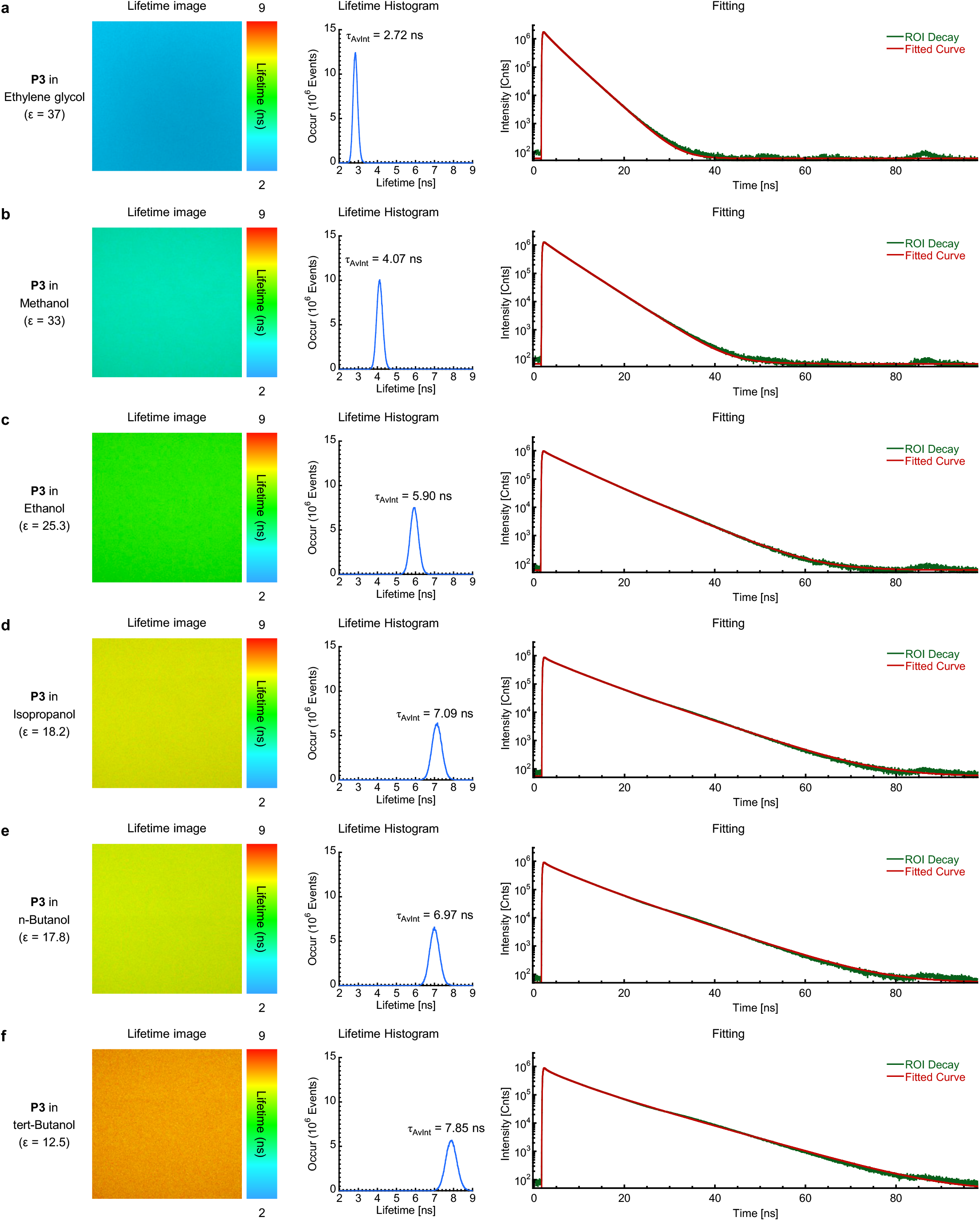
Average fluorescence lifetime (τ_AvInt_) of P3 increases as the dielectric constant (ε) of solvent tested decreases. The fluorescence lifetime images (left panel), lifetime histograms (center panel), and lifetime decay curves (right panel) of **P3** in various polar protic solvents with different dielectric constant values. (**a**) in ethylene glycol, (**b**) in methanol, (**c**) in ethanol, (**d**) isopropanol, (**e**) n-butanol, (**f**) tert-butanol.

**Figure S11.**
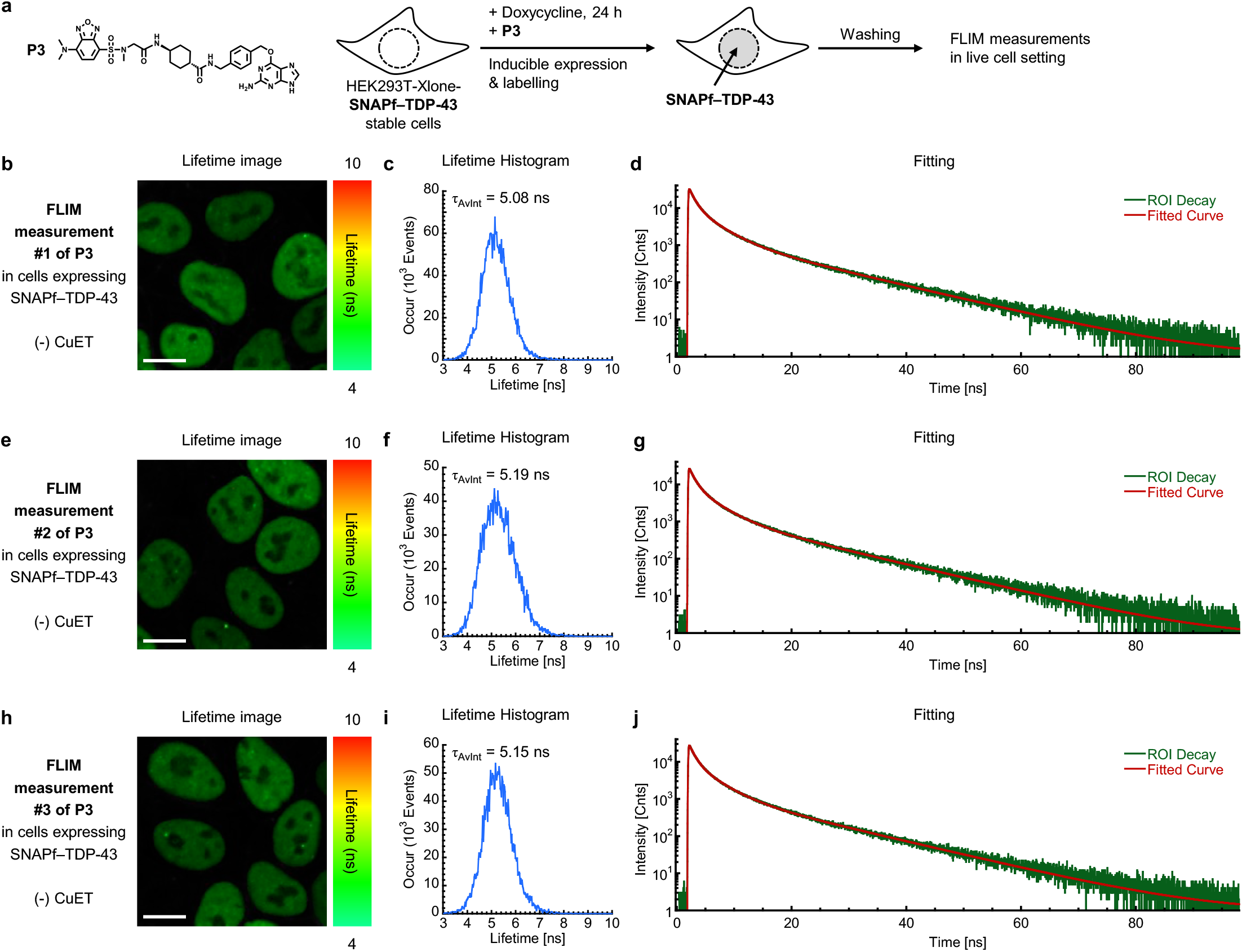
FLIM measurements of P3 with live cells stably expressing SNAPf–TDP-43 without CuET treatment. (**a**) The stable cells were treated with doxycycline (140 ng/mL) and **P3** (0.5 μM) for 24 hours for the inducible expression and labeling of SNAPf–TDP-43, respectively. After the cells were washed with fresh fluorobrite^TM^ DMEM media supplemented with fetal bovine serum (10%) to remove excess probes, the lifetime of **P3** was measured using Zeiss LSM 880 microscope with a PicoQuant-FLIM LSM upgrade KIT. (**b**), (**e**), (**h**) lifetime images from three individual measurements. (**c**), (**f**), (**i**) lifetime histograms from the images b, e, and h, respectively. (**d**), (**g**), (**j**) lifetime decay fitting from the images b, e, and h, respectively. Scale bar = 10 µm.

**Figure S12.**
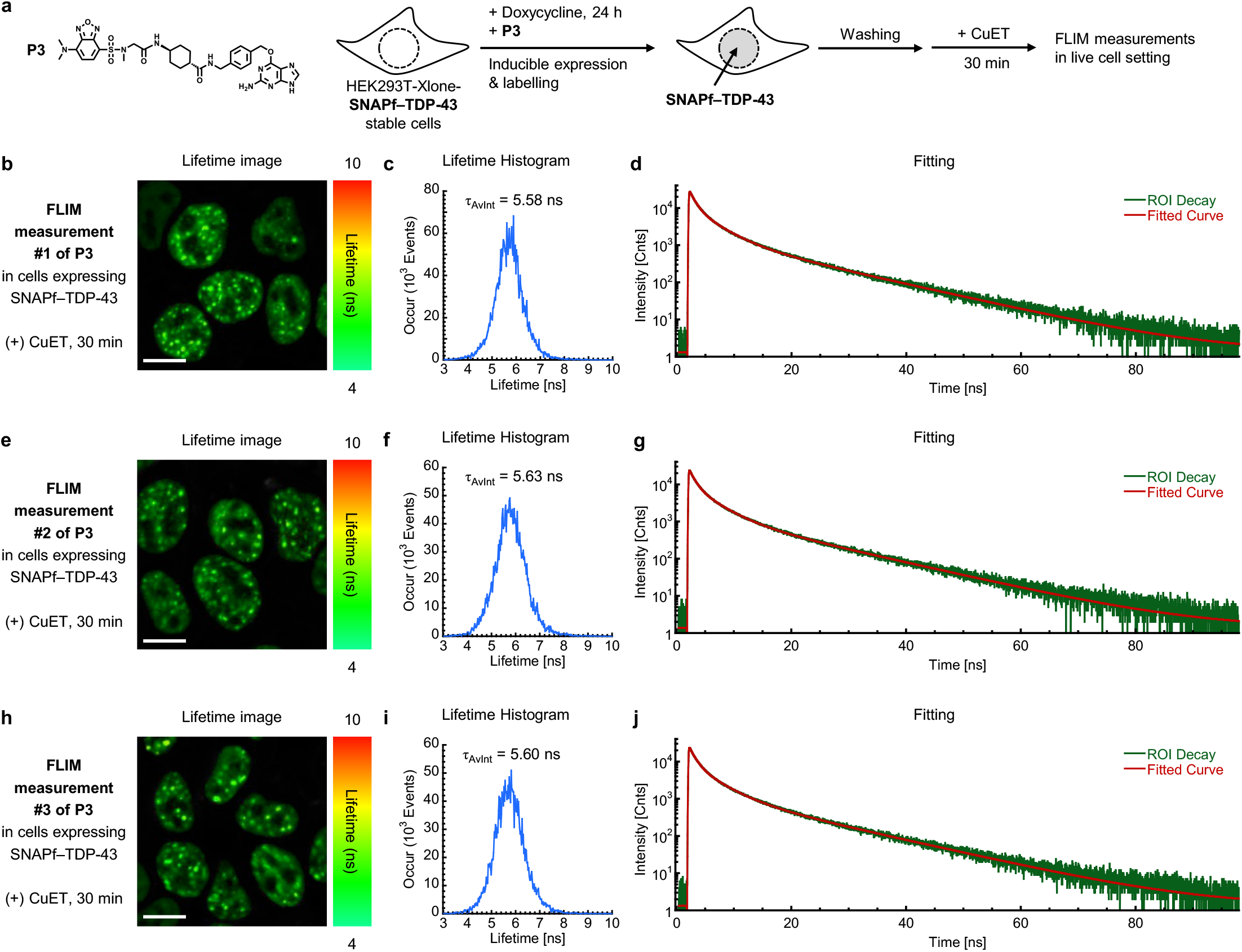
FLIM measurements of P3 with live cells stably expressing SNAPf–TDP-43 upon CuET treatment 30 minutes. (**a**) The stable cells were treated with doxycycline (140 ng/mL) and **P3** (0.5 μM) for 24 hours for the inducible expression and labeling of SNAPf–TDP-43, respectively. After the cells were washed with fresh fluorobrite^TM^ DMEM media supplemented with fetal bovine serum (10%) to remove excess probes, the cells were treated with CuET for 30 minutes at 37 °C under CO_2_ (5%) and subsequently the lifetime of **P3** was measured using Zeiss LSM 880 microscope with a PicoQuant-FLIM LSM upgrade KIT. (**b**), (**e**), (**h**) lifetime images from three individual measurements. (**c**), (**f**), (**i**) lifetime histograms from the images b, e, and h, respectively. (**d**), (**g**), (**j**) lifetime decay fitting from the images b, e, and h, respectively. Scale bar = 10 µm.

**Figure S13.**
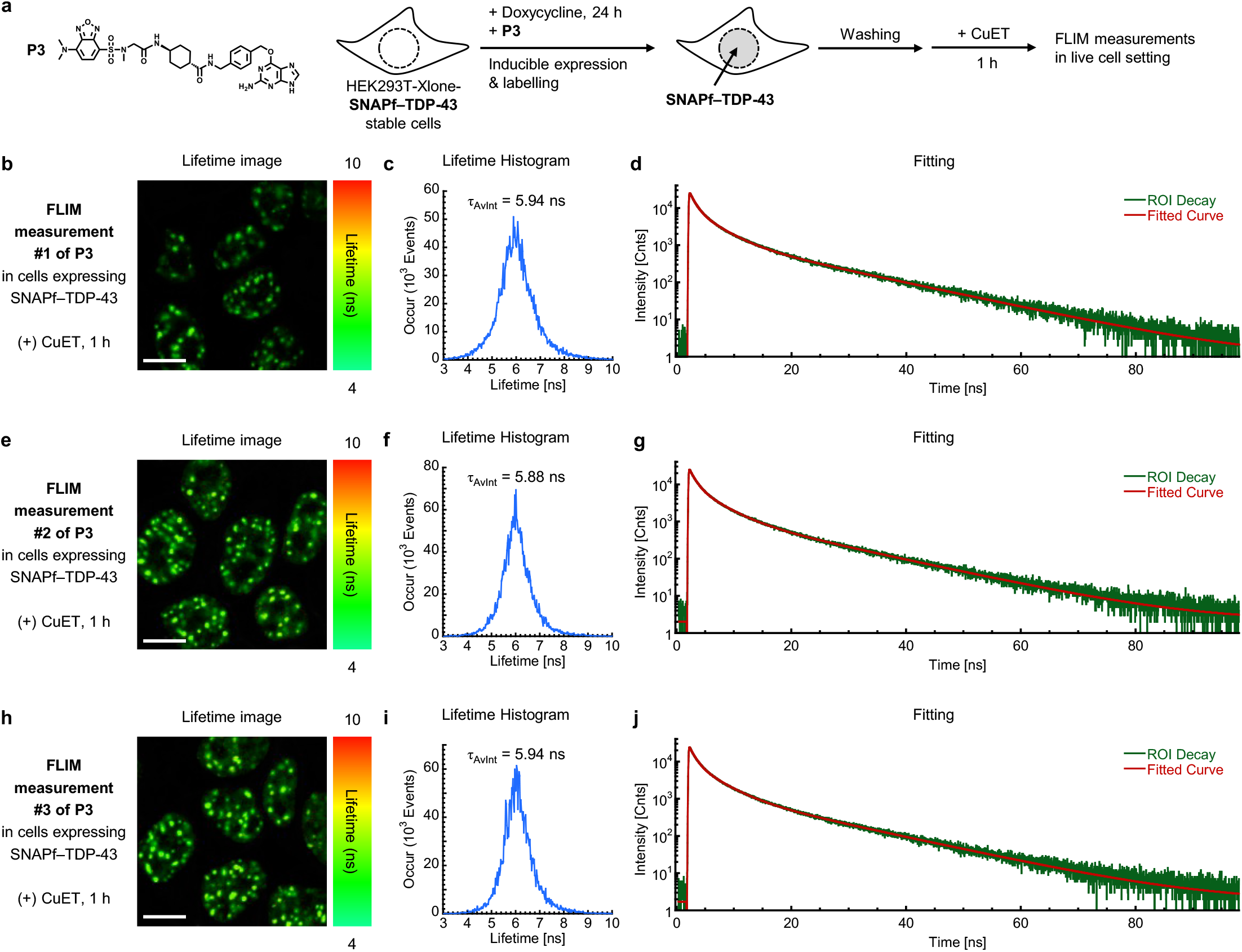
FLIM measurements of P3 with live cells stably expressing SNAPf–TDP-43 upon CuET treatment for 1 hour. (**a**) The stable cells were treated with doxycycline (140 ng/mL) and **P3** (0.5 μM) for 24 hours for the inducible expression and labeling of SNAPf–TDP-43, respectively. After the cells were washed with fresh fluorobrite^TM^ DMEM media supplemented with fetal bovine serum (10%) to remove excess probes, the cells were treated with CuET for 1 hour at 37 °C under CO_2_ (5%) and subsequently the lifetime of **P3** was measured using Zeiss LSM 880 microscope with a PicoQuant-FLIM LSM upgrade KIT. (**b**), (**e**), (**h**) lifetime images from three individual measurements. (**c**), (**f**), (**i**) lifetime histograms from the images b, e, and h, respectively. (**d**), (**g**), (**j**) lifetime decay fitting from the images b, e, and h, respectively. Scale bar = 10 µm.

**Figure S14.**
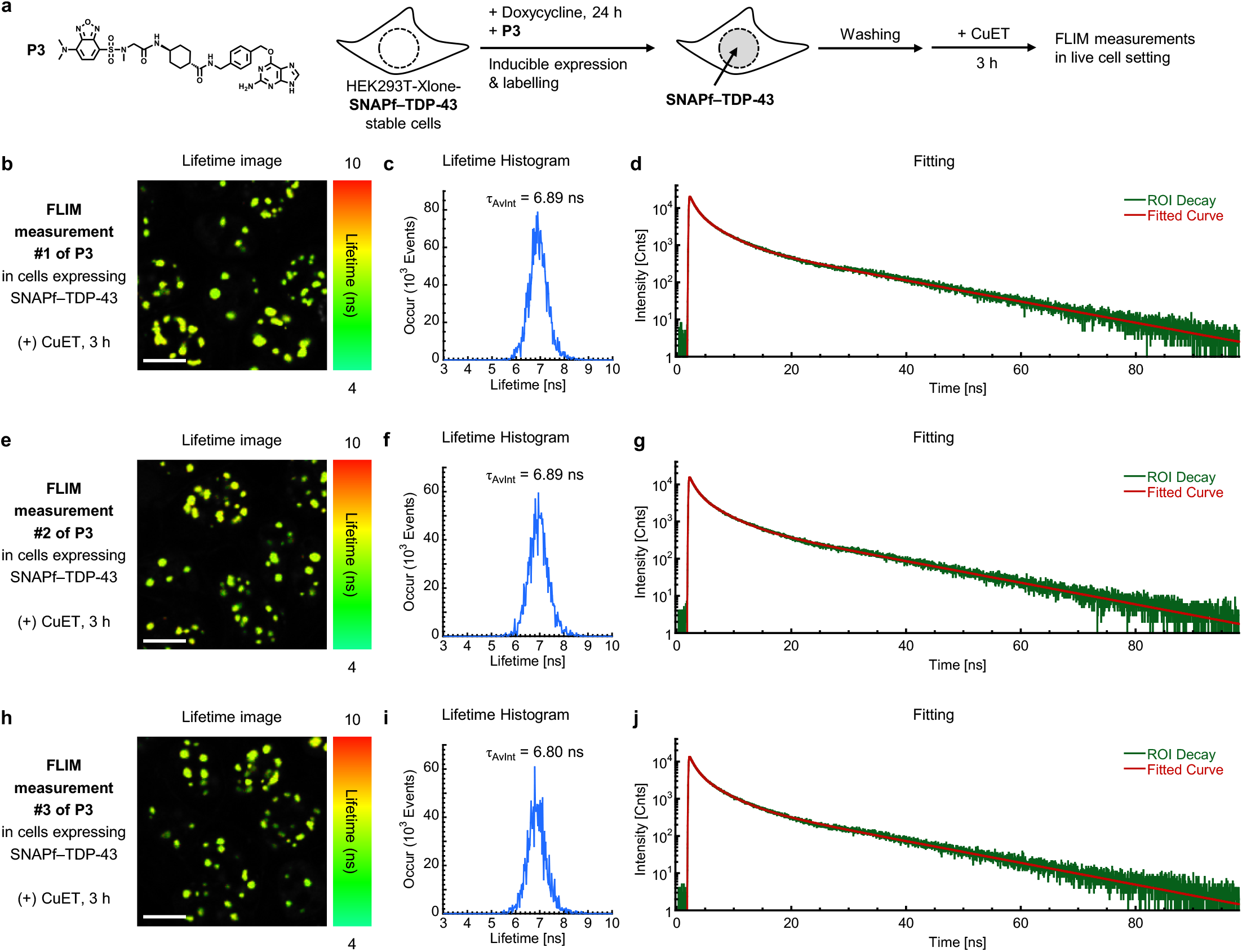
FLIM measurements of P3 with live cells stably expressing SNAPf–TDP-43 upon CuET treatment for 3 hours. The stable cells were treated with doxycycline (140 ng/mL) and **P3** (0.5 μM) for 24 hours for the inducible expression and labeling of SNAPf–TDP-43, respectively. After the cells were washed with fresh fluorobrite^TM^ DMEM media supplemented with fetal bovine serum (10%) to remove excess probes, the cells were treated with CuET for 3 hours at 37 °C under CO_2_ (5%) and subsequently the lifetime of **P3** was measured using Zeiss LSM 880 microscope with a PicoQuant-FLIM LSM upgrade KIT. (**b**), (**e**), (**h**) lifetime images from three individual measurements. (**c**), (**f**), (**i**) lifetime histograms from the images b, e, and h, respectively. (**d**), (**g**), (**j**) lifetime decay fitting from the images b, e, and h, respectively. Scale bar = 10 µm.

**Figure S15.**
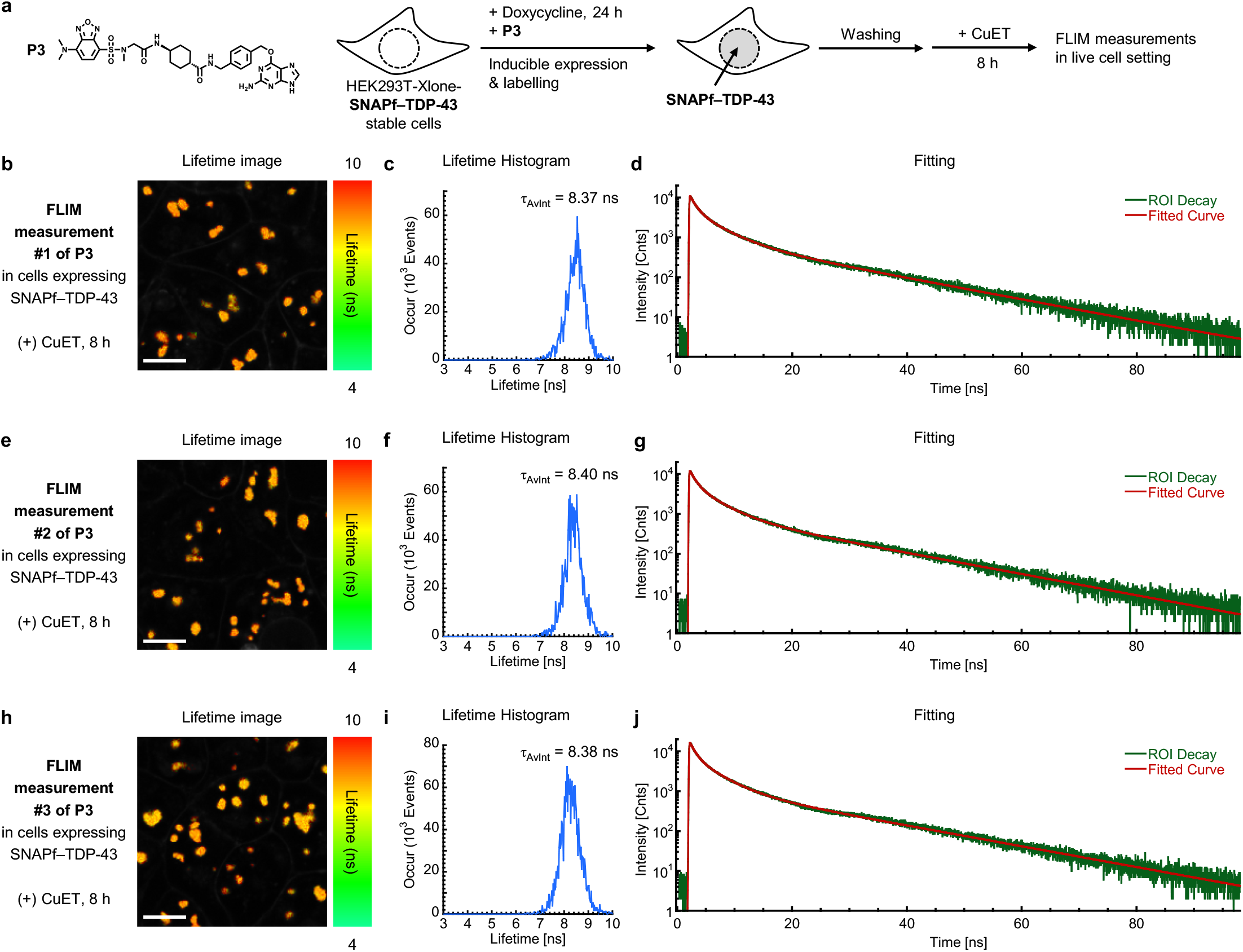
FLIM measurements of P3 with live cells stably expressing SNAPf–TDP-43 upon CuET treatment for 8 hours. The stable cells were treated with doxycycline (140 ng/mL) and **P3** (0.5 μM) for 24 hours for the inducible expression and labeling of SNAPf–TDP-43, respectively. After the cells were washed with fresh fluorobrite^TM^ DMEM media supplemented with fetal bovine serum (10%) to remove excess probes, the cells were treated with CuET for 8 hours at 37 °C under CO_2_ (5%) and subsequently the lifetime of **P3** was measured using Zeiss LSM 880 microscope with a PicoQuant-FLIM LSM upgrade KIT. (**b**), (**e**), (**h**) lifetime images from three individual measurements. (**c**), (**f**), (**i**) lifetime histograms from the images b, e, and h, respectively. (**d**), (**g**), (**j**) lifetime decay fitting from the images b, e, and h, respectively. Scale bar = 10 µm.

**Figure S16.**
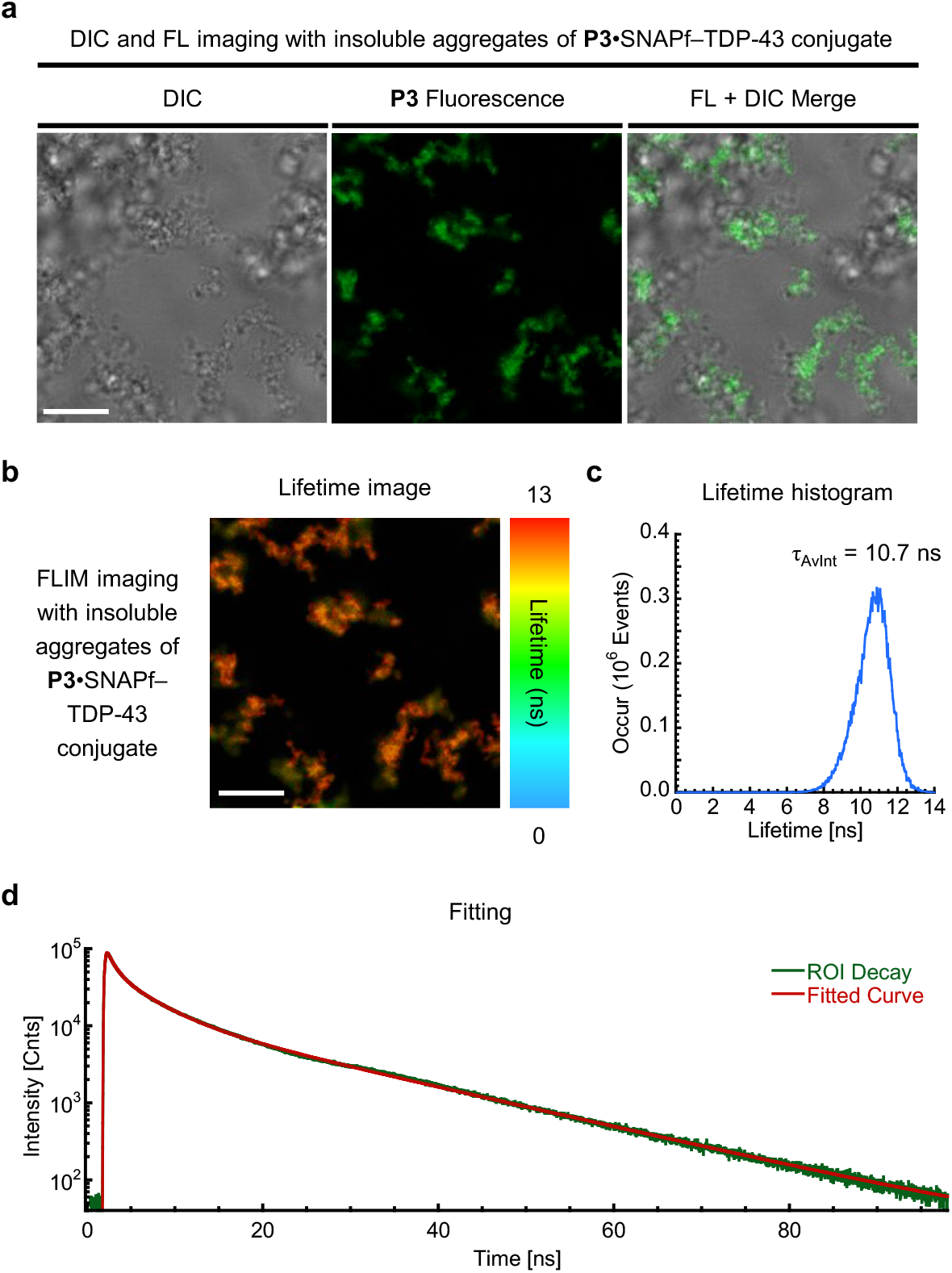
Confocal and FLIM imaging experiments with insoluble aggregates of P3•SNAPf–TDP-43 conjugate. (**a**) DIC and FL imaging, (**b**) lifetime imaging, (**c**) lifetime histogram, (**d**) lifetime decay fitting with insoluble aggregates of **P3**•SNAPf–TDP-43 conjugate. Scale bar = 10 µm. The insoluble aggregates of **P3**•SNAPf–TDP-43 conjugate were generated from recombinantly purified SNAPf–TDP-43–TEV–Halo fusion protein. The protein was constructed and purified from *E. Coli* according to the previously reported procedures.^73^ For labeling, the fusion protein (10 µM) was incubated with **P3** (5 µM) at 37 °C for 30 minutes in HEPES buffer (20 mM, pH 7.5, 140 mM NaCl) containing DTT (1 mM). After labeling, the protein conjugate solution was treated with TEV protease (0.75 µM) and PEG3350 (5%) at room temperature for 1 hour to induce the insoluble aggregates.

**Figure S17.**
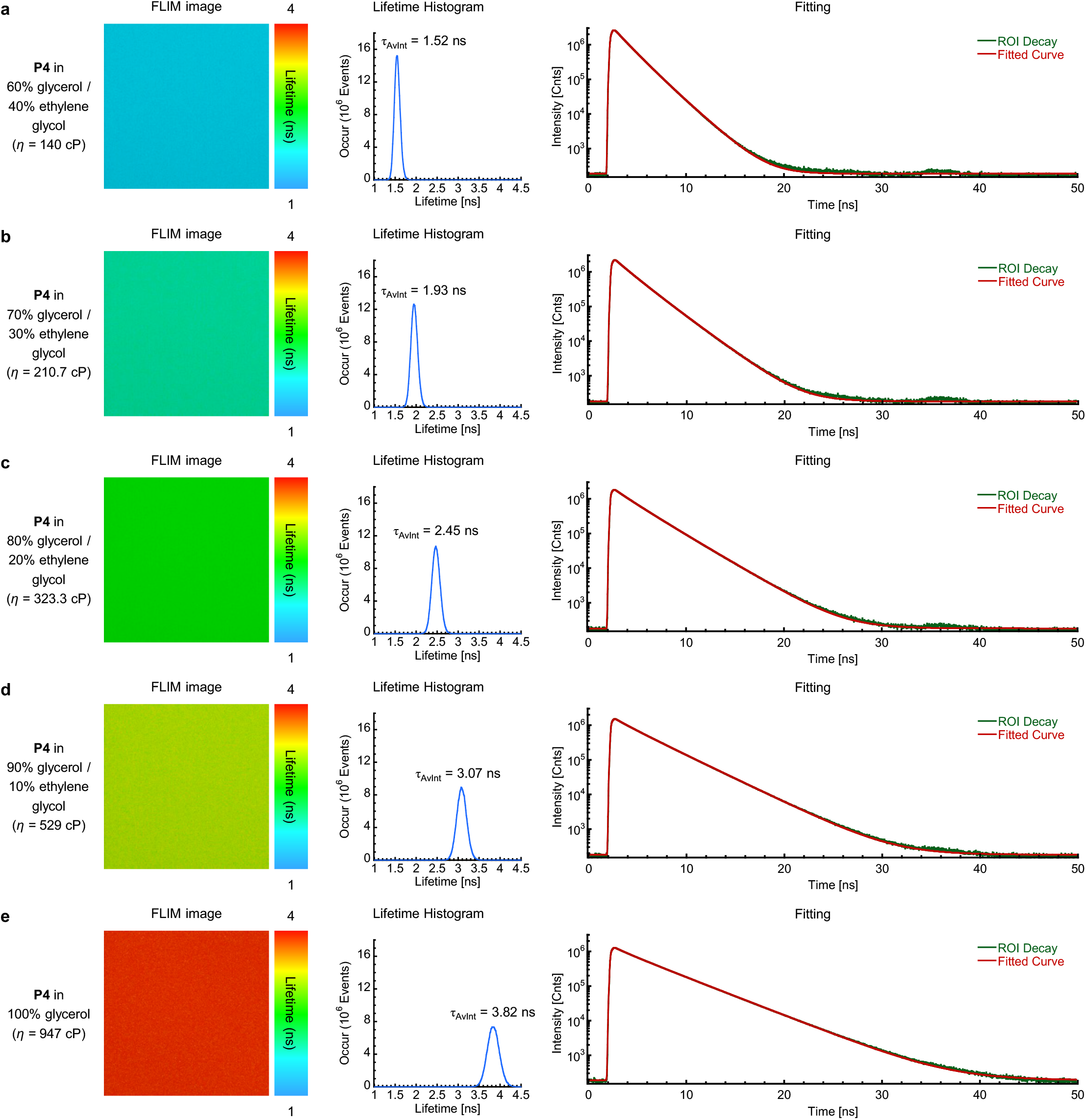
Average fluorescence lifetime (τ_AvInt_) of P4 increases as the viscosity (η) of solvent mixtures tested increases. The fluorescence lifetime images (left panel), lifetime histograms (center panel), and lifetime decay curves (right panel) of **P4** in mixtures of glycerol and ethylene glycol with different volume fractions (v/v, %). (**a**) in 60% glycerol and 40% ethylene glycol, (**b**) in 70% glycerol and 30% ethylene glycol, (**c**) in 80% glycerol and 20% ethylene glycol, (**d**) in 90% glycerol and 10% ethylene glycol, (**e**) in 100% glycerol.

**Figure S18.**
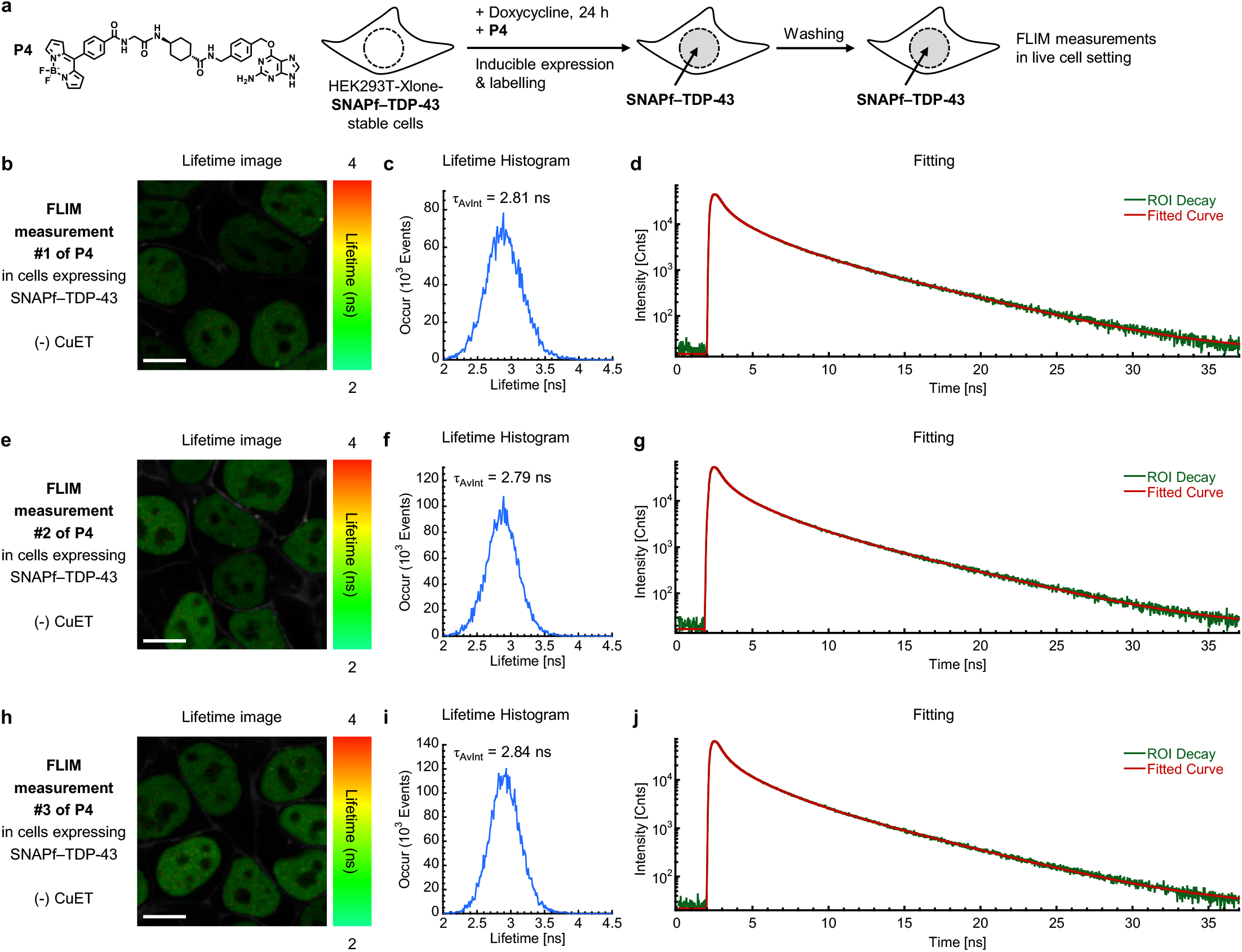
FLIM measurements of P4 with live cells stably expressing SNAPf–TDP-43 without CuET treatment. (**a**) The stable cells were treated with doxycycline (140 ng/mL) and **P4** (0.5 μM) for 24 hours for the inducible expression and labeling of SNAPf–TDP-43, respectively. After the cells were washed with fresh fluorobrite^TM^ DMEM media supplemented with fetal bovine serum (10%) to remove excess probes, the lifetime of **P4** was measured using Zeiss LSM 880 microscope with a PicoQuant-FLIM LSM upgrade KIT. (**b**), (**e**), (**h**) lifetime images from three individual measurements, (**c**), (**f**), (**i**) lifetime histograms from the images b, e, and h, respectively, (**d**), (**g**), (**j**) lifetime decay fitting from the images b, e, and h, respectively. Scale bar = 10 µm.

**Figure S19.**
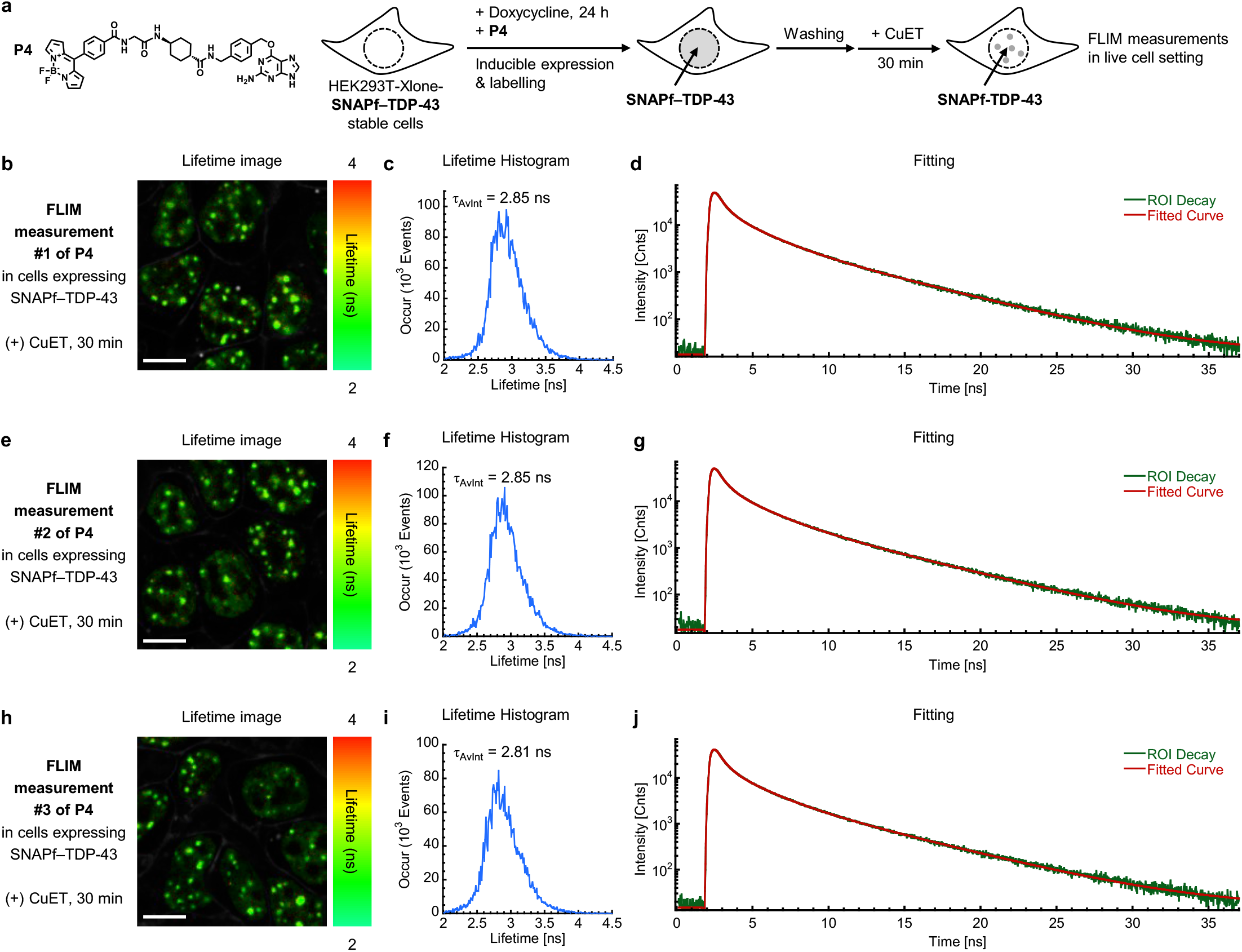
FLIM measurements of P4 with live cells stably expressing SNAPf–TDP-43 upon CuET treatment for 30 minutes. (**a**) The stable cells were treated with doxycycline (140 ng/mL) and **P4** (0.5 μM) for 24 hours for the inducible expression and labeling of SNAPf–TDP-43, respectively. After the cells were washed with fresh fluorobrite^TM^ DMEM media supplemented with fetal bovine serum (10%) to remove excess probes, the cells were treated with CuET for 30 minutes at 37 °C under CO_2_ (5%) and subsequently the lifetime of **P4** was measured using Zeiss LSM 880 microscope with a PicoQuant-FLIM LSM upgrade KIT. (**b**), (**e**), (**h**) lifetime images from three individual measurements. (**c**), (**f**), (**i**) lifetime histograms from the images b, e, and h, respectively. (**d**), (**g**), (**j**) lifetime decay fitting from the images b, e, and h, respectively. Scale bar = 10 µm.

**Figure S20.**
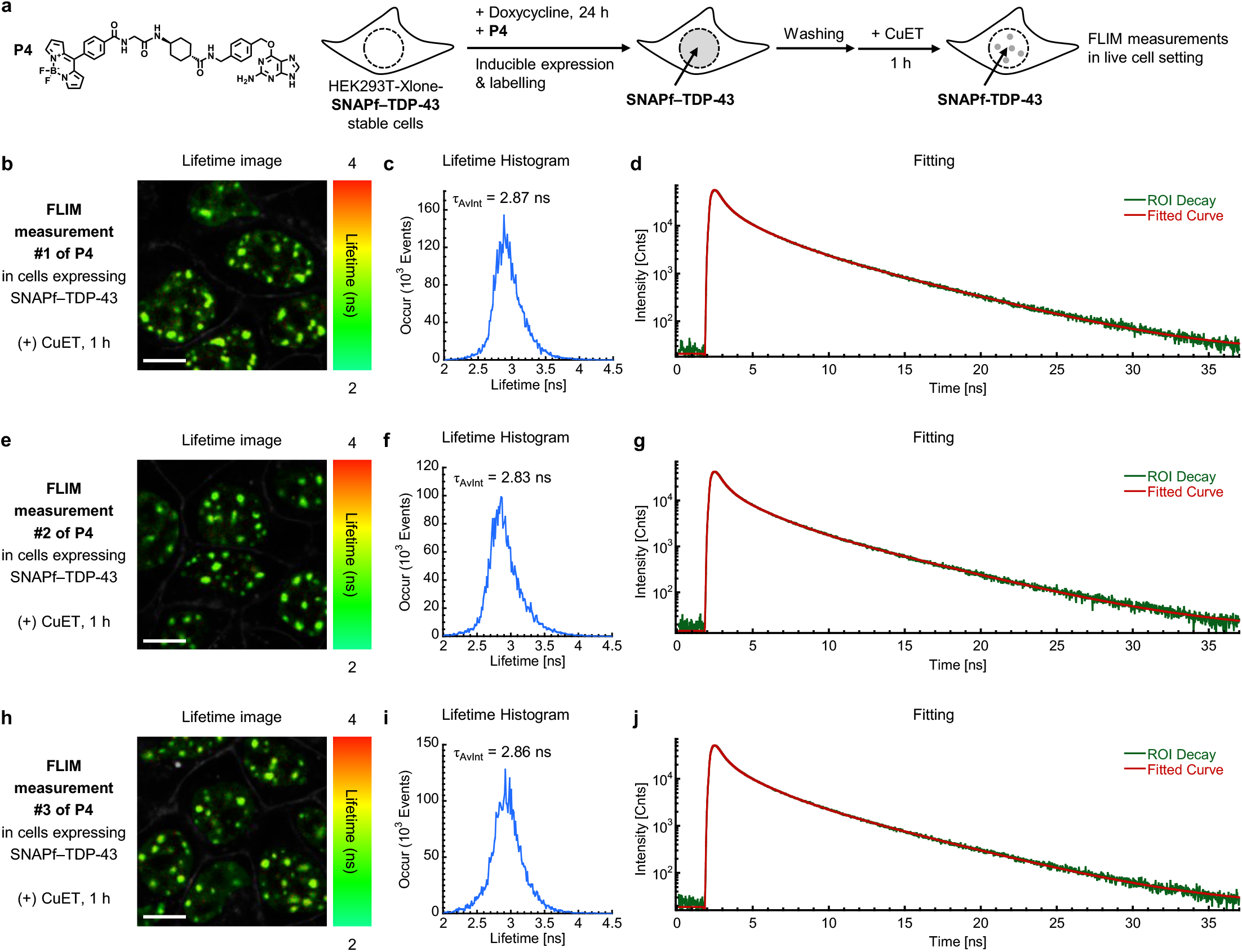
FLIM measurements of P4 with live cells stably expressing SNAPf–TDP-43 upon CuET treatment for 1 hour. (**a**) The stable cells were treated with doxycycline (140 ng/mL) and **P4** (0.5 μM) for 24 hours for the inducible expression and labeling of SNAPf–TDP-43, respectively. After the cells were washed with fresh fluorobrite^TM^ DMEM media supplemented with fetal bovine serum (10%) to remove excess probes, the cells were treated with CuET for 1 hour at 37 °C under CO_2_ (5%) and subsequently the lifetime of **P4** was measured using Zeiss LSM 880 microscope with a PicoQuant-FLIM LSM upgrade KIT. (**b**), (**e**), (**h**) lifetime images from three individual measurements. (**c**), (**f**), (**i**) lifetime histograms from the images b, e, and h, respectively. (**d**), (**g**), (**j**) lifetime decay fitting from the images b, e, and h, respectively. Scale bar = 10 µm.

**Figure S21.**
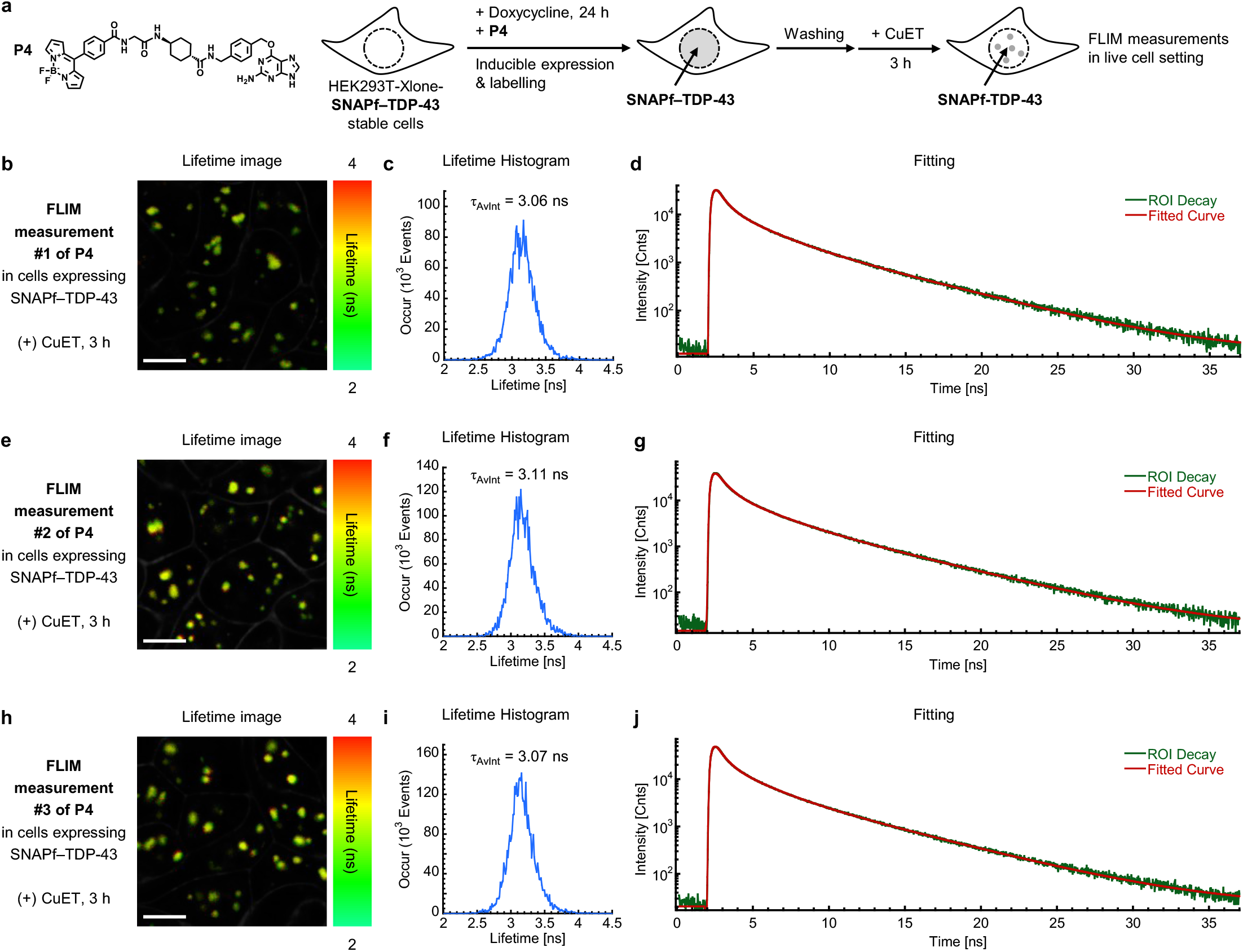
FLIM measurements of P4 with live cells stably expressing SNAPf–TDP-43 upon CuET treatment for 3 hours. (**a**) The stable cells were treated with doxycycline (140 ng/mL) and **P4** (0.5 μM) for 24 hours for the inducible expression and labeling of SNAPf–TDP-43, respectively. After the cells were washed with fresh fluorobrite^TM^ DMEM media supplemented with fetal bovine serum (10%) to remove excess probes, the cells were treated with CuET for 3 hours at 37 °C under CO_2_ (5%) and subsequently the lifetime of **P4** was measured using Zeiss LSM 880 microscope with a PicoQuant-FLIM LSM upgrade KIT. (**b**), (**e**), (**h**) lifetime images from three individual measurements. (**c**), (**f**), (**i**) lifetime histograms from the images b, e, and h, respectively. (**d**), (**g**),(**j**) lifetime decay fitting from the images b, e, and h, respectively. Scale bar = 10 µm.

**Figure S22.**
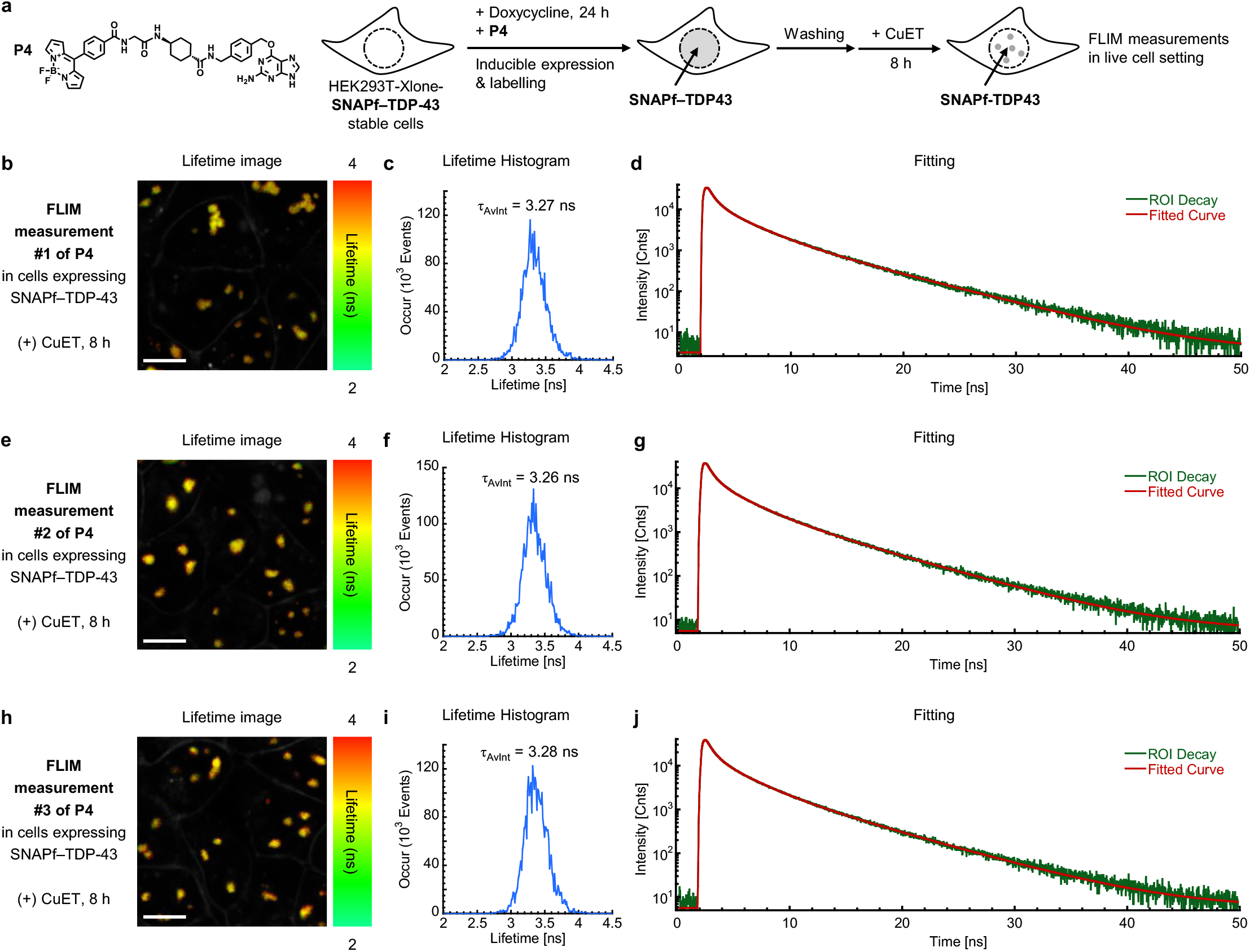
FLIM measurements of P4 with live cells stably expressing SNAPf–TDP-43 upon CuET treatment for 8 hours. (**a**) The stable cells were treated with doxycycline (140 ng/mL) and **P4** (0.5 μM) for 24 hours for the inducible expression and labeling of SNAPf–TDP-43, respectively. After the cells were washed with fresh fluorobrite^TM^ DMEM media supplemented with fetal bovine serum (10%) to remove excess probes, the cells were treated with CuET for 8 hours at 37 °C under CO_2_ (5%) and subsequently the lifetime of **P4** was measured using Zeiss LSM 880 microscope with a PicoQuant-FLIM LSM upgrade KIT. (**b**), (**e**), (**h**) lifetime images from three individual measurements. (**c**), (**f**), (**i**) lifetime histograms from the images b, e, and h, respectively. (**d**), (**g**), (**j**) lifetime decay fitting from the images b, e, and h, respectively. Scale bar = 10 µm.

**Figure S23.**
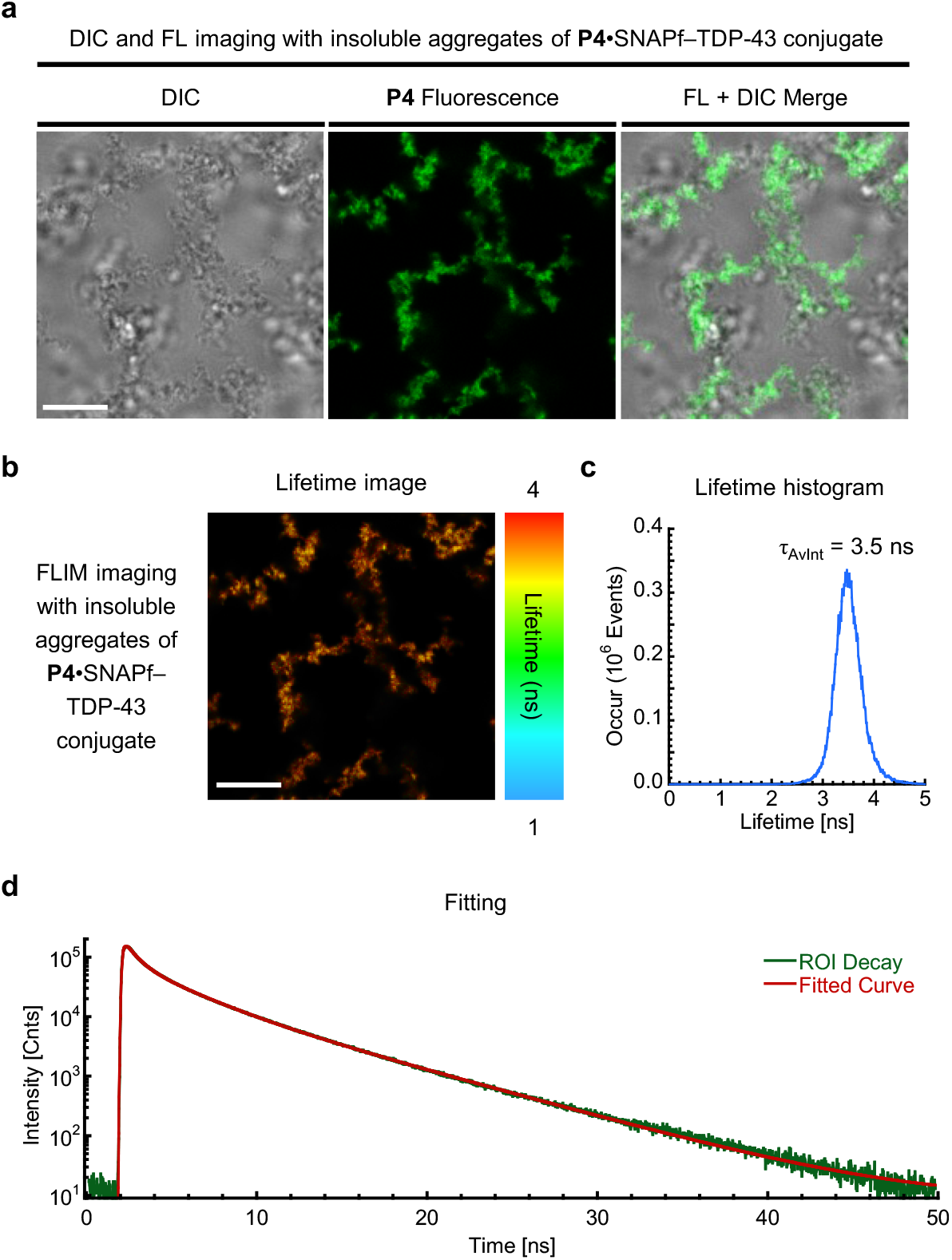
Confocal and FLIM imaging experiments with insoluble aggregates of P4•SNAPf–TDP-43 conjugate. (**a**) DIC and FL imaging, (**b**) lifetime imaging, (**c**) lifetime histogram, (**d**) lifetime decay fitting with insoluble aggregates of **P4**•SNAPf–TDP-43 conjugate. Scale bar = 10 µm. The insoluble aggregates of **P4**•SNAPf–TDP-43 conjugate were generated from recombinantly purified SNAPf–TDP-43–TEV–Halo fusion protein. The protein was constructed and purified from *E. Coli* according to the previously reported procedures.^73^ For labeling, the fusion protein (10 µM) was incubated with **P4** (5 µM) at 37 °C for 30 minutes in HEPES buffer (20 mM, pH 7.5, 140 mM NaCl) containing DTT (1 mM). After labeling, the protein conjugate solution was treated with TEV protease (0.75 µM) and PEG3350 (5%) at room temperature for 1 hour to induce the insoluble aggregates.

**Figure S24.**
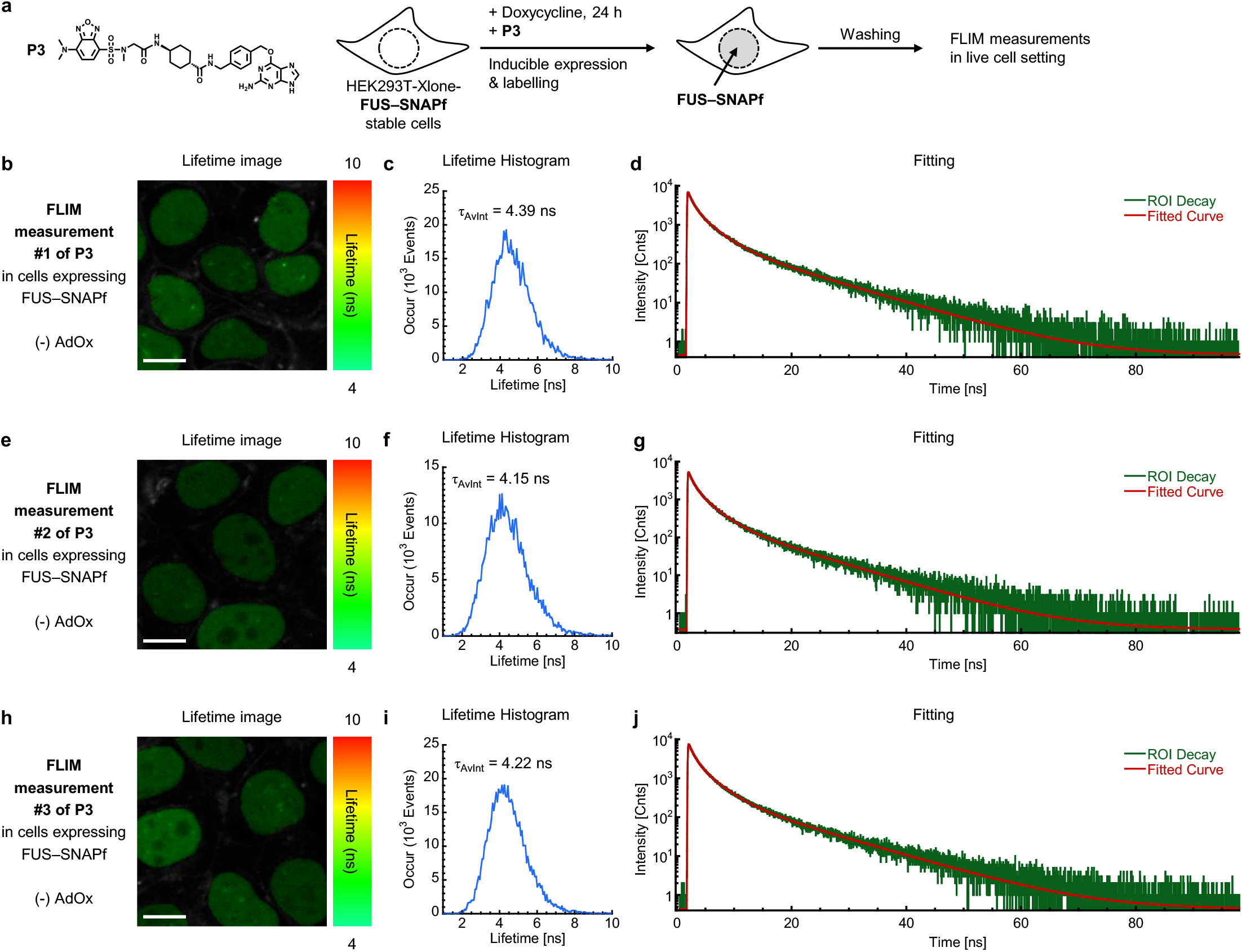
FLIM measurements of P3 with live cells stably expressing FUS–SNAPf without AdOx pretreatment. (**a**) The stable cells were treated with doxycycline (140 ng/mL) and **P3** (0.5 μM) for 24 hours for the inducible expression and labeling of FUS–SNAPf, respectively. After the cells were washed with fresh fluorobrite^TM^ DMEM media supplemented with fetal bovine serum (10%) to remove excess probes, the lifetime of **P3** was measured using Zeiss LSM 880 microscope with a PicoQuant-FLIM LSM upgrade KIT. (**b**), (**e**), (**h**) lifetime images from three individual measurements. (**c**), (**f**), (**i**) lifetime histograms from the images b, e, and h, respectively. (**d**), (**g**), (**j**) lifetime decay fitting from the images b, e, and h, respectively. Scale bar = 10 µm.

**Figure S25.**
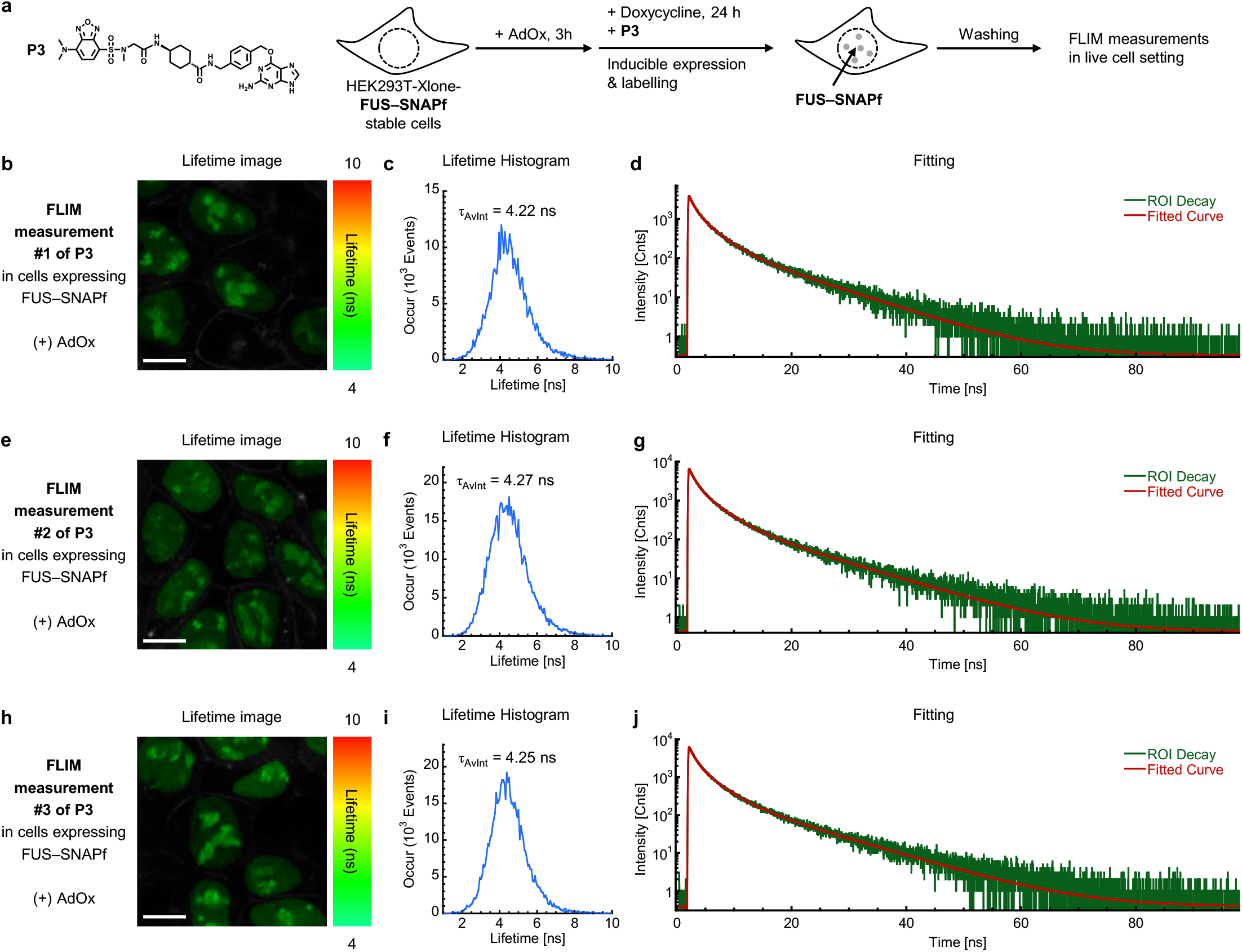
FLIM measurements of P3 with live cells stably expressing FUS–SNAPf with AdOx pretreatment. (**a**) The stable cells were pretreated with AdOx (25 μM) 3 hours before they were further treated with doxycycline (140 ng/mL) and **P3** (0.5 μM) for 24 hours for the inducible expression and labeling of FUS– SNAPf, respectively. After the cells were washed with fresh fluorobrite^TM^ DMEM media supplemented with fetal bovine serum (10%) to remove excess probes, the lifetime of **P3** was measured using Zeiss LSM 880 microscope with a PicoQuant-FLIM LSM upgrade KIT. (**b**), (**e**), (**h**) lifetime images from three individual measurements. (**c**), (**f**), (**i**) lifetime histograms from the images b, e, and h, respectively. (**d**), (**g**), (**j**) lifetime decay fitting from the images b, e, and h, respectively. Scale bar = 10 µm.

**Figure S26.**
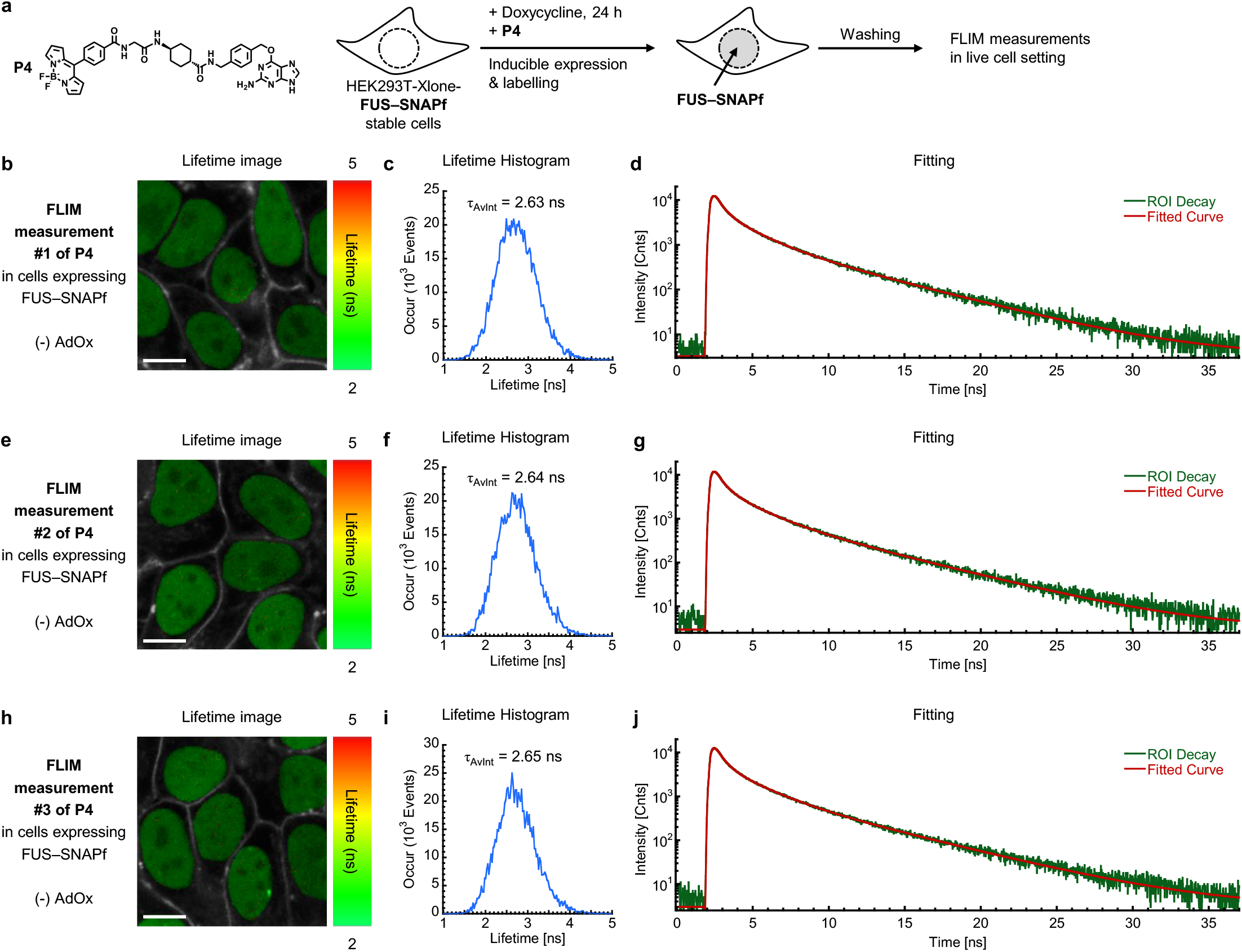
FLIM measurements of P4 with live cells stably expressing FUS–SNAPf without AdOx pretreatment. (**a**) The stable cells were treated with doxycycline (140 ng/mL) and **P4** (0.5 μM) for 24 hours for the inducible expression and labeling of FUS–SNAPf, respectively. After the cells were washed with fresh fluorobrite^TM^ DMEM media supplemented with fetal bovine serum (10%) to remove excess probes, the lifetime of **P4** was measured using Zeiss LSM 880 microscope with a PicoQuant-FLIM LSM upgrade KIT. (**b**), (**e**), (**h**) lifetime images from three individual measurements. (**c**), (**f**), (**i**) lifetime histograms from the images b, e, and h, respectively. (**d**), (**g**), (**j**) lifetime decay fitting from the images b, e, and h, respectively. Scale bar = 10 µm.

**Figure S27.**
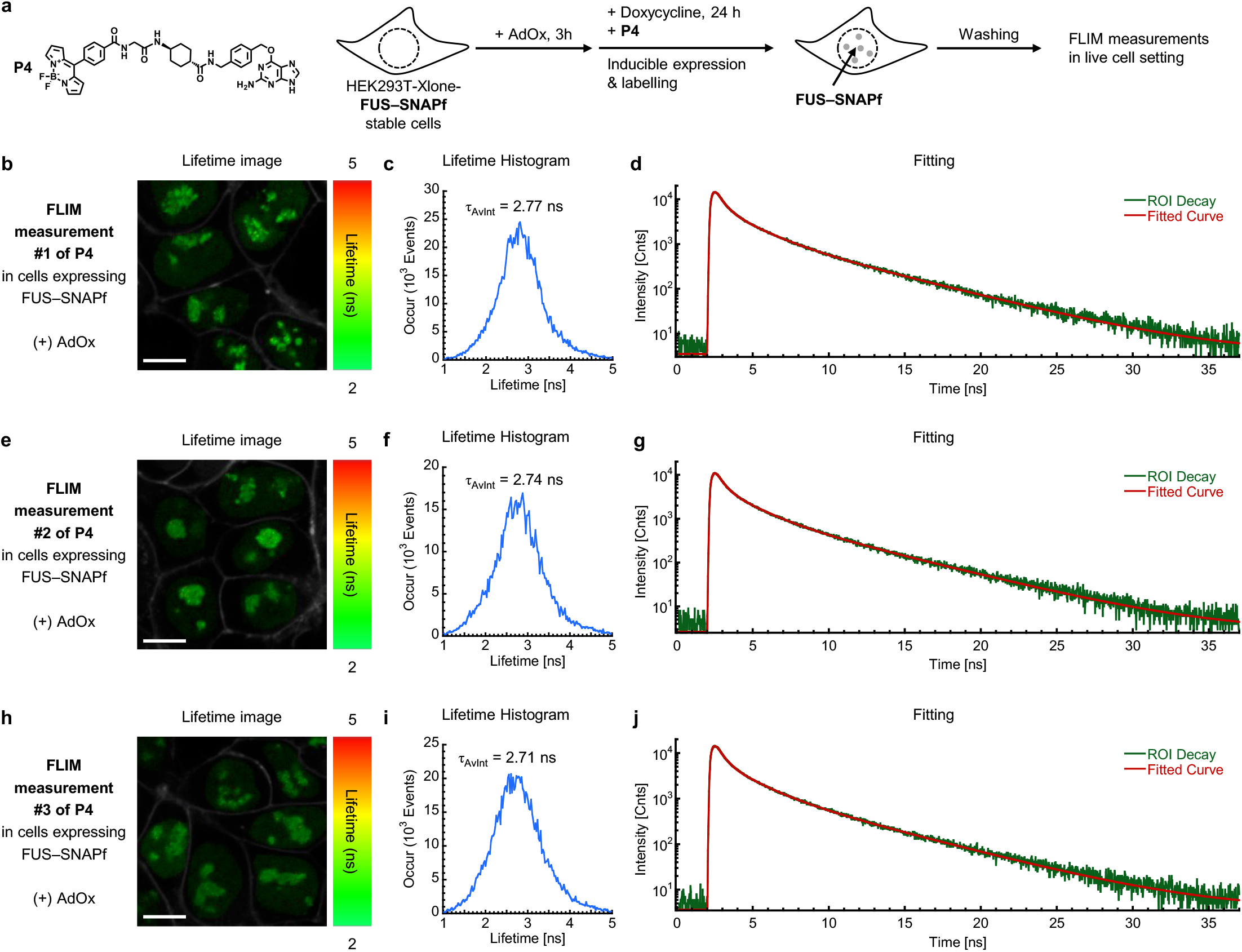
FLIM measurements of P4 with live cells stably expressing FUS–SNAPf with AdOx pretreatment. (**a**) The stable cells were pretreated with AdOx (25 μM) 3 hours before they were further treated with doxycycline (140 ng/mL) and **P4** (0.5 μM) for 24 hours for the inducible expression and labeling of FUS– SNAPf, respectively. After the cells were washed with fresh fluorobrite^TM^ DMEM media supplemented with fetal bovine serum (10%) to remove excess probes, the lifetime of **P4** was measured using Zeiss LSM 880 microscope with a PicoQuant-FLIM LSM upgrade KIT. (**b**), (**e**), (**h**) lifetime images from three individual measurements. (**c**), (**f**), (**i**) lifetime histograms from the images b, e, and h, respectively. (**d**), (**g**), (**j**) lifetime decay fitting from the images b, e, and h, respectively. Scale bar = 10 µm.

**Figure S28.**
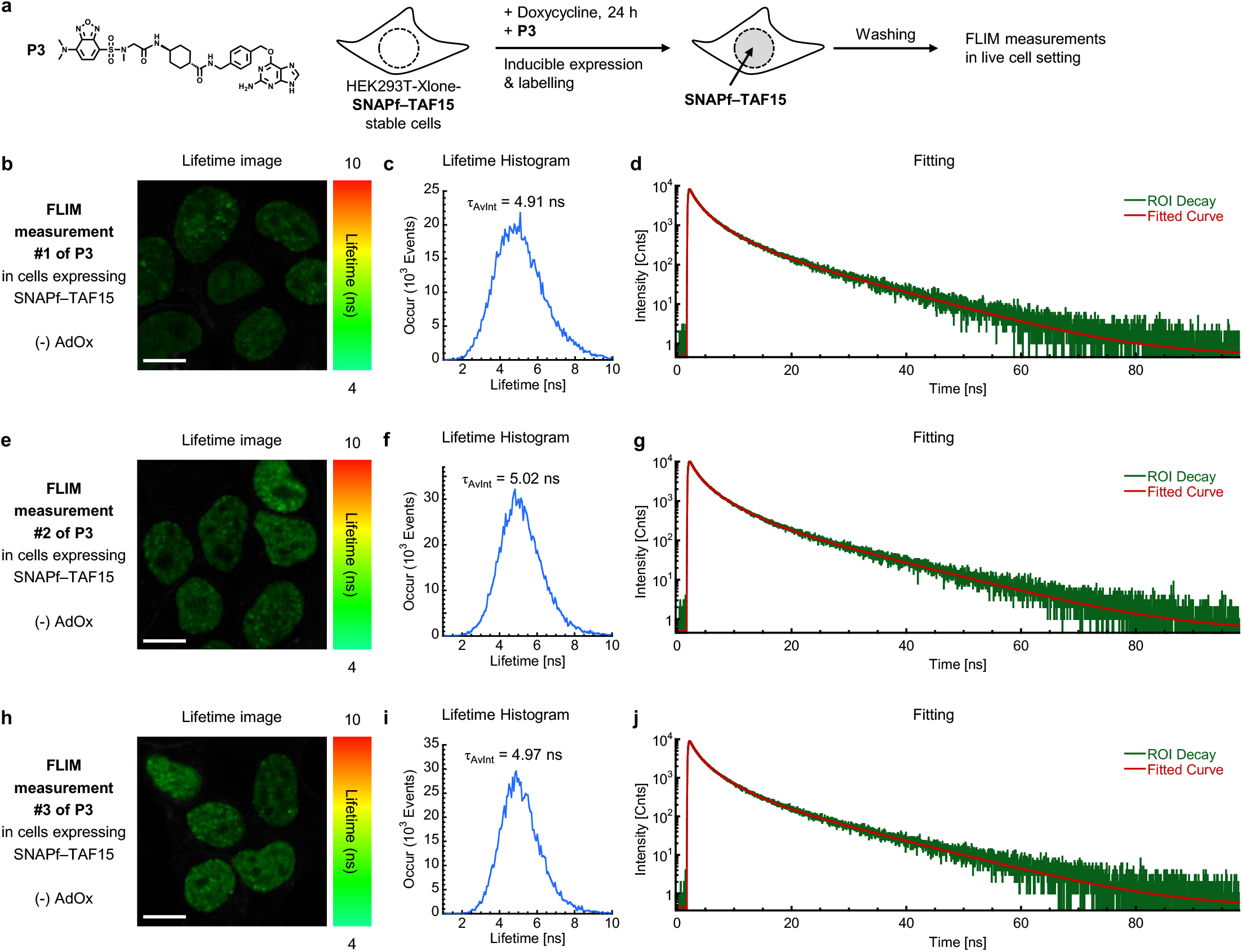
FLIM measurements of P3 with live cells stably expressing SNAPf–TAF15 without AdOx pretreatment. (**a**) The stable cells were treated with doxycycline (140 ng/mL) and **P3** (0.5 μM) for 24 hours for the inducible expression and labeling of SNAPf–TAF15, respectively. After the cells were washed with fresh fluorobrite^TM^ DMEM media supplemented with fetal bovine serum (10%) to remove excess probes, the lifetime of **P3** was measured using Zeiss LSM 880 microscope with a PicoQuant-FLIM LSM upgrade KIT. (**b**), (**e**), (**h**) lifetime images from three individual measurements. (**c**), (**f**), (**i**) lifetime histograms from the images b, e, and h, respectively. (**d**), (**g**), (**j**) lifetime decay fitting from the images b, e, and h, respectively. Scale bar = 10 µm.

**Figure S29.**
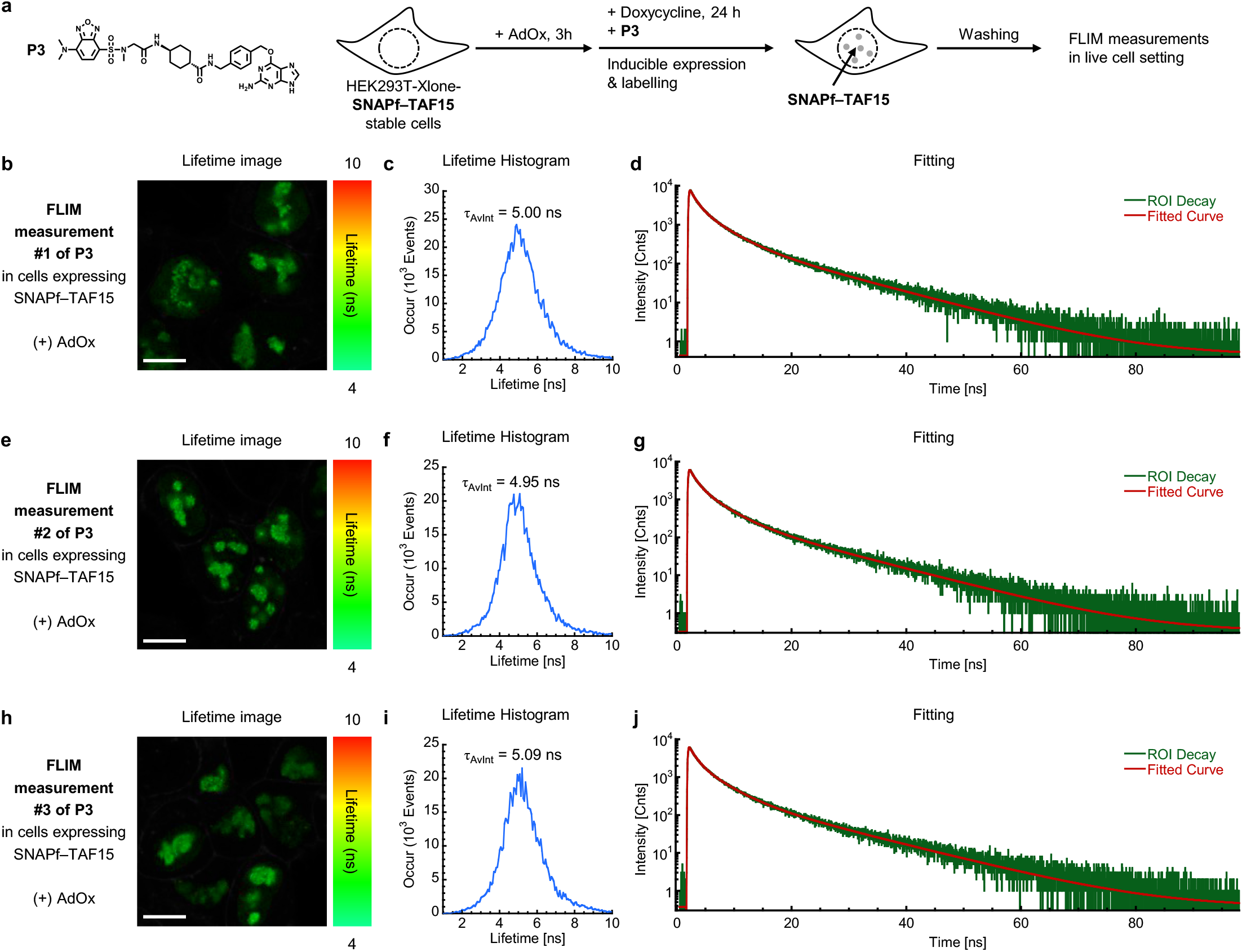
FLIM measurements of P3 with live cells stably expressing SNAPf–TAF15 with AdOx pretreatment. (**a**) The stable cells were pretreated with AdOx (25 μM) 3 hours before they were further treated with doxycycline (140 ng/mL) and **P3** (0.5 μM) for 24 hours for the inducible expression and labeling of SNAPf– TAF15, respectively. After the cells were washed with fresh fluorobrite^TM^ DMEM media supplemented with fetal bovine serum (10%) to remove excess probes, the lifetime of **P3** was measured using Zeiss LSM 880 microscope with a PicoQuant-FLIM LSM upgrade KIT. (**b**), (**e**), (**h**) lifetime images from three individual measurements. (**c**), (**f**), (**i**) lifetime histograms from the images b, e, and h, respectively. (**d**), (**g**), (**j**) lifetime decay fitting from the images b, e, and h, respectively. Scale bar = 10 µm.

**Figure S30.**
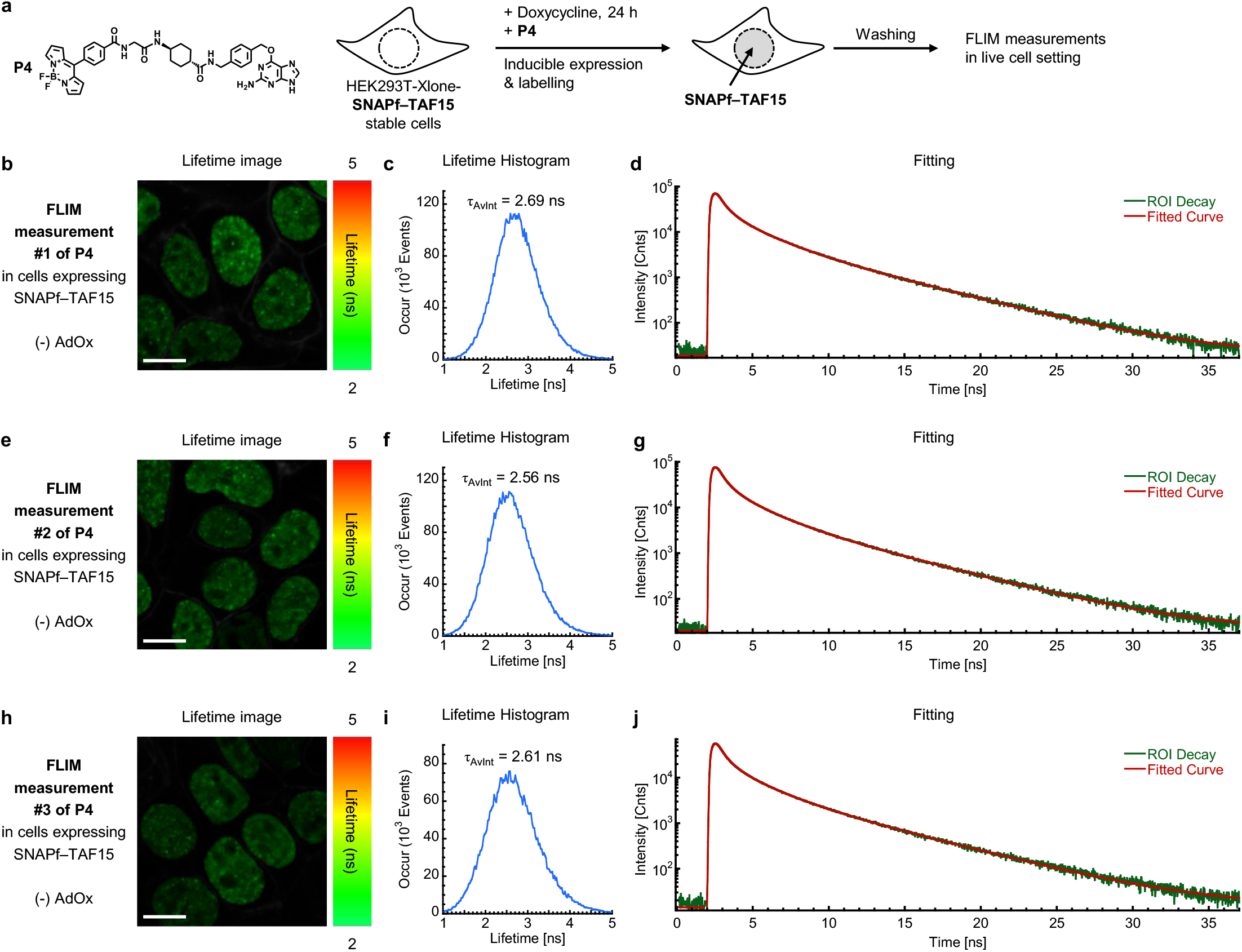
FLIM measurements of P4 with live cells stably expressing SNAPf–TAF15 without AdOx pretreatment. (**a**) The stable cells were treated with doxycycline (140 ng/mL) and **P4** (0.5 μM) for 24 hours for the inducible expression and labeling of SNAPf–TAF15, respectively. After the cells were washed with fresh fluorobrite^TM^ DMEM media supplemented with fetal bovine serum (10%) to remove excess probes, the lifetime of **P4** was measured using Zeiss LSM 880 microscope with a PicoQuant-FLIM LSM upgrade KIT. (**b**), (**e**), (**h**) lifetime images from three individual measurements. (**c**), (**f**), (**i**) lifetime histograms from the images b, e, and h, respectively. (**d**), (**g**), (**j**) lifetime decay fitting from the images b, e, and h, respectively. Scale bar = 10 µm.

**Figure S31.**
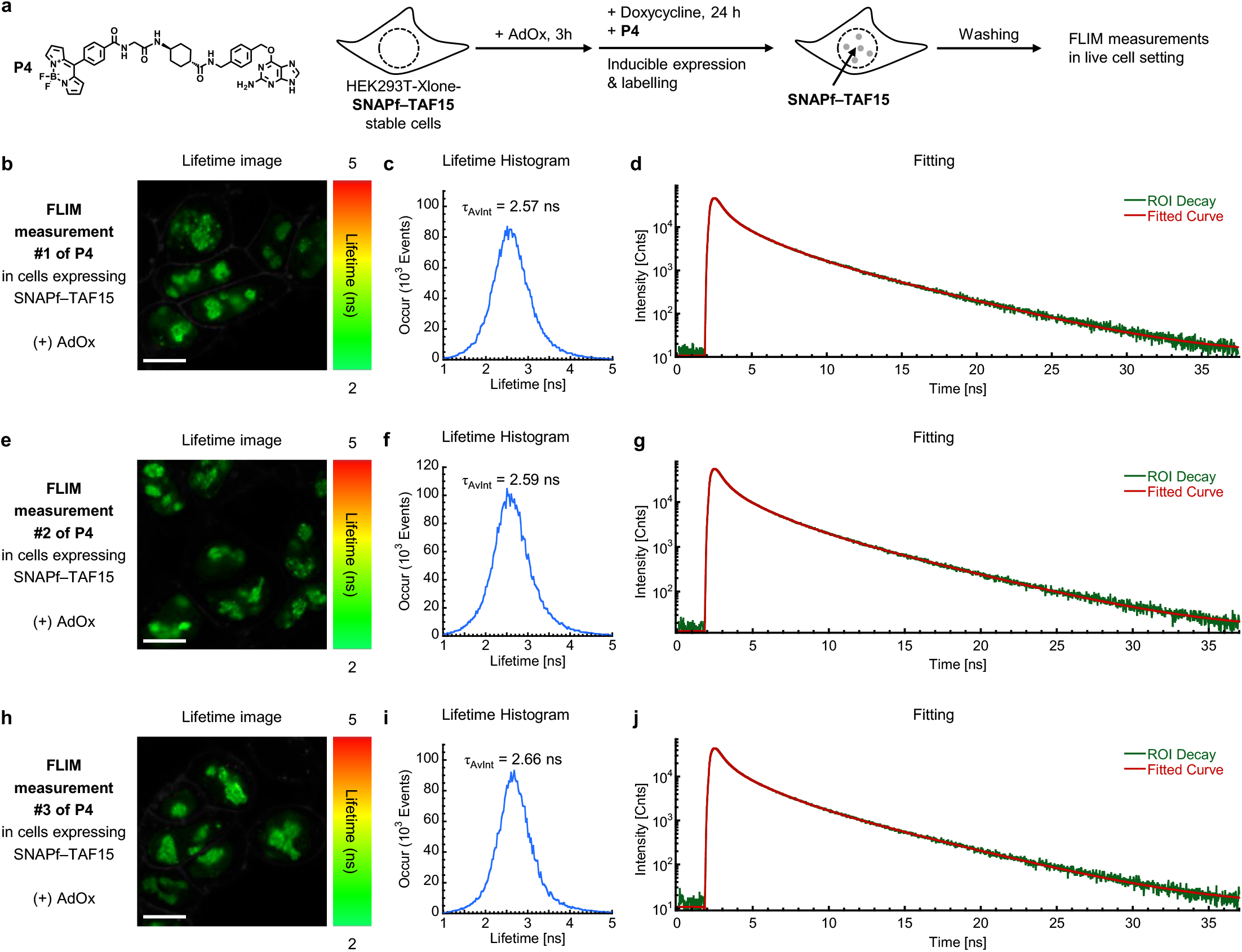
FLIM measurements of P4 with live cells stably expressing SNAPf–TAF15 with AdOx pretreatment. (**a**) The stable cells were pretreated with AdOx (25 μM) 3 hours before they were further treated with doxycycline (140 ng/mL) and **P4** (0.5 μM) for 24 hours for the inducible expression and labeling of SNAPf– TAF15, respectively. After the cells were washed with fresh fluorobrite^TM^ DMEM media supplemented with fetal bovine serum (10%) to remove excess probes, the lifetime of **P4** was measured using Zeiss LSM 880 microscope with a PicoQuant-FLIM LSM upgrade KIT. (**b**), (**e**), (**h**) lifetime images from three individual measurements. (**c**), (**f**), (**i**) lifetime histograms from the images b, e, and h, respectively. (**d**), (**g**), (**j**) lifetime decay fitting from the images b, e, and h, respectively. Scale bar = 10 µm.

## NMR Characterizations

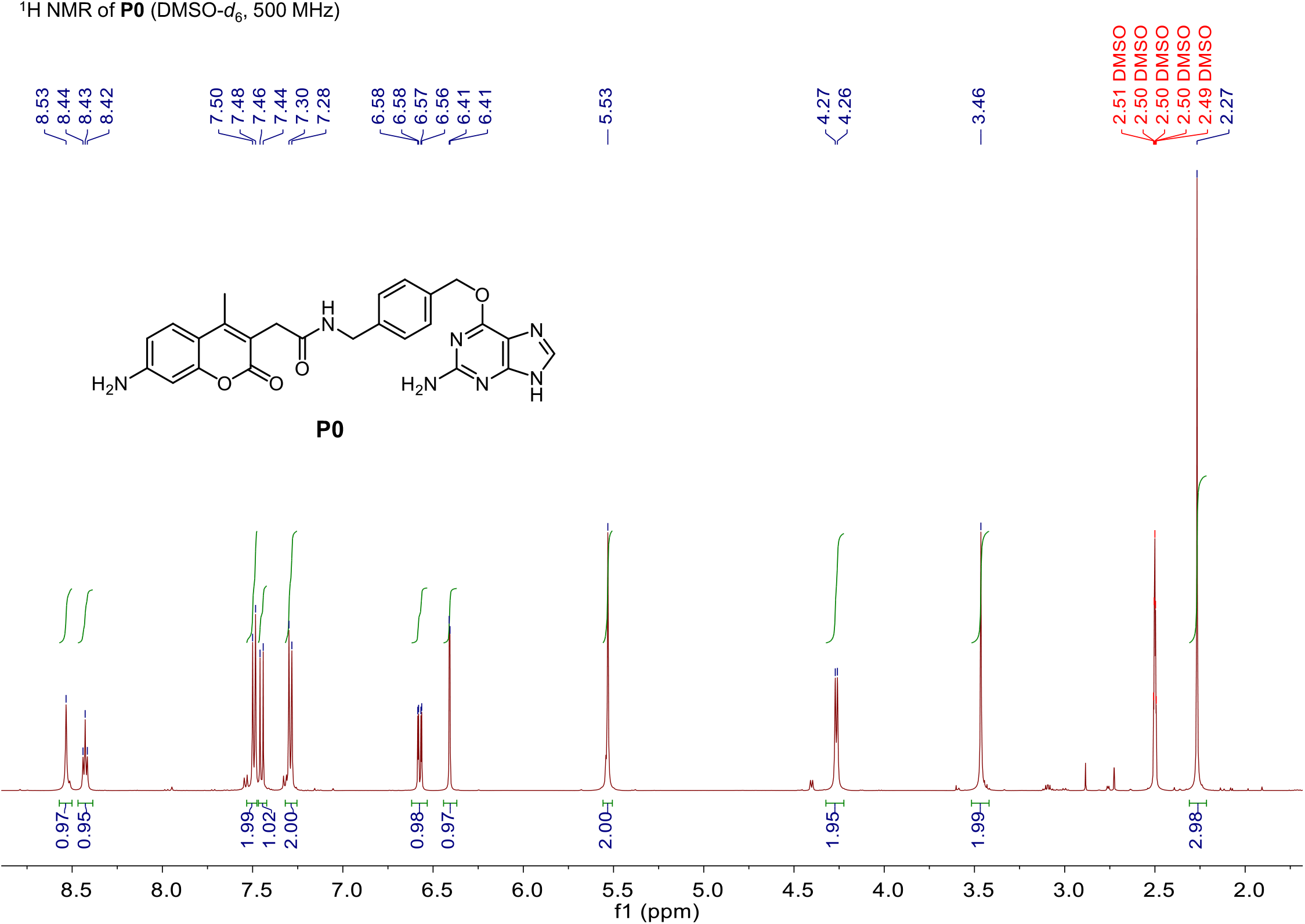

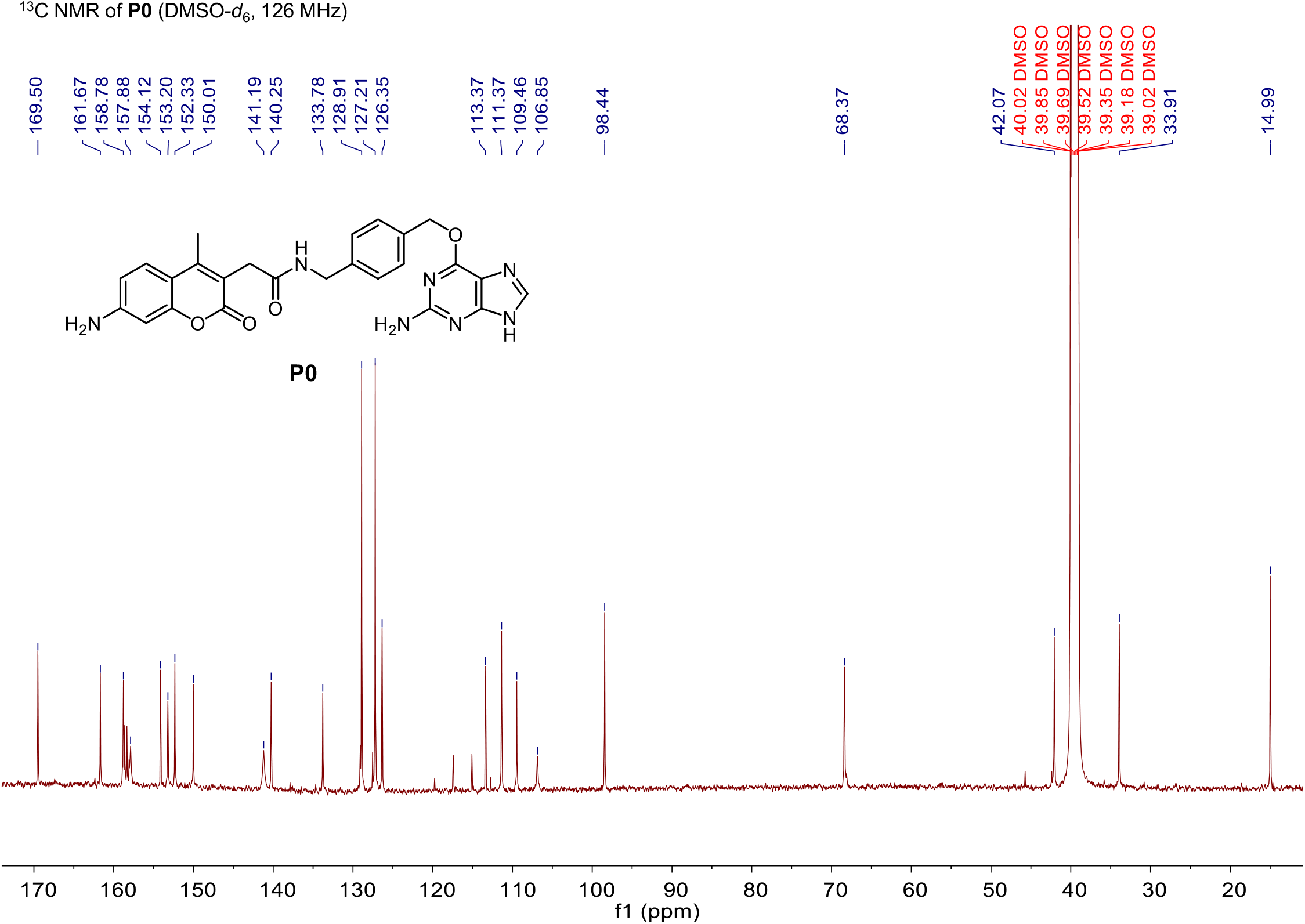

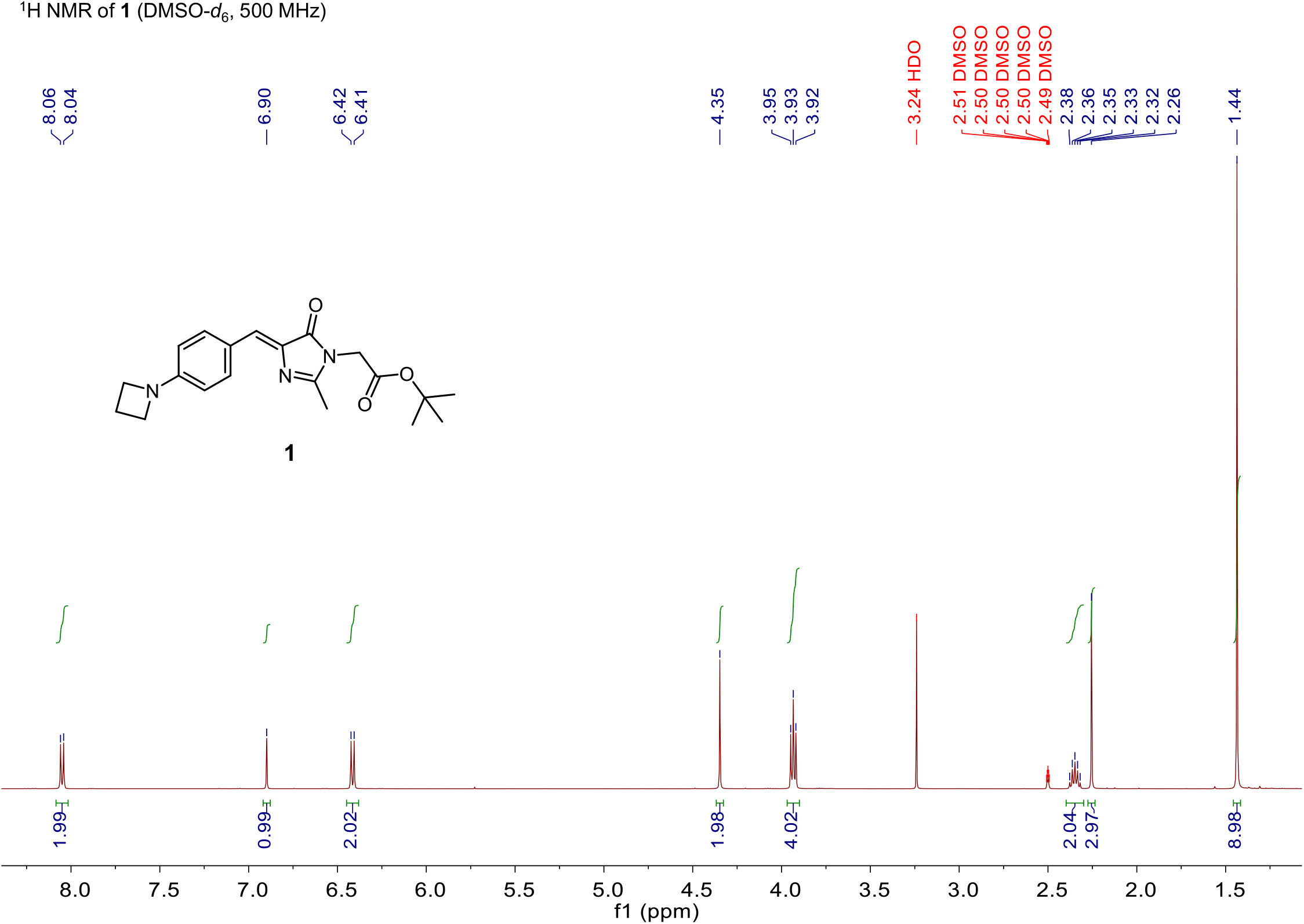

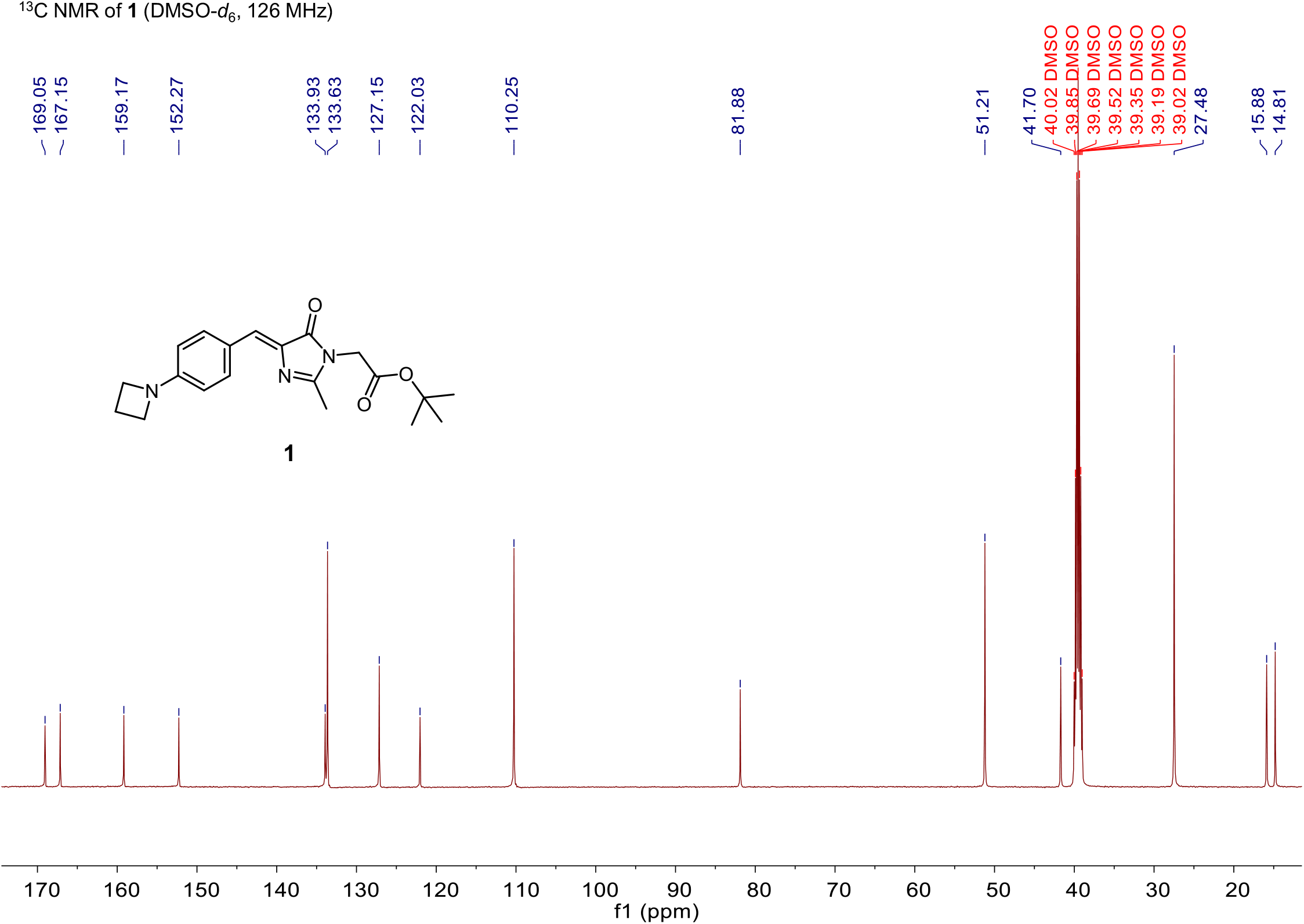

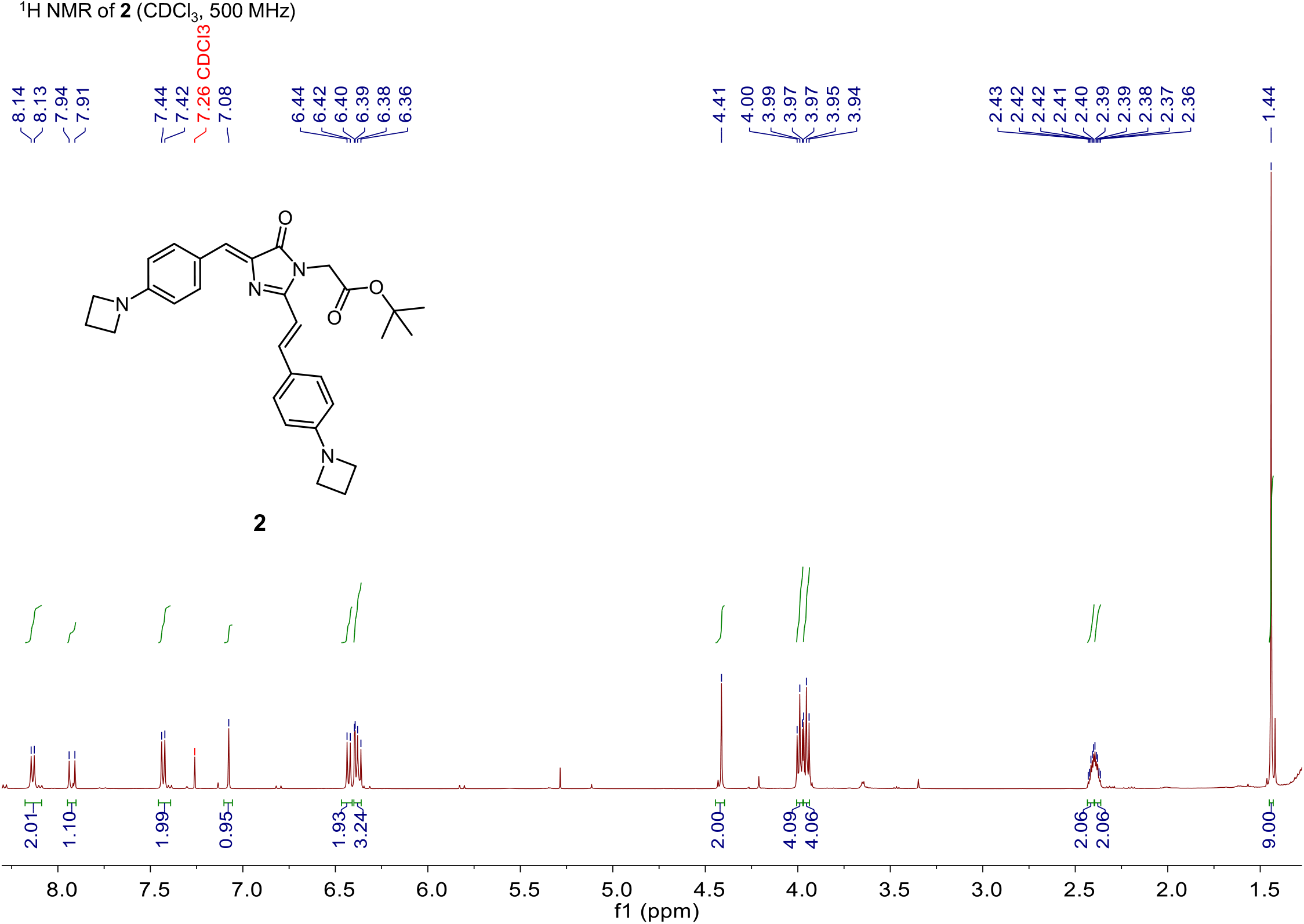

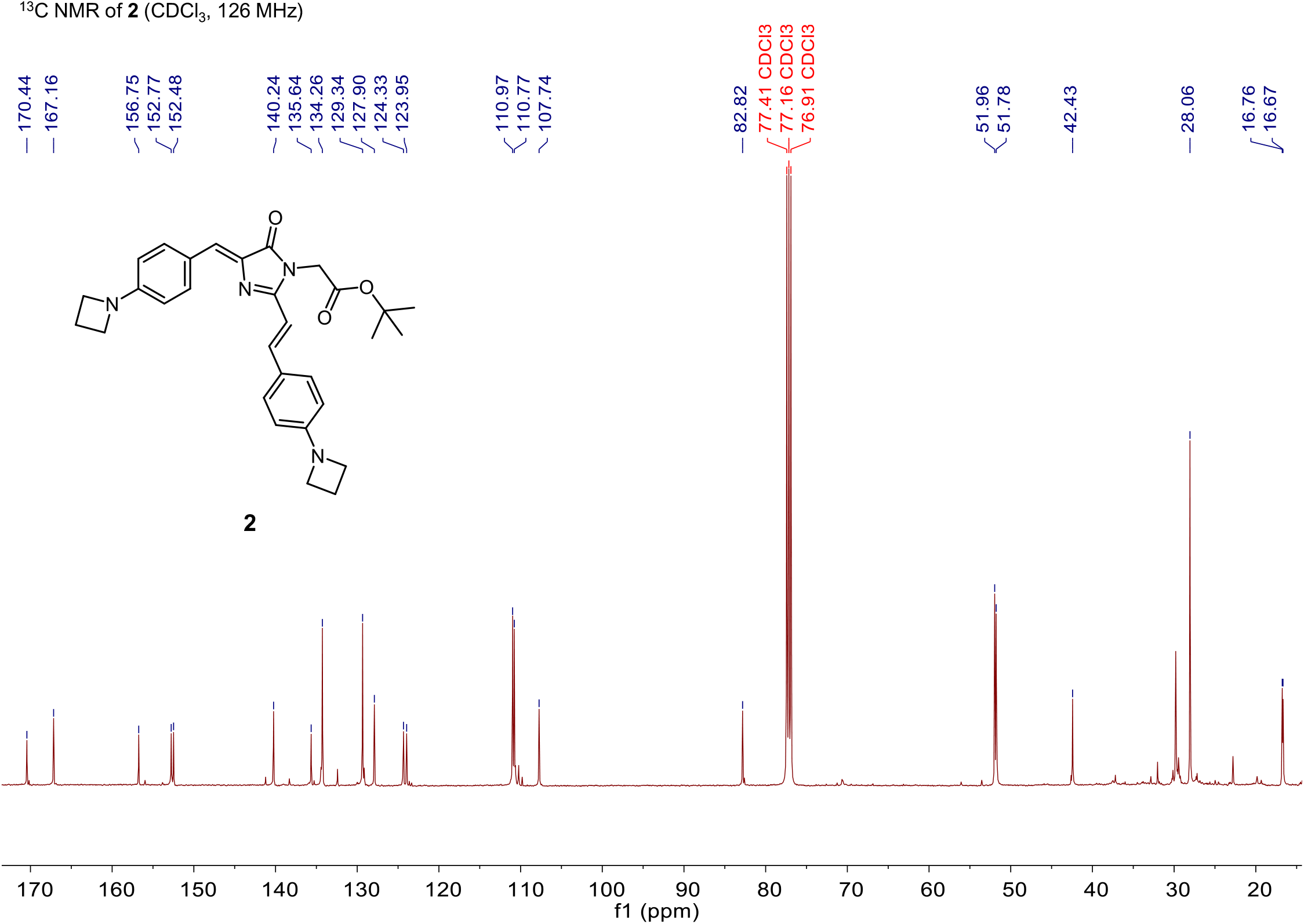

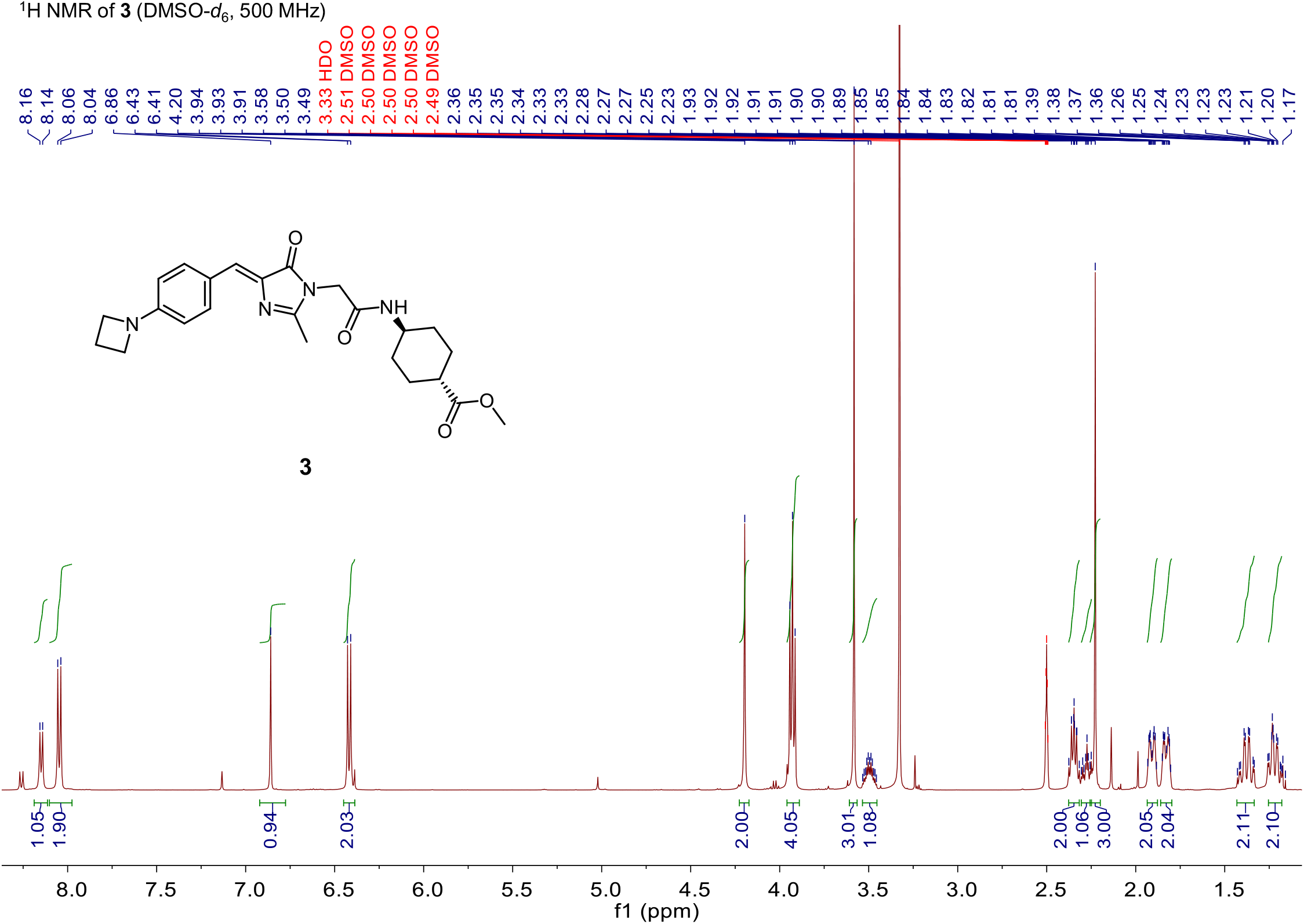

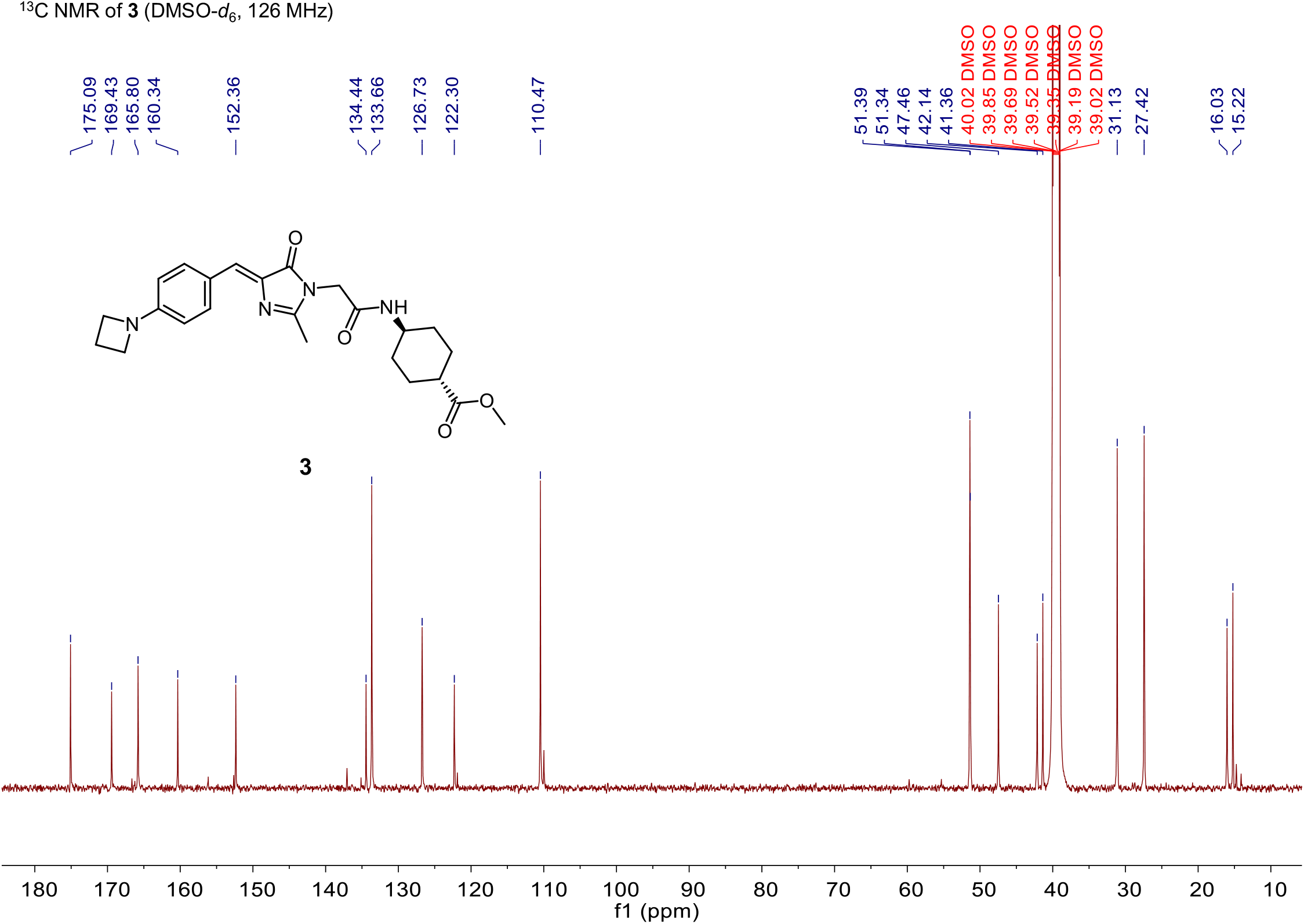

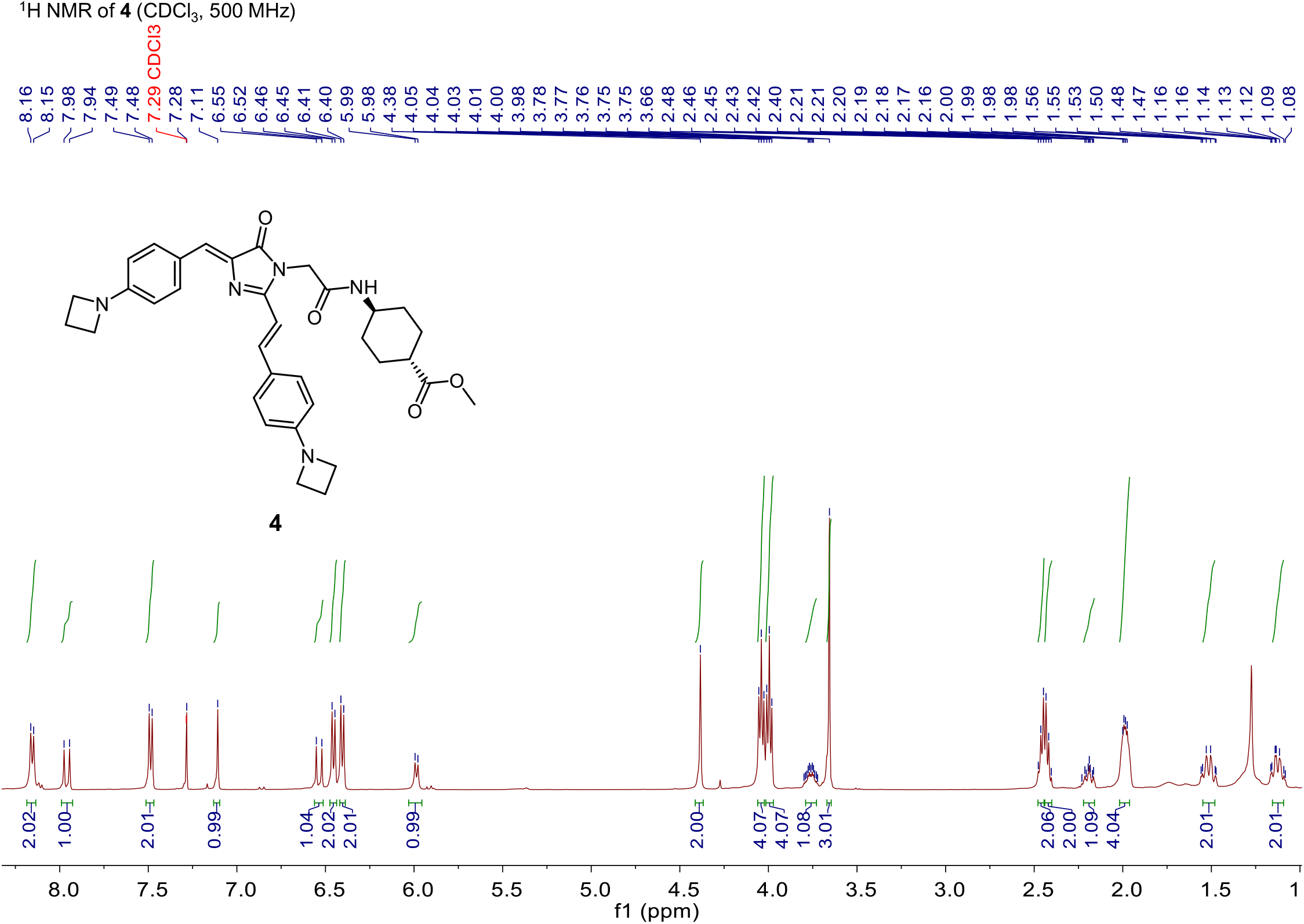

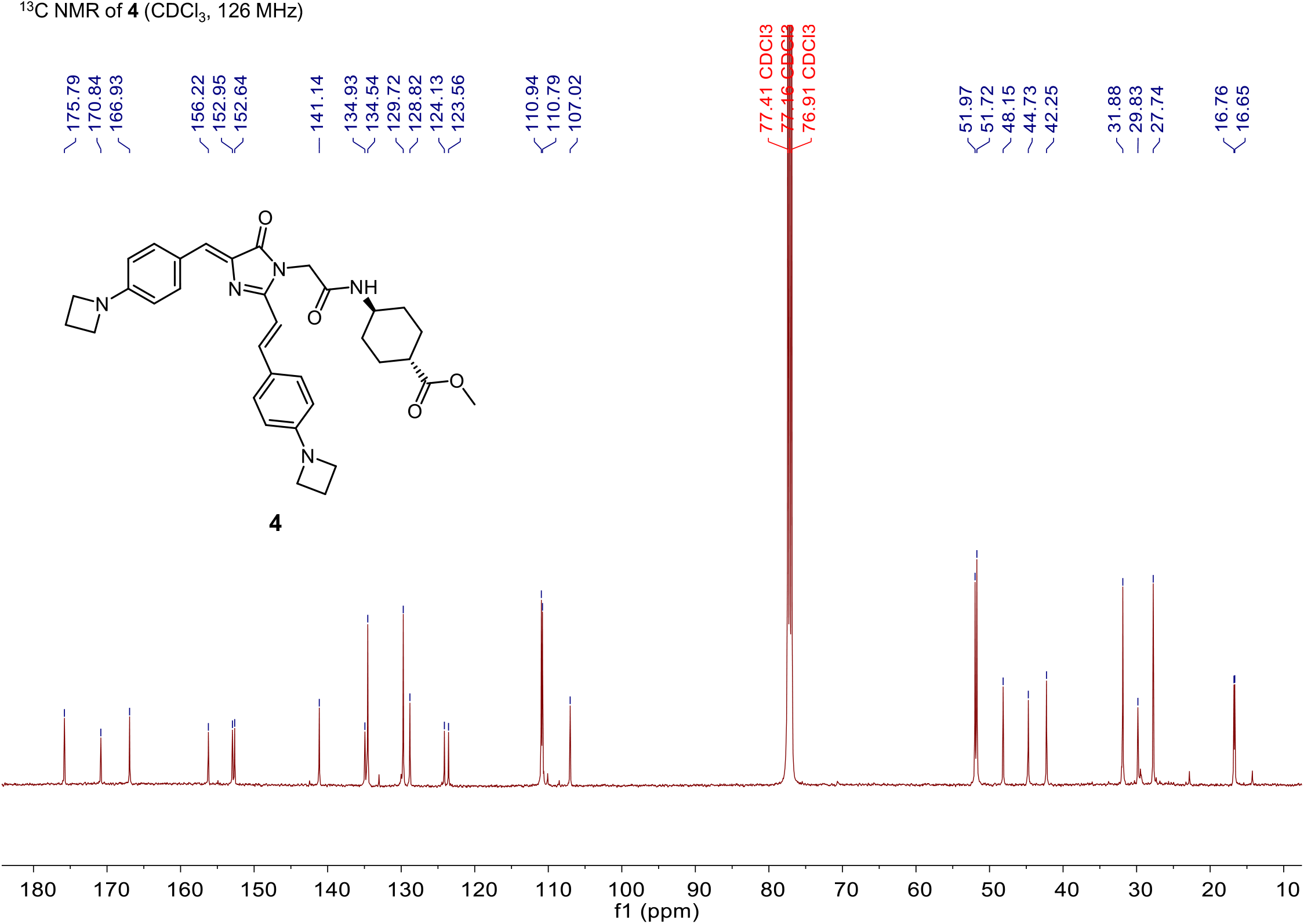

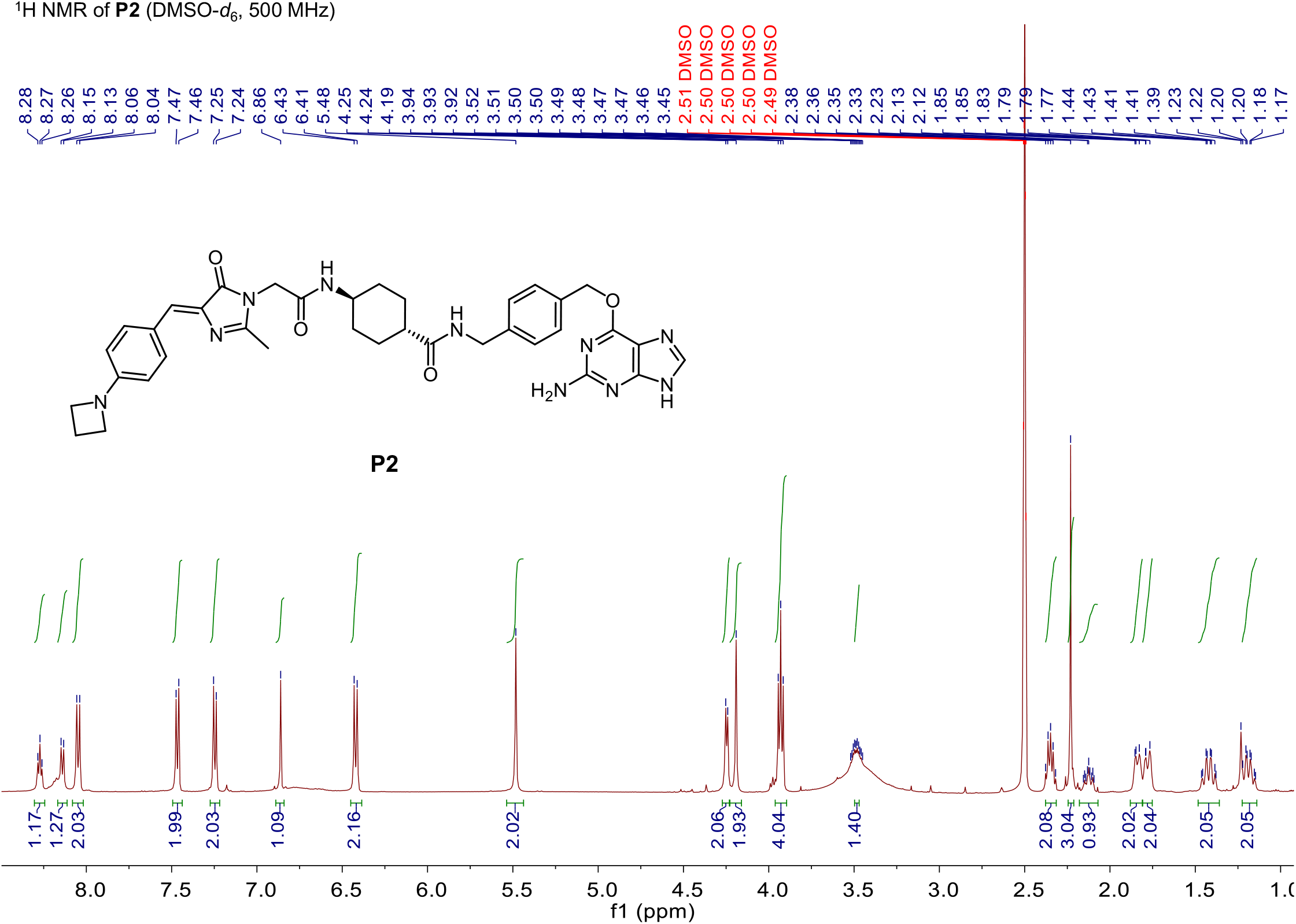

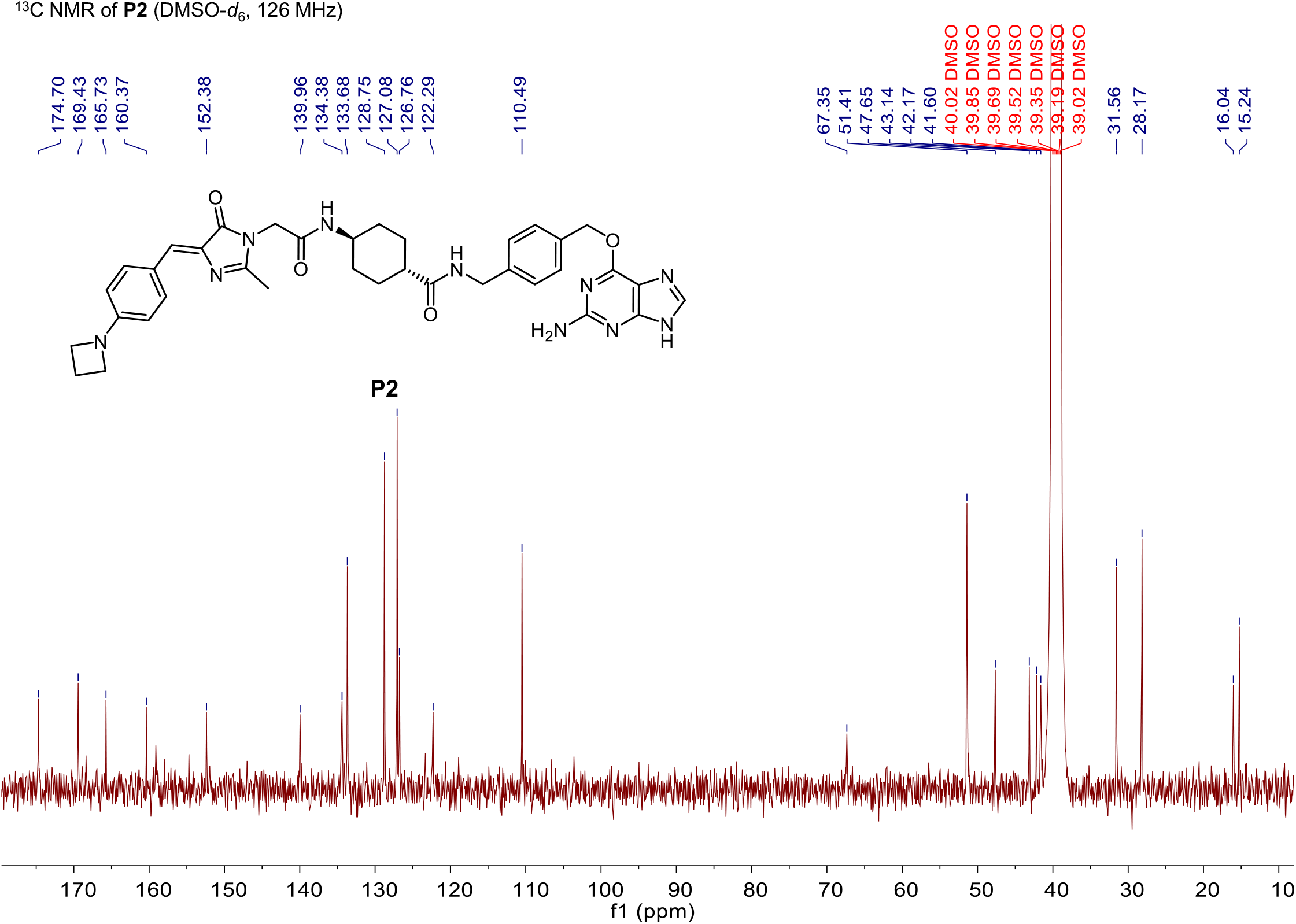

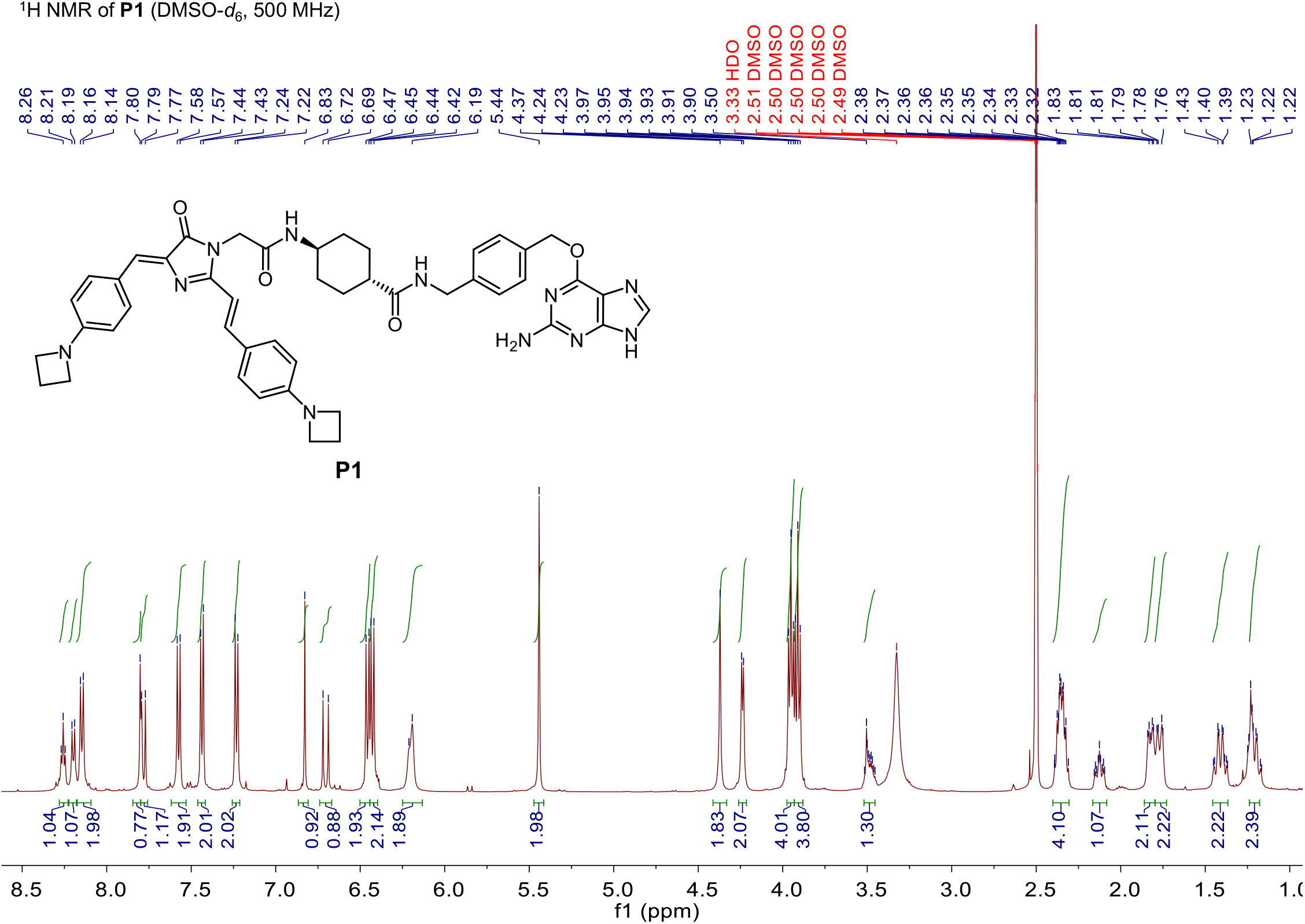

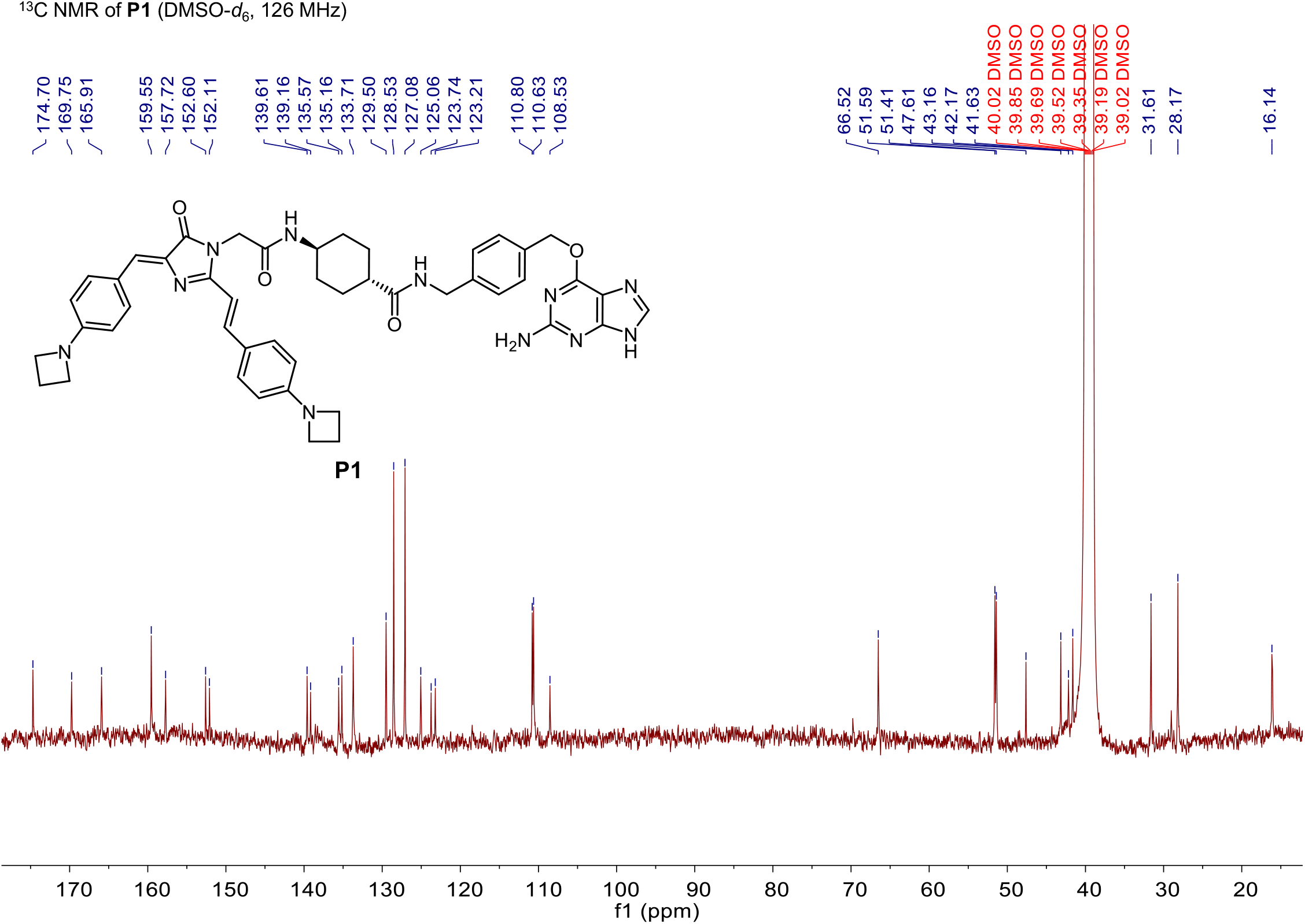

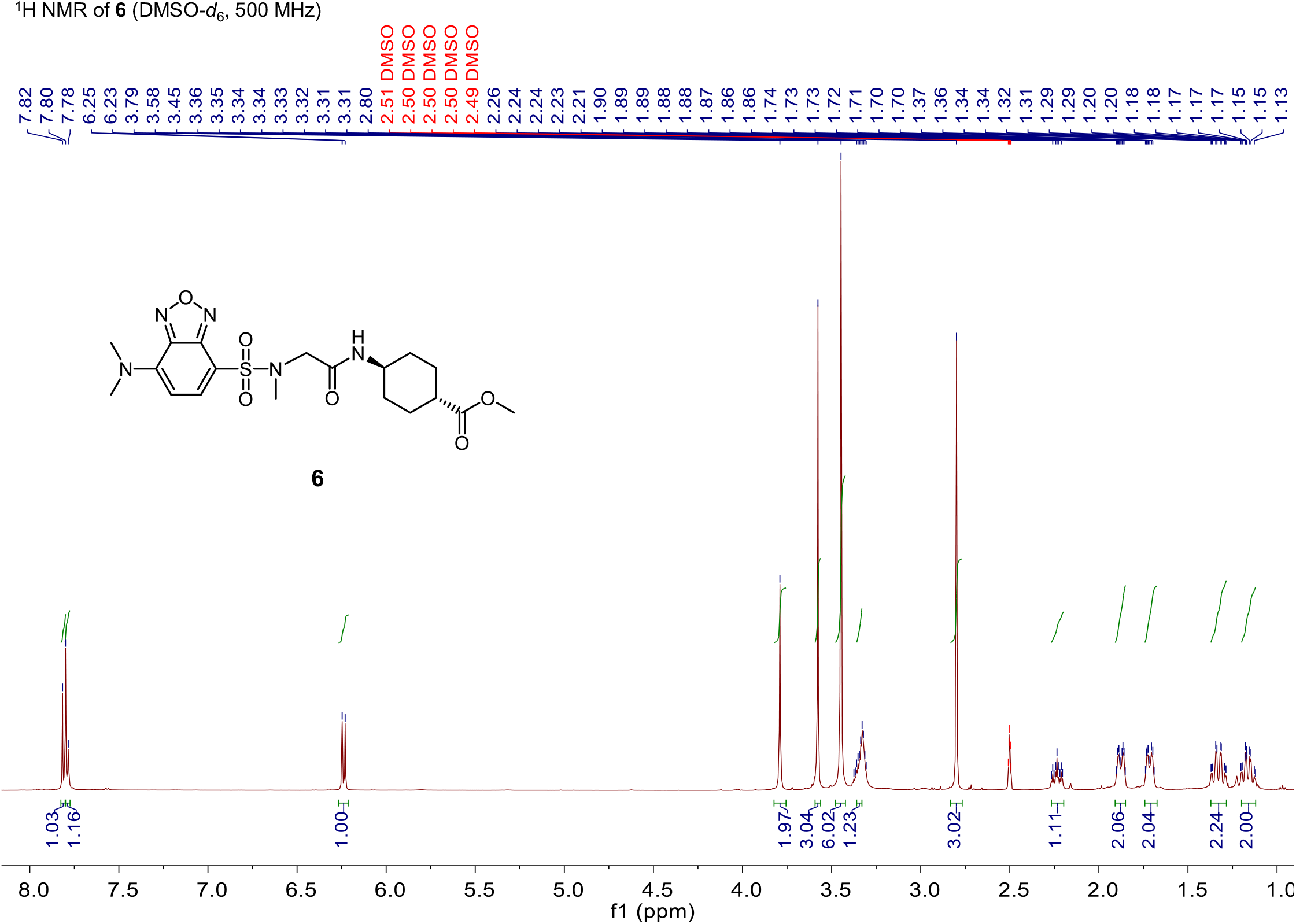

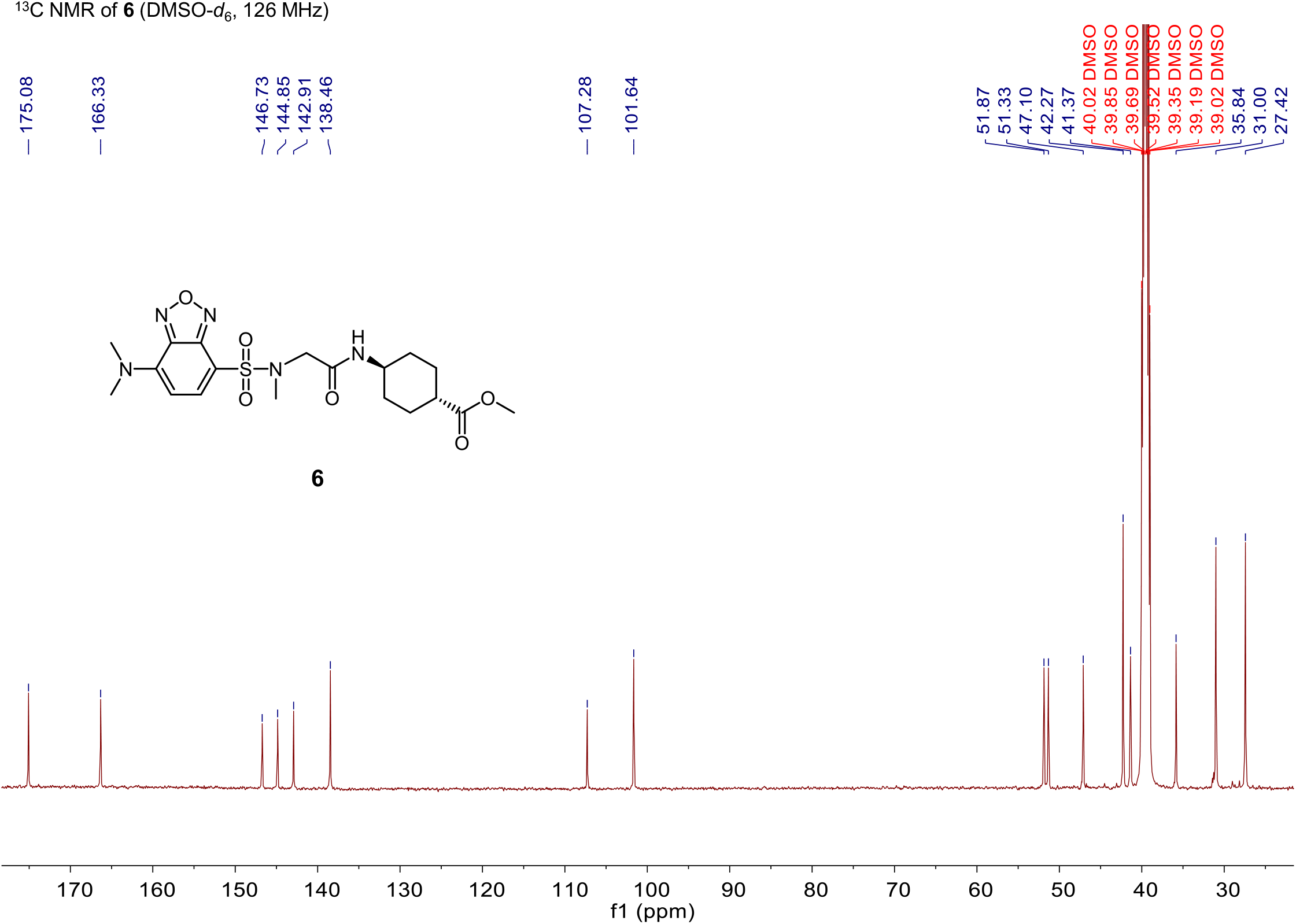

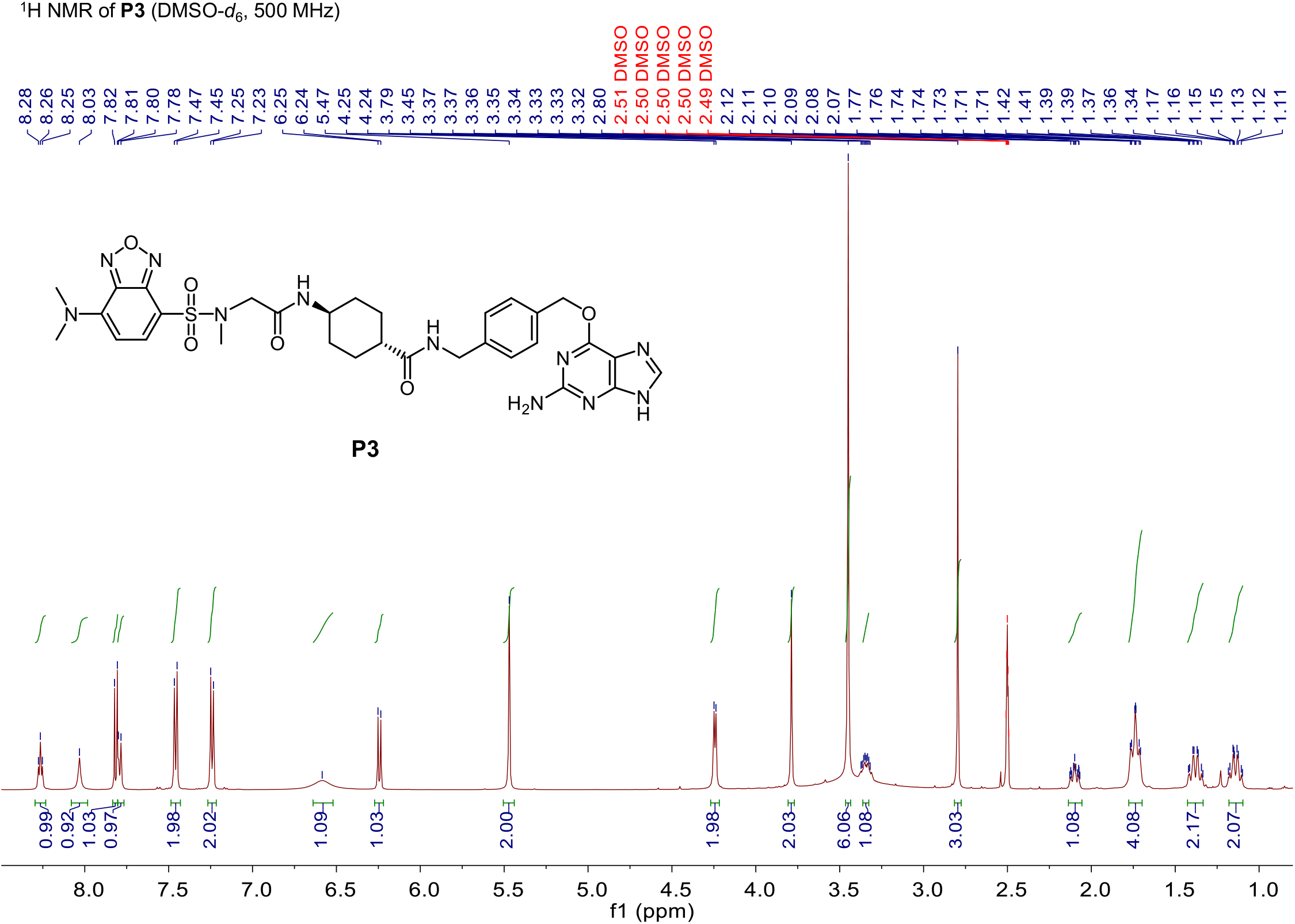

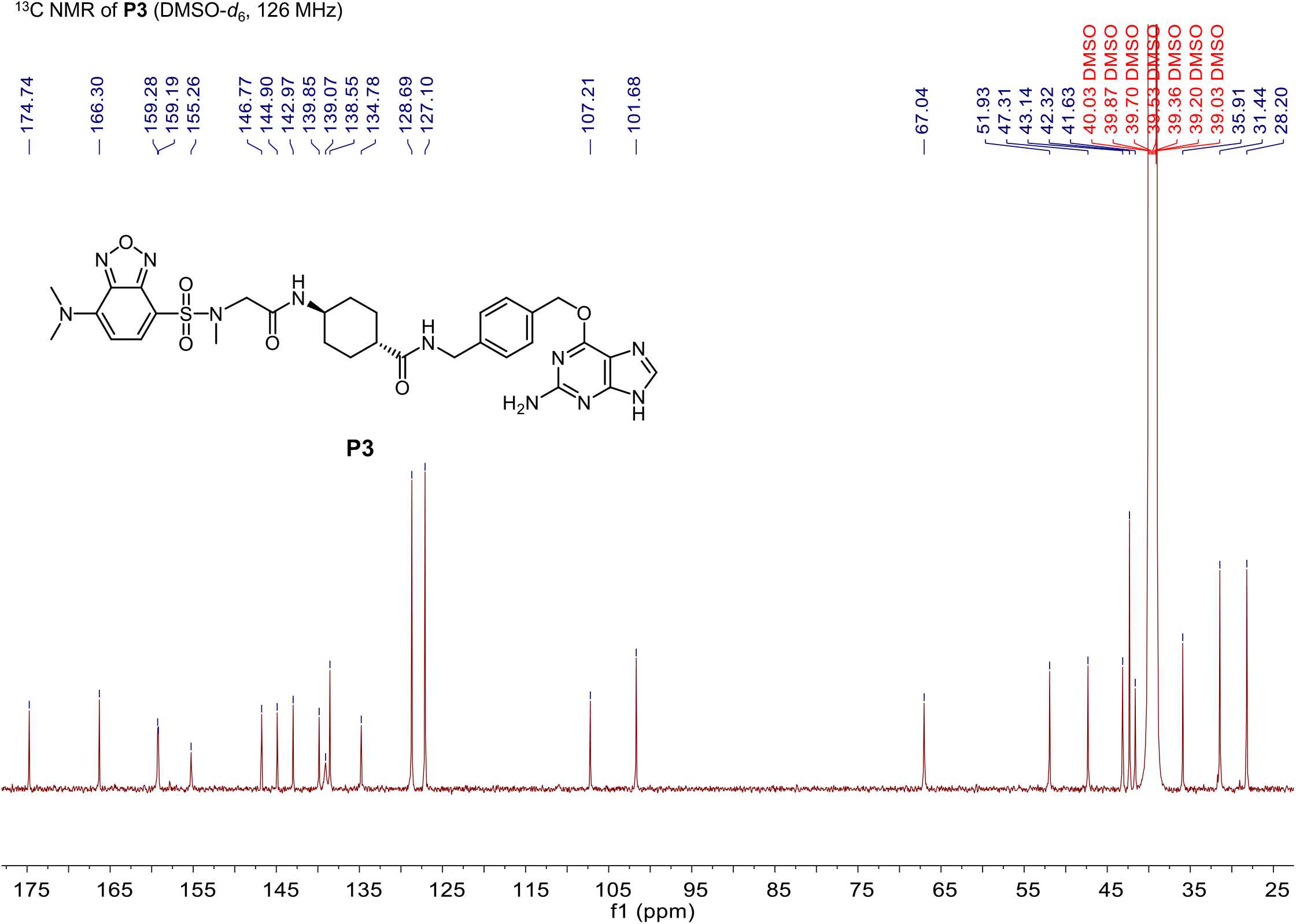

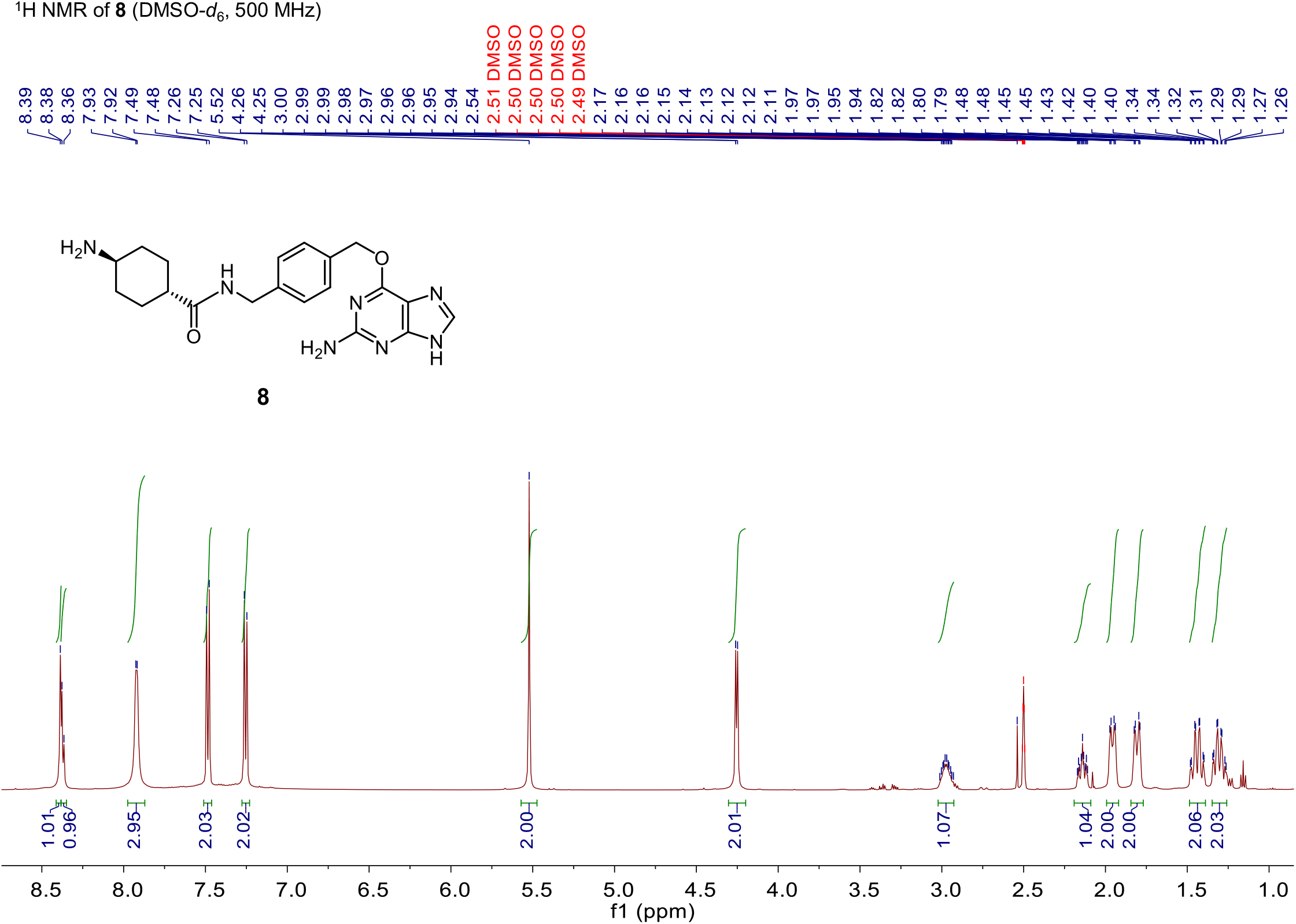

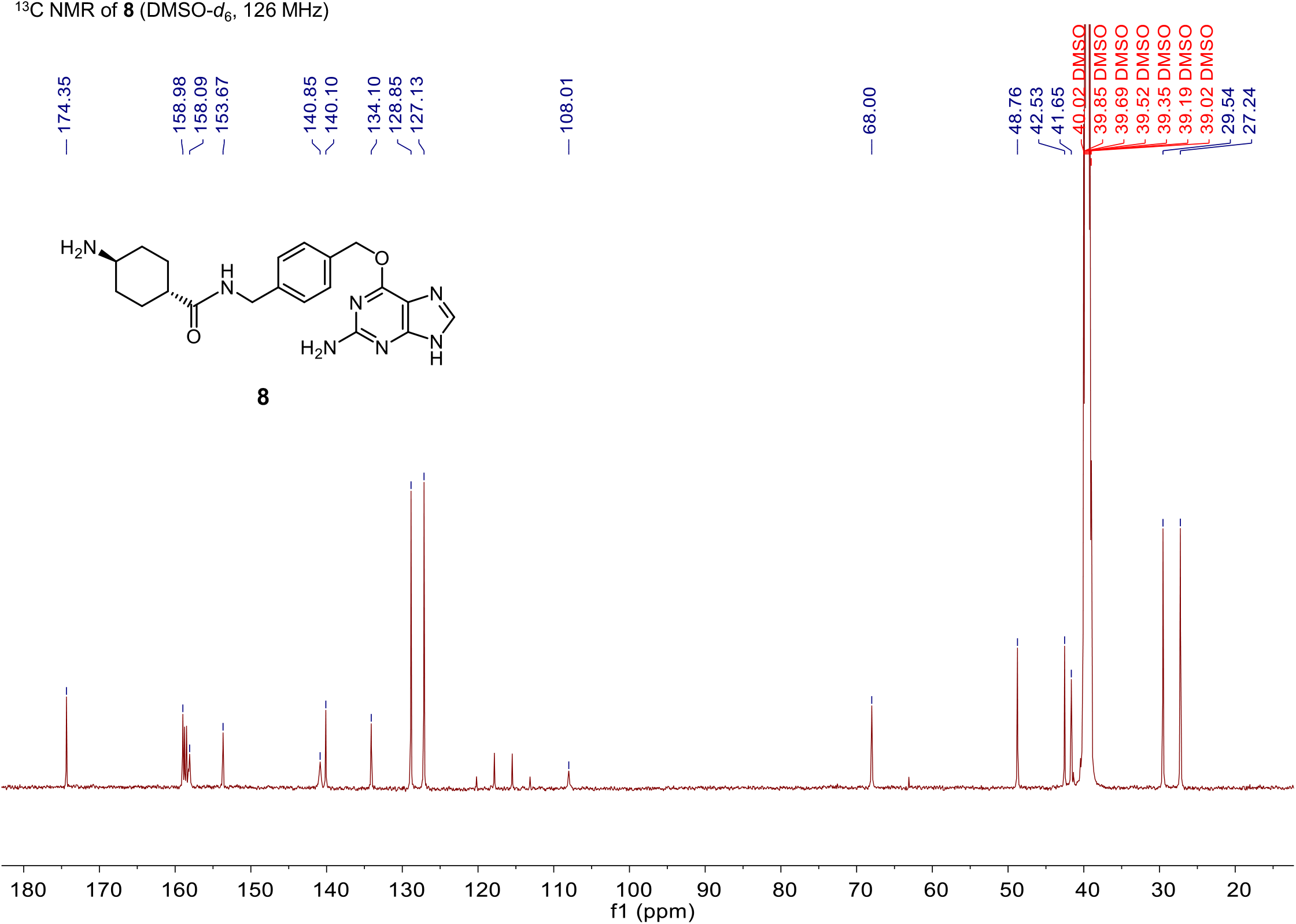

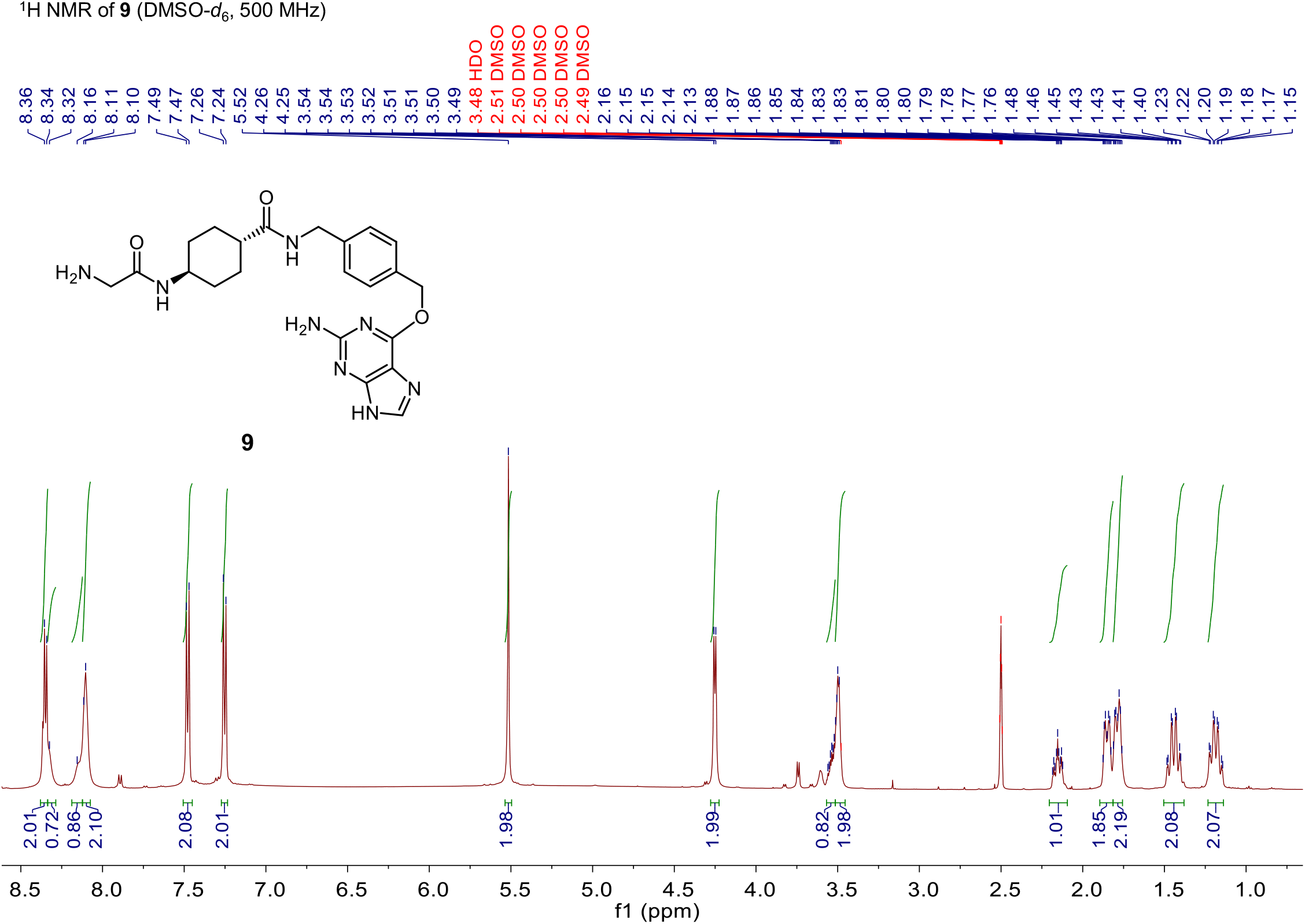

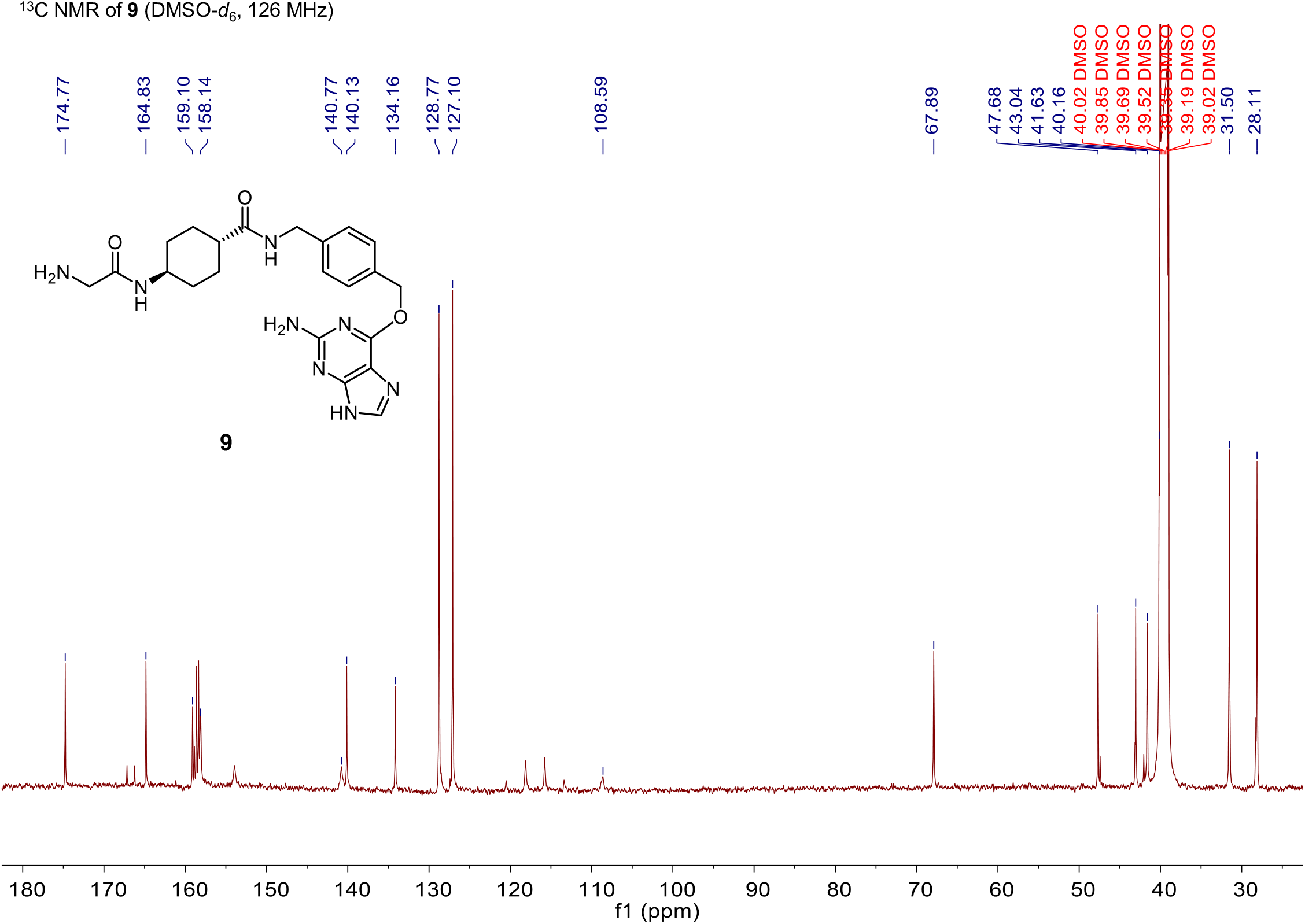

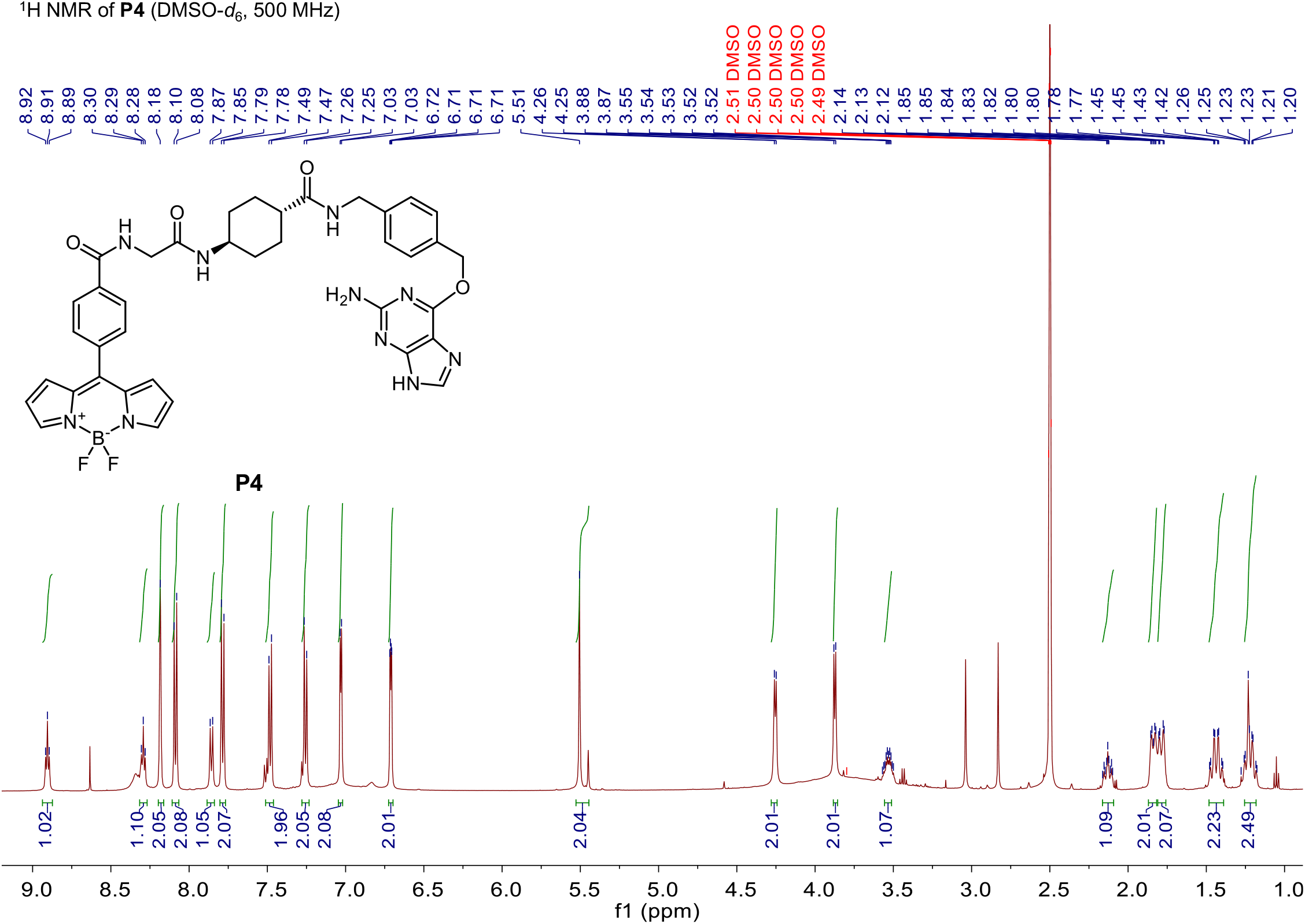

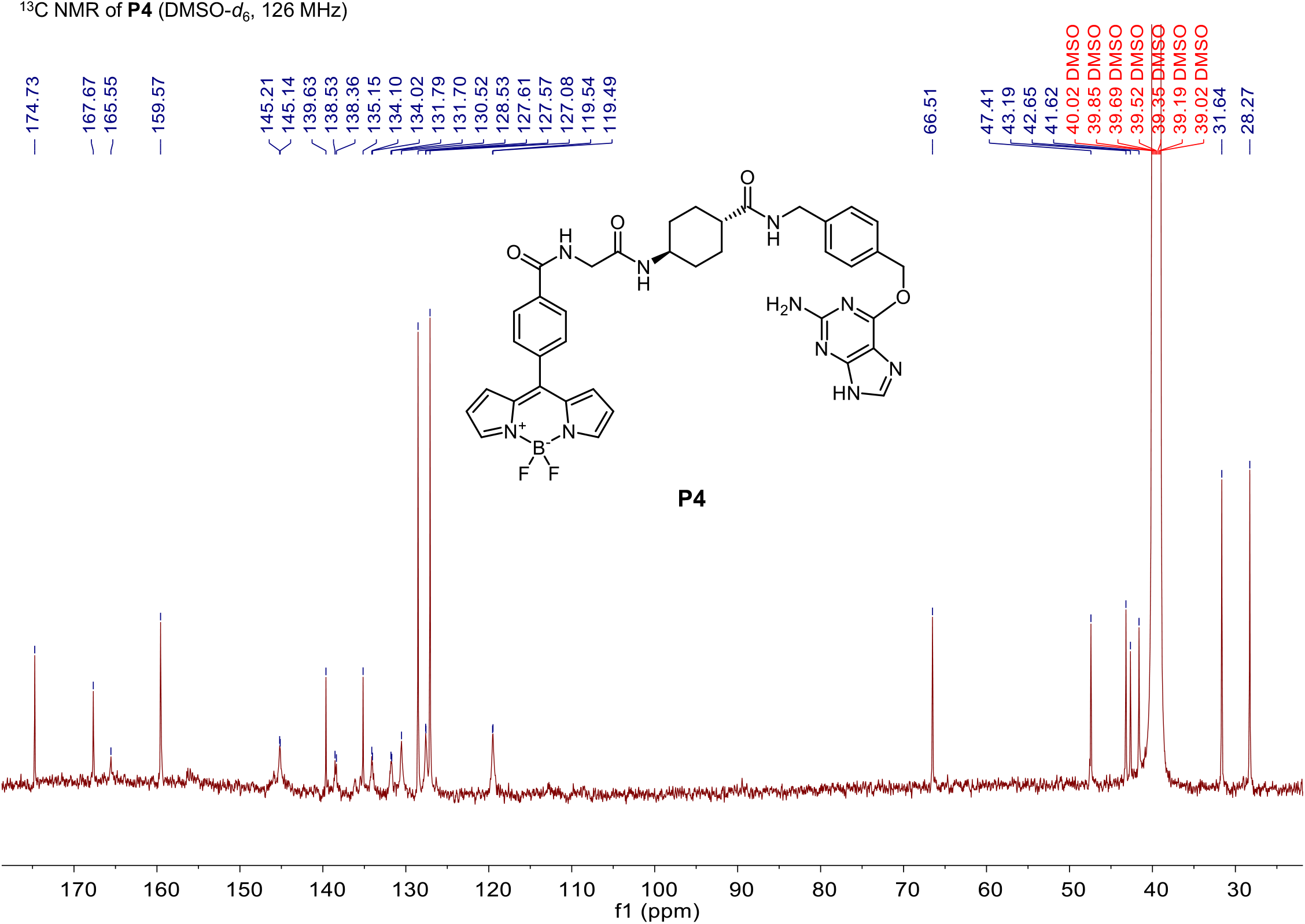

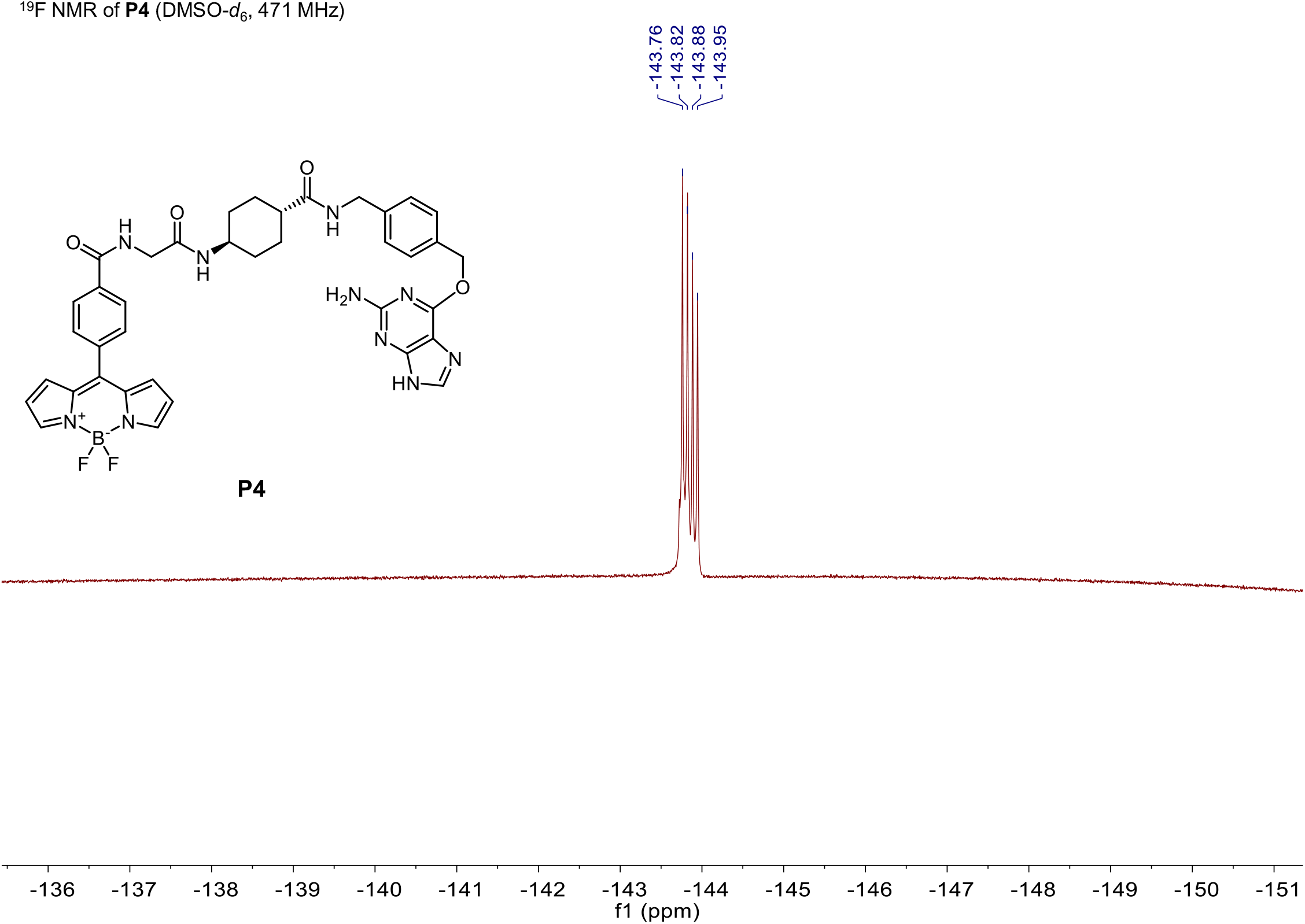

## Notes

### Competing Interest Statement

The authors have declared no competing interest.

